# A quantitative and spatial analysis of cell cycle regulators during the fission yeast cycle

**DOI:** 10.1101/2022.04.13.488127

**Authors:** Scott Curran, Gautam Dey, Paul Rees, Paul Nurse

## Abstract

We have carried out a systems-level analysis of the spatial and temporal dynamics of cell cycle regulators in the fission yeast *Schizosaccharomyces pombe*. In a comprehensive single cell analysis we have precisely quantified the levels of 38 proteins previously identified as regulators of the G2 to mitosis transition, and of 7 proteins acting at the G1 to S-phase transition. Only two of the 38 mitotic regulators exhibit changes in concentration at the whole cell level, the mitotic B-type cyclin Cdc13 which accumulates continually throughout the cell cycle, and the regulatory phosphatase Cdc25 which exhibits a complex cell cycle pattern. Both proteins show similar patterns of change within the nucleus as in the whole cell but at higher concentrations. In addition, the concentrations of the major fission yeast cyclin dependent kinase (CDK) Cdc2, the CDK regulator Suc1 and the inhibitory kinase Wee1 also increase in the nucleus peaking at mitotic onset but are constant in the whole cell. The significant increase in concentration with size for Cdc13 supports the model that mitotic B-type cyclin accumulation acts as a cell size sensor. We propose a two-step process for the control of mitosis. First, Cdc13 accumulates in a size-dependent manner which drives increasing CDK activity. Second, from mid G2 the increasing nuclear accumulation of Cdc25 and the counteracting Wee1 introduces a bistability switch that results in a rapid rise of CDK activity at the end of G2 and thus brings about an orderly progression into mitosis.

**Significance Statement:** Across eukaryotes the increasing level of cyclin dependent kinase (CDK) activity drives progression through the cell cycle. As most cells divide at specific sizes, information responding to the size of the cell must feed into the regulation of CDK activity. In this study, we use fission yeast to precisely measure how proteins that have been previously identified in genome wide screens as cell cycle regulators change in their levels with cell cycle progression. We identify the mitotic B-type cyclin Cdc13 and mitotic inhibitory phosphatase Cdc25 as the only two proteins that change in both whole cell and nuclear concentration through the cell cycle, making them candidates for universal cell size sensors at the onset of mitosis and cell division.

## Introduction

Steady state growing eukaryotic cells generally coordinate their cell cycles with cell growth by ensuring that mitosis and the associated cell division takes place when a particular cell size is attained (1–3). The mechanisms that bring about mitotic onset are known to be accurate because cell size at mitosis exhibits little variation, and also efficiently homeostatic because perturbation from a mean population size at mitosis is corrected within one to two subsequent cycles (4, 5). These control mechanisms are also integrated with cell ploidy as cells roughly double their size at mitosis with each doubling of DNA content (6–9). Given the conservation of genes involved in cell cycle control from yeasts to mammalian cells (10) and that co-ordination of mitosis and cell division with cell size is observed across eukaryotes, the molecular mechanisms involved are likely to share commonalties. A number of models for monitoring cell size have been proposed (11–17), with one of the most straightforward being for changes in the concentration of a mitotic regulatory component to accompany cell size increase until a threshold level is reached that allows mitosis to proceed (18). This may be achieved either by an increase in the concentration of a mitotic activator, or by a decrease in the concentration of an inhibitor (11, 12). Such cell size sensing mechanisms could be integrated with the monitoring of ploidy by interactions such as titration of activators or inhibitors onto the DNA or chromatin (19). These sensing mechanisms must also be coupled to the activation of the cyclin dependent kinase (CDK) which brings about mitosis followed by cell division in all eukaryotes (20). The dynamics of CDK activation must be such that there is a sharp and irreversible entry into mitosis which could be influenced by the molecular mechanisms sensing cell size. In this paper we investigate these mechanisms by measuring the levels of mitotic regulators during the cell cycle, both in the whole cell and in the nucleus where critical mitotic events occur. These studies of the levels of mitotic regulators in single cells are aimed at informing our understanding of how cells sense their size and regulate the dynamics of CDK activation during the cell cycle and at the onset of mitosis.

The fission yeast *Schizosaccharomyces pombe* is an ideal model organism for investigating the coordination of mitosis with cell size as 99% of genes (5059 genes) have previously been deleted (21) and systematically screened for cell cycle phenotypes (22). This screen uncovered a total of 513 cell cycle genes. Two further screens of this collection revealed genes that specifically regulate mitotic signalling. One exploited heterozygous deletions of diploid cells where gene expression levels are reduced to half of normal to identify haploinsufficient genes, whereby cells are either advanced or delayed into mitosis at a smaller or larger size, respectively. This screen identified 17 genes (23). A second screen covering 82% of viable haploid deletions (totaling 2969 genes), identified 18 genes that were advanced into the G2 to mitosis transition at a smaller size (24). These genes encode proteins which are rate limiting for entry into mitosis, three of which overlapped with the haploinsufficiency study. Together these two screens identified 32 genes encoding potential mitotic regulatory molecules that also serve as candidates for cell size sensing. This gene set includes those encoding proteins at the core of the CDK cell cycle control network, such as the main CDK in fission yeast Cdc2, the G2/mitosis B-type cyclin Cdc13, the activating phosphatase Cdc25, and the inhibitory protein kinase Wee1 (25–28).

A further reason why fission yeast lends itself well to the study of cell cycle regulation is because their simple rod shaped geometry and growth by tip extension allows for cell cycle position to be determined by cell length (29). Changes in the levels of molecules during the cell cycle can therefore be determined in asynchronous, steady state growing cultures by measuring the levels of endogenously-tagged fluorescent proteins in single cells. Such a method avoids any perturbations induced by synchronising procedures, which can disturb measurements of protein levels. Using two independent single cell approaches, we have accurately measured the precise concentrations and absolute levels of these potential mitotic regulators in the whole cell and nucleus for up to hundreds of thousands of cells at single-cell resolution throughout the cell cycle. This study has identified the subset of mitotic regulators that exhibit changes in their whole cell or nuclear concentration, making them prime candidates for cell size or ploidy sensors whose readout tunes the dynamics of CDK activity. These proteins are conserved across eukaryotes and so may therefore be of universal relevance.

## Results

Progression through the fission yeast cell cycle is concomitant with growth of the cell by tip extension allowing for cell length to be used as an indicator of cell cycle position (Fig. 1A). Fission yeast cells have a very short G1 which occurs after mitotic exit. Binucleate and septated cells contain two 1C nuclei which undergo S-phase as cell division takes place. Thus, taking account of cell length and whether cells are mononucleate, binucleate, or binucleate and septated, allows the specification of the fission yeast cell cycle. To assess the changes in the levels of mitotic regulatory molecules during the cell cycle, we endogenously tagged 30 proteins encoded by genes identified in the previously described haploinsufficiency and rate limiting screens with single colour fluorescent labels in individual wild type fission yeast strains (The 30 excludes *nsp1* and *nup186* from the original 32 because these 2 showed no measureable fluorescence when tagged). We also tagged 8 more genes which could be involved in mitotic control including *cdr2, plo1* and *pyp3*, and for comparative purposes 9 further genes acting at the G1 to S-phase transition (*cdc10, cdc18, cdc20, cig1, cig2, mik1, puc1, rum1* and *srw1*). All genes were tagged with mNeonGreen, chosen for its fast maturation time and bright signal (30), other than *wee1* (N-terminal GFP) (31), *cdc13* (internal superfolder GFP) (32) and *pka1* (C-terminal GFP) (Table S1 for the full list of tagged proteins). To check that tagging of these genes did not have a major effect on cell size at mitotic onset, the length at septation was measured for all of the strains (Fig. S1). There were negligible effects for the majority, the exceptions being the tagging of *wee1* and *pyp1* which induced elongation and *cdr1*and *ppa2* which induced shortening (Fig. S1A & SC). This suggests there is some loss of function for Cdr1, Ppa2 and Pyp1 and perhaps stabilisation of Wee1, due to the tagging.

**Figure 1.**
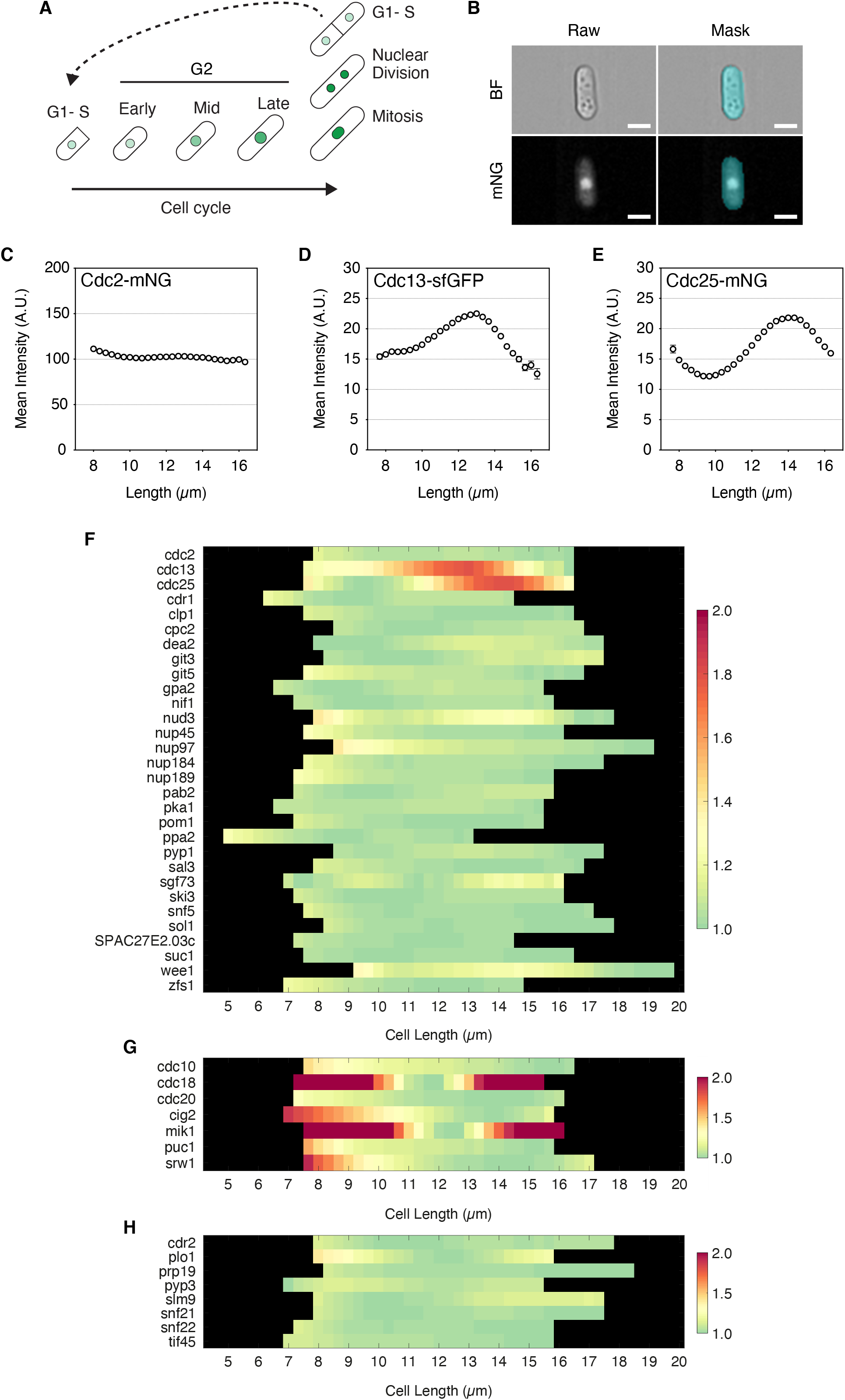
Only Cdc13 and Cdc25 increase in whole cell concentration to a peak at mitotic entry. **A**, Schematic of the fission yeast cell cycle. Cells increase in length with progression through G2. Green indicates a representative protein that increases in nuclear concentration with cell cyle progression and increasing cell length that decreases in nuclear concentration after nuclear division. Cell growth slows as cells enter mitosis and undergo nuclear division. G1 and S-phase occur in binucleates as cells septate. For *YeaZ* cell segmentation binucleate cells are split at the point of septation, so cells in G1-S are the shortest cells in the plots for Figures 2-4. **B**, Representative images of a mononucleate Cdc2-mNG cell imaged by imaging flow cytometry. Top left, Brightfield; Bottom left, mNG. Top right, Segmentation mask of BF image. Bottom right, Overlay of mask onto fluorescent image. Scale bar, 5 μm. **C**-**E**, Plots of imaging flow cytometry data for Cdc2-mNG (**C**), Cdc13-sfGFP (**D**) and Cdc25-mNG (**E**) populations showing mean whole cell fluorescence intensity against cell length. Circles indicate mean, Error bars = 95% CI. Length bins = 0.33 μm, >500 cells/bin. n =**C**, 183,435 cells; **D**, 171,425 cells; and **E**, 178,598 cells. **F**, Heatmap showing the mean cellular fluorescence intensity for asynchronous populations of strains fluorescently tagged for mitotic regulators. Mean intensity for each 0.33 μm length bin is normalised to each strain’s minimum and plotted against cell length. >500 cells/bin. All strains are endogenously tagged with mNG, except for Cdc13 (internal sfGFP), Wee1 (N-terminal GFP) and Pka1 (C-terminal GFP). **G**, as **F** except for G1 to S-phase transition genes, and **H**, for other genes of interest involved in mitotic control.

Each strain was individually imaged using imaging flow cytometry (Amnis Imagestream X) from exponentially growing asynchronous cell cultures. This allowed us to image >100,000 cells per strain in each experiment giving high cell cycle coverage. Brightfield segmentation masks were overlaid onto fluorescence images (Fig. 1B) to allow for cell intensity measurements (Fig. 1C-H). For each strain, we plotted the mean fluorescence intensity relative to its own minimum against cell length to give an indication of the fold-change of protein level across the cell cycle (Fig. 1F-H). We show the data for three example strains (Fig. 1C-E), one that is constant in whole cell concentration through the cell cycle, Cdc2-mNG (Fig. 1C), and two that show changes, Cdc13-sfGFP (Fig. 1D) and Cdc25-mNG (Fig. 1E). As expected for a mitotic B-type cyclin, Cdc13 accumulates as cell size increases and falls at the end of the cell cycle. Cdc25 showed an unexpected pattern of change by oscillating in level through the cell cycle. Its concentration fell with cell size increase in the first third of the cycle, before reaccumulation from mid G2 to a peak at mitotic onset.

The data for genes encoding mitotic regulatory proteins are shown in Fig. 1F, for proteins acting at the G1 to S-phase transition in Fig. 1G, and for other potential regulators of mitotic control in Fig. 1H. The data for G1 to S phase transition proteins excluded data for Cig1, whose G2 levels were too low to meaningfully plot, and Rum1 which was not visible under physiological growth conditions. What is striking about our results is that the vast majority of potentially mitotic regulatory molecules showed almost no change in concentration as cell size increased during the cell cycle. The changes were often less than 1.1×, and at most 1.3×, with no evidence for a consistent increase in concentration as cells increased in size. The two exceptions out of the 38 mitotic control cell cycle proteins assayed were Cdc13 and Cdc25 (Fig. 1D-E & 1F), which change as cell size increases. The concentration of both Cdc13 and Cdc25 peak at mitosis, and so these proteins are candidates for being cell size sensing regulatory molecules at the G2 to mitosis transition as has been suggested previously (11, 17, 33).

The situation with respect to the proteins acting at the G1 to S-phase transition is different. Of the 7 proteins assayed (excuding Cig1 and Rum1), Cdc18, Cig2, Mik1, and Srw1 showed a significant change in concentration of 2-fold or more, though Puc1 was less so (Fig. 1G). They all peak at G1/S and then fall in level as the cells proceed through G2. In the case of Cdc18 and Mik1 their concentration peak as cells proceed through mitosis, while Cig2, Puc1 and Srw1 all peak after mitosis. We conclude that protein concentration changes appear to be more of a feature of the G1 to S-phase transition than is the case for the G2 to mitosis transition. The reductions in concentration of these proteins observed as cells increase in size through G2 to mitosis could in principle indicate that they have a role in co-ordinating cell size increase with mitotic onset, but there is no evidence from our previous screens that the loss of any of these 7 genes influences cell size at mitosis (23, 24).

We next turned from imaging flow cytometry to widefield microscopy which allowed us to investigate changes in cell cycle protein levels as well as spatial distributions. Utilising a camera with a large field-of-view and the use of accurate cell segmentation with the neural network segmentation software *YeaZ* (34) we could image thousands of cells at high spatial resolution. Septating G1 cells were analysed separately even if still physically connected (Fig. 1A).

The proteins analysed in this study cover a broad range of regulatory and biological pathways (23, 24). Visual inspection revealed a range of localisation patterns, including the cytoplasm, nucleus, spindle, spindle pole body (SPB), cell tips and septum (Images and full descriptions of all strains used in this study are available in the Extended Supplement). Of the 47 proteins examined, 36 showed a nuclear localisation, 29 of which accumulated at a higher level in the nucleus compared to the cytoplasm, and 7 of which showed accumulation at the SPB (Cdc2, Cdc13, Clp1, Suc1, Wee, Cig1, Cig2 and Cdr2). For each strain we plotted the whole cell mean intensity versus length and absolute intensity versus length. Focussing first on the mitotic regulatory proteins analysed in Fig. 1F, we confirmed that only Cdc13 (Fig. 2F) and Cdc25 (Fig. 2J) changed their whole cell concentrations to peak at mitotic entry, with patterns of increase across the cell cycle similar to those observed from imaging flow cytometry (Fig. 1F).

**Figure 2.**
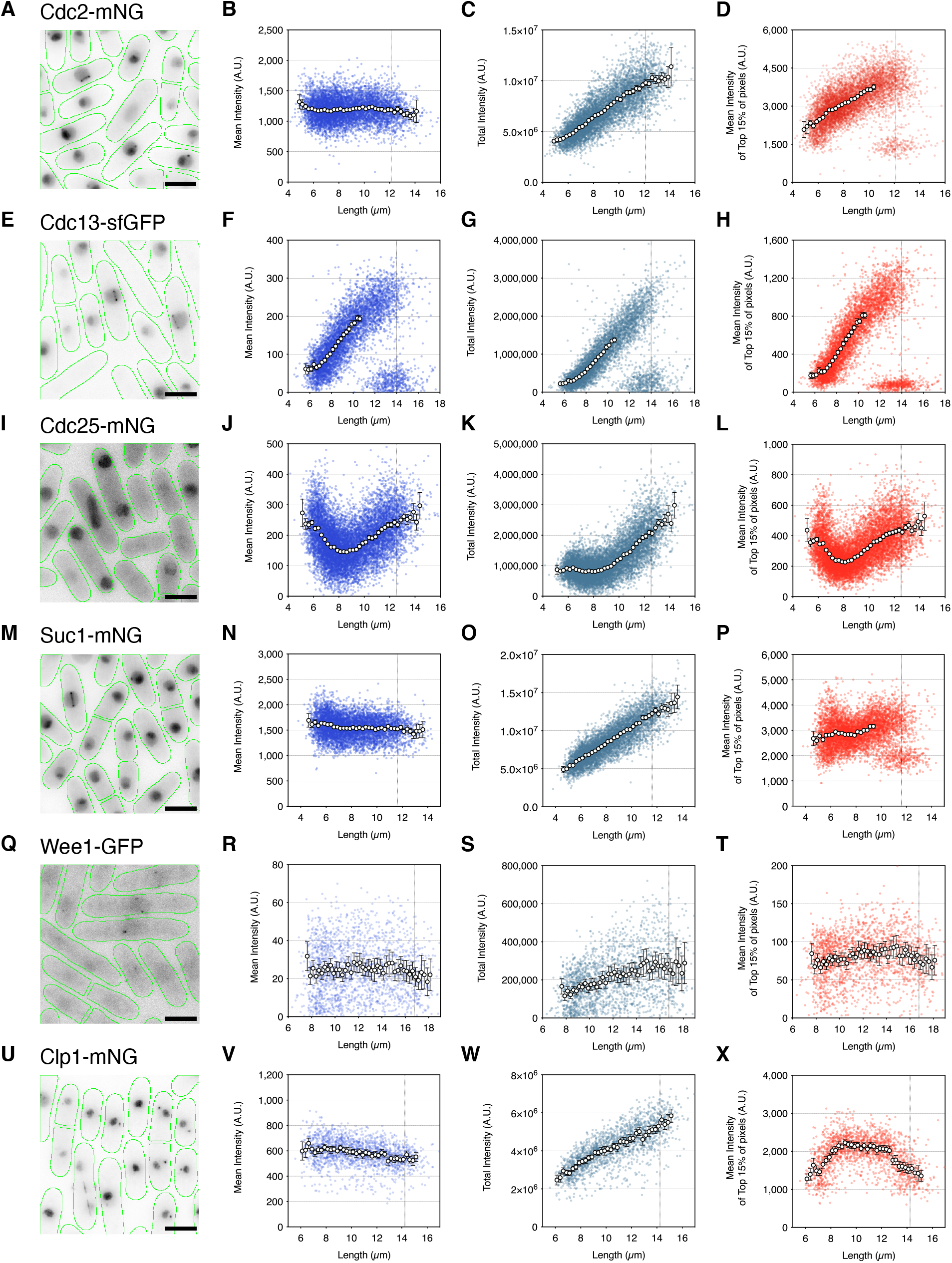
The core mitotic regulators Cdc2, Cdc13, Cdc25, Suc1, and Wee1 increase their localised concentrations to a peak at mitotic entry, whilst Clp1 increases to a peak by early G2. **A**, **E**, **I**, **M**, **Q** & **U**, Images of fission yeast cells (green outline) for Cdc2-mNG (**A**), Cdc13-sfGFP (**E**), Cdc25-mNG (**I**), Suc1-mNG (**M**), Wee1-GFP (**Q**) and Clp1-mNG (**U**) from asynchronous cell cultures each showing a range of cells across the cell cycle. Images are maximum intensity projections. Scale bar 5 μm. **B**-**D**, **F**-**H**, **J**-**L**, **N**-**P**, **R**-**T** & **V**-**X**, Plots show mean whole cell fluorescence intensity (blue), total cellular fluorescence intensity (teal) and mean of the top 15% of cellular pixel values (red) plotted against cell length for Cdc2-mNG (**B**-**D**), Cdc13-sfGFP (**F**-**H**), Cdc25-mNG (**J**-**L**) Suc1-mNG (**N**-**P**), Wee1-GFP (**R**-**T**) and Clp1-mNG (**V**-**X**). Coloured dots represent individual cell values. Circles represent mean values at 0.25 μm length bins with >10 cells/bin. Error bars represent 95% CI. Means stop for **D**, **F**-**H** & **P** before cell populations become bimodally distributed. Vertical dotted line represents septation length. n =**A**-**D**, 6,989 cells; **E**-**H**, 5,895 cells; **I**-**L**, 11,091 cells; **M**-**P**, 6,513 cells; **Q**-**T**, 1,863 cells; **U**-**X**, 1,838 cells.

Next, we turned our attention to changes in the localised concentration of proteins during the cell cycle. Due to the localisation of proteins being specific to each strain we calculated a mean of a top percentage of pixels in order to estimate local changes in concentration. Particular attention was paid to levels in the nucleus as a mitotic regulatory protein at this location could be indicative of an interaction with DNA and thus ploidy sensing. The nuclear volume in fission yeast increase as cells proceed through the cell cycle as a fixed proportion of cell size (9, 35–37). In a 2D image the nucleus occupies approximately 15% of the total cell area, so for strains that appeared to be nuclear localised we approximated the changes in nuclear level from single colour imaging by determining the mean of the top 15% of pixels (nuclei could not be segmented from single colour imaging for proteins that leave the nucleus at mitosis such as Cdc2 and Cdc13). This analysis allowed a rapid and indicative screen of the ‘nuclear’ localisation behaviour of the cell cycle proteins. Of the 29 proteins that preferentially localised to the nucleus, 12 showed concentration changes through the cell cycle (Cdc2, Cdc10, Cdc13, Cdc18, Cdc25, Cig1, Cig2, Clp1, Mik1, Srw1, Suc1 and Wee1). Five of the 12 peaked in concentration at the G2 to mitosis transition: Cdc2 (Fig. 2D), Cdc13 (Fig. 2H), Cdc25 (Fig. 2L), Suc1 (Fig. 2P) and Wee1 (Fig. 2T). Cdc13 (Fig. 2F & H) and Cdc25 (Fig. 2J & L) both showed similar patterns of increase in the ‘nucleus’ as in the whole cell, whilst Cdc2 (Fig. 2B & D), Suc1 (Fig. 2N & P) and Wee1 (Fig. 2R & T) all exhibited ‘nuclear’ concentration increases with increasing cell size while their whole cell concentrations remained constant. Clp1 showed a dynamic change in localisation from the nucleolus and SPB to the spindle (Fig. 2U). This was emulated by the mean of the top 15% of pixels showing a concentration peak in early G2 before being held at a stably high level through to mitosis (Fig. 2X). For Clp1, such an early cell cycle pattern of accumulation is not consistent with models for size sensing at the G2 to mitosis transition.

In the case of the the G1 to S-phase proteins (Fig. 3), Cig2 (Fig. 3A-B) Cdc18 (Fig. 3E-F), Mik1 (Fig. 3G-H) and Srw1 (Fig. 3I-J) also showed whole cell patterns of expression consistent with the imaging flow cytometry data (Fig. 1G). Being able to split septated cells from widefield microscopy images meant that peak intensity for these proteins could be observed more clearly at the G1 to S-phase transition. For Cdc18 (Fig. 3F), Mik1 (Fig. 3H) and Srw1 (Fig. 3J) the highest concentrations were found in the shortest cells indicative of peak levels after binucleation and at the point of septation. For all three proteins this level reduced to a low level by mid G2. For Cdc10, as for imaging flow cytometry, the peak was only slightly elevated in short cells (approximately 1.2×) and was relatively stable though the rest of the cycle (Fig. 3K-L). Cig2 concentration increased from G1 to a peak early in G2. Its level reduced as cells proceeded through the cell cycle to a low at mitosis (Fig. 3A-B). The pattern of whole cell Cig2 expression was matched by its ‘nuclear’ concentration (Fig. 3D). We also report the first visulisation of endogenous Cig1 expression (see Extended Supplement). While its expression level proved too low to accurately quantify by our methods of analysis, the images show visible Cig1-mNG fluorescence at both single and duplicated SPBs in G2, and then transiently throughout the nucleus at anaphase and nuclear division (Fig. 3M). Overall, these results confirm our earlier conclusion that changes in cell cycle protein concentrations is more a feature of the G1 to S-phase transition than for G2 to mitosis.

**Figure 3.**
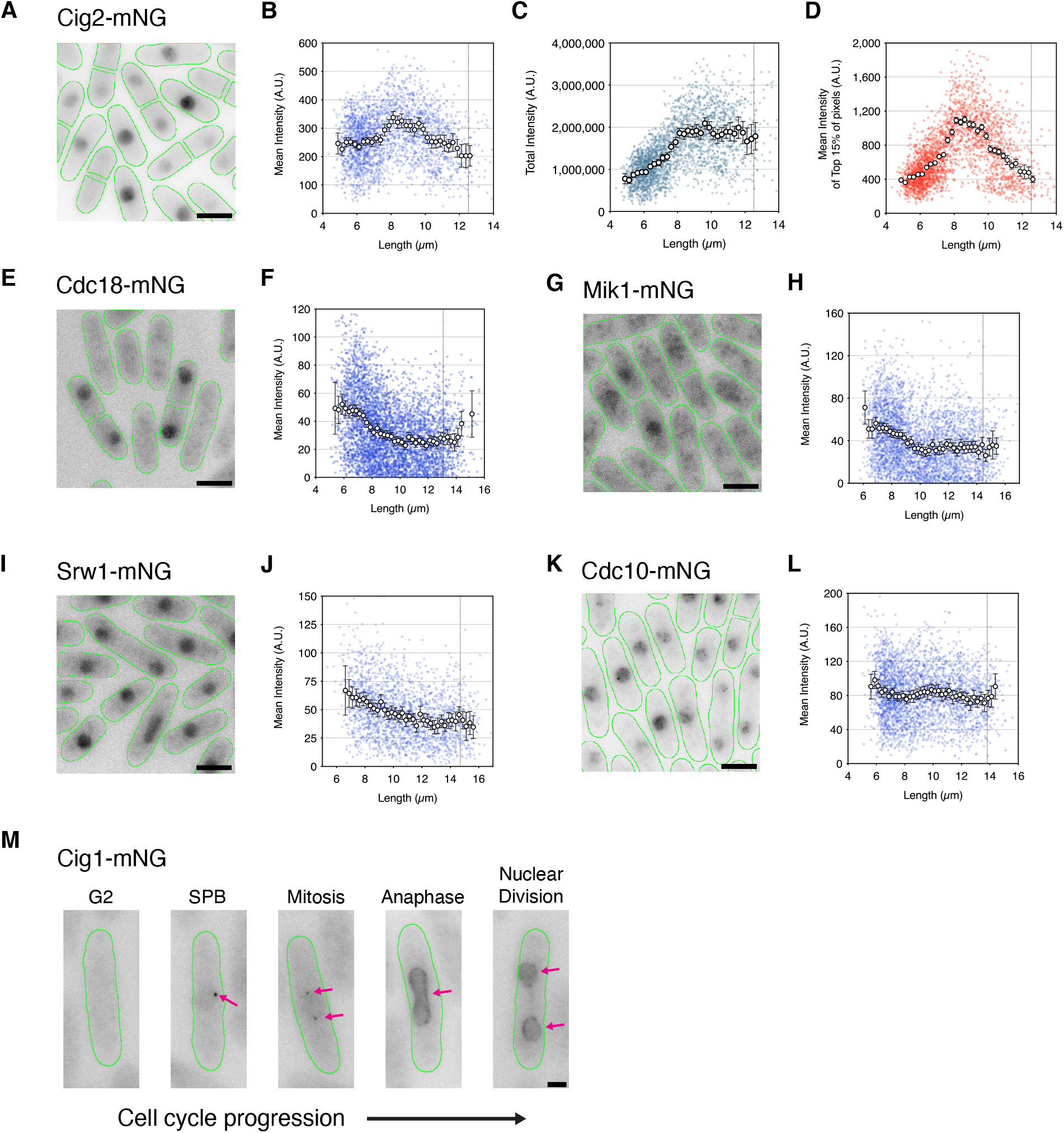
G1 to S phase transition proteins are transiently expressed early in the cell cycle. **A**, **E**, **G**, **I** & **K**, Images of fission yeast cells (green outline) for Cig2-mNG (**A**), Cdc18-mNG (**E**), Mik1-mNG (**G**), Srw1-mNG (**I**) and Cdc10-mNG (**K**) from asynchronous cell cultures each showing a range of cells across the cell cycle. Images are maximum intensity projections. Scale bar 5 μm. **B**-**C**, Cig2-mNG plots show mean whole cell fluorescence intensity (blue), total cellular fluorescence intensity (teal) and mean of the top 15% of cellular pixel values (red) plotted against cell length. **F**, **H**, **J** & **L**, Mean whole cell fluorescence intensity plots against cell length for Cdc18-mNG (**F**), Mik1-mNG (**H**), Srw1-mNG (**J**) and Cdc10-mNG (**L**). Coloured dots represent individual cell values. Circles represent mean at 0.25 μm length bins with >10 cells/bin. Error bars represent 95% CI. Vertical dotted line represents septation length. n =**B**-**D**, 2,953 cells; **F**, 4,644 cells; **H**, 3683 cells; **J**, 1,946 cells; **L**, 2,978 cells. **M**, representative images of Cig1-mNG cells (green outline) at progressive cell cycle stages. Images are maximum projections normalised to peak intensity level (at SPB). Pink arrows indicate Cig1-mNG localisation. Scale bar, 2 μm.

Our ‘nuclear’ screen suggested that 5 members of the core set of cell cycle CDK regulators Cdc2, Cdc13, Cdc25, Suc1 and Wee1 increase in nuclear concentraton to a peak at mitotic entry. To confirm nuclear localisation and quantify precise nuclear levels we used a dual-colour imaging approach to image each fluorescently tagged regulator alongside Cut11-mCherry, a component of the nuclear core complex, to mark the nucleus for segmentation. Fission yeast are particularly amenable to tracking of nuclear localisation patterns throughout the whole cell cycle due to having a closed mitosis. We used a combination of whole cell neural network segmentation with *YeaZ*, and machine learning nuclear segmentation with *Ilastik* (38) (Fig. S2). Examining cells from asynchronous populations from early G2 through to septation, allowed us to visually determine how the localisation of these regulators changed (Fig. 4 & Fig. S3). Taking Cdc2-mNG as an example (Fig. 4A), for cells in G2 the signal can be seen to be nuclear enriched (Fig. 3A) and to accumulate on the SPB, as previously observed (39). After SPB duplication, Cdc2 concentrates on both SPBs and the connecting spindle. As the nucleus elongates in anaphase, Cdc2 is exported from the nucleus, prior to reaccumulation in the next cycle after nuclear division. In the associated dot plot (Fig. 4B), the mean cellular concentration for each cell (represented in blue) remains constant throughout the cell cycle consistent with previous whole cell measurements (Fig. 1F & 2A). Mononucleate nuclear Cdc2 concentration (green) increases with cell length thoughout the cell cycle (Fig. 4B) reaching a peak at mitotic entry. In long mononucleates that are >12 μm, Cdc2 levels decrease again, thus indicating that Cdc2 leaves the nucleus in mitosis. Pink dots represent the nuclear concentration of binucleate cells that have not yet septated and show that nuclear levels of Cdc2 post mitosis reduce to a level equal to the whole cell concentration.

**Figure 4.**
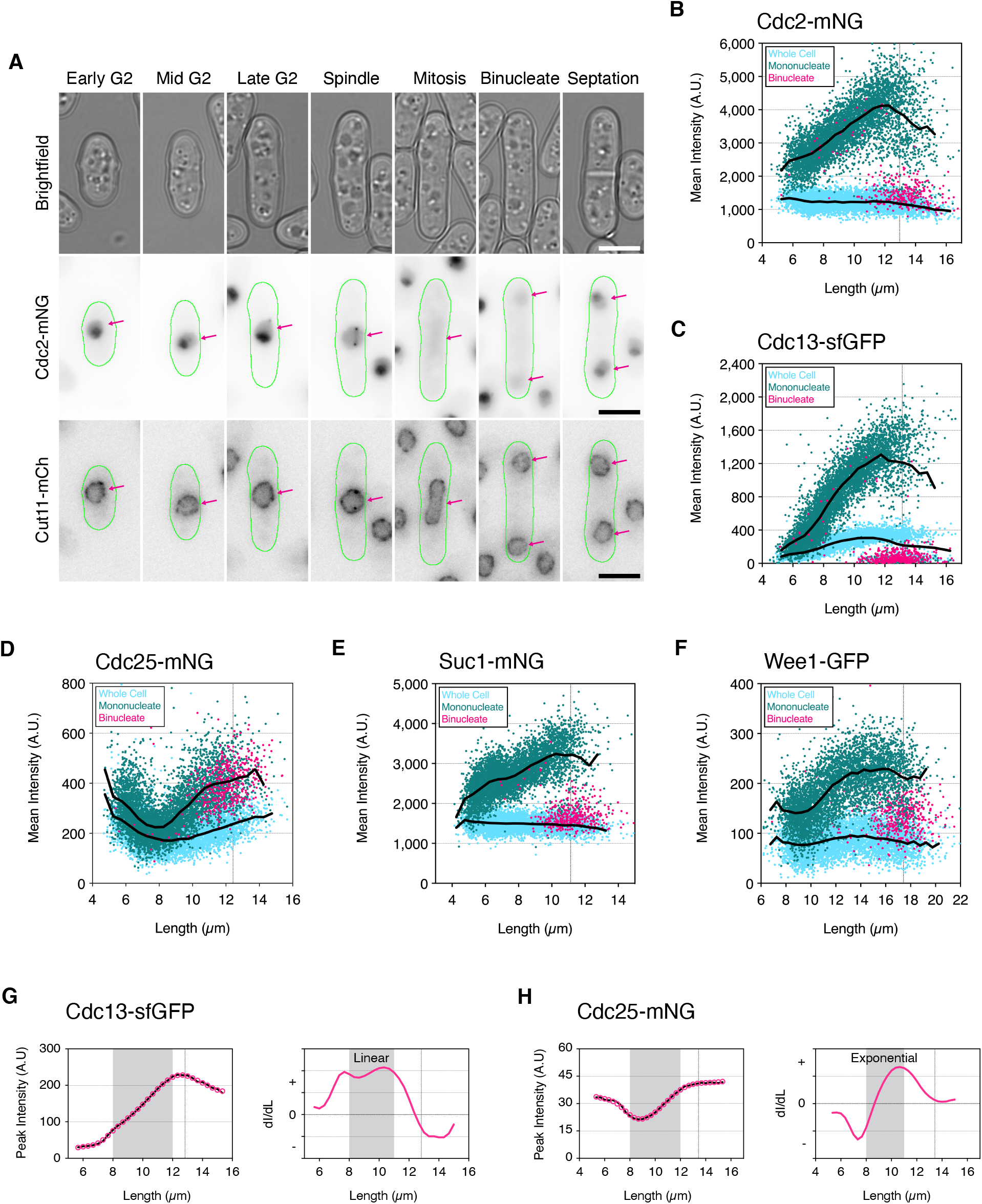
The core mitotic regulators increase their nuclear concentrations to a peak in late G2. **A**, Montage showing representative Cdc2-mNG cells (green outline) selected from an asynchronous population at progressive stages of the fission yeast cell cycle from early G2 through to septation. Top, Brightfield; Middle, Inverted fluorescence images for Cdc2-mNG; Bottom, Inverted fluorescence images for Cut11-mCh (nuclear marker). mNG intensities are maximum intensity projections normalised min-to-max for all pixels within the montage. Cut11-mCh images are maximum intensity projections re-normalised from 10,000 to 50,000-pixel values (64-bit images). Pink arrows indicate nuclei position for comparison of Cdc2 localisation. Scale bars = 5 μm. **B**-**F**, Plots showing mean cellular fluorescence intensity (light blue), mean nuclear fluorescence for mononucleate cells (green) and binucleates (pink) against cell length for Cdc2-mNG (**B**), Cdc13-sfGFP (**C**), Cdc25-mNG (**D**), Suc1-mNG (**E**) and Wee1-GFP (**F**). Coloured dots represent individual cell values. Black lines represent connected mean values calculated at 0.5 μm bins for whole cell values (bottom), and nuclear mononucleate values (top). Note, binucleates with a septum are split during segmentation with each half treated as an individual mononucleate. Vertical dotted line indicates septation length. n =**B**, 5,857 cells; **C**, 6,235 cells; **D**, 5,920 cells; **E**, 5,832 cells; **F**, 5,189 cells. **G & H**, Rate of change plots for nuclear accumulation of Cdc13-sfGFP (**F**) and Cdc25-mNG (**G**). For each strain, plots on the left show mean peak nuclear intensity (Peak Intensity) against cell length. Raw data values in black, smoothed data values in pink. Plots on the right are the rate of change of Intensity of smoothed data (dI)/Cell Length (dL) from each unit of length to the next. Positive values indicate accumulation, whilst negative points indicate loss. Increasing values indicate an increasing rate of accumulation. Values at 0 indicate a stable concentration. Vertical dotted line on each plot indicates cell length at mitosis. n = **G**, 194,326 cells; **H**, 223,364 cells.

Nuclear concentration patterns for Cdc13 (Fig. 4C), Cdc25 (Fig. 4D), Suc1 (Fig. 4E), Wee1 (Fig. 4F) match their ‘top 15%’ analyses (Fig. 2H, L, P, T) and establish that these core CDK mitotic regulators all increase in concentration within the nucleus as cell size increases. For Cdc2 this appears to be a continuous increase correlated with size and more than doubles in concentration from early G2 to mitosis (Fig. 4B). For Suc1, nuclear concentration increases at the beginning of the cell cycle, plateaus in mid-G2 and then reaccumulates into mitosis, increasing 1.5× across the full cycle (Fig. 4E). Likewise, Wee1 nuclear concentration also increases 1.5× with a continuous increase to mid-G2 followed by a plateau until mitosis (Fig. 4F). Unlike Cdc2, Cdc13 and Suc1, which show a rapid exit from the nucleus at mitotic exit as shown by nucleus levels of binucleate cells matching the level of the whole cell, Wee1 appears to be only partially exported as indicated by binucleate nuclear levels being raised above the level of the whole cell. For Cdc25 in Fig. 2J we showed that whole cell levels increased nearly two-fold from mid-G2 to mitosis and this is recapitulated in Fig. 4D. Nuclear accumulation of Cdc25 follows a similar pattern but at a higher concentration than compared to whole cell levels, with nuclear levels again increasing around two-fold from mid-G2 to mitosis and then maintained at a high level after binucleation. Cdc25 nuclear export begins at the point of septation, with faint puncta observed at the nuclear periphery suggestive of active nuclear export (Fig. S3B). Nuclear levels gradually decrease in concentration in early G2 (in short cells < 8 μm).

The protein that changed most significantly was the mitotic B-type cyclin Cdc13. In the whole cell the span of the increase is approximately 4-5× (Fig. 2D and Fig. 4C) which is amplified in the nucleus to 8-10× (Fig. 4C). This makes Cdc13 the best candidate cell size sensor due to its high dynamic range through the cell cycle, particularly in the nucleus.

Returning to our imaging flow cytometry data, we looked to measure the nuclear rate of change for Cdc13 and Cdc25. From mid G2 towards mitotic entry (8-12 μm) we find that nuclear accumulation of Cdc13 occurs at an almost constant linear rate after a rise at the beginning and a fall at the end of the cell cycle (Fig. 4G). In contrast, the rate of Cdc25 nuclear accumulation increases in an exponential manner during the second half of the cell cycle (Fig. 4H). Cdc13 and Cdc25 both activate the G2-M CDK but in different ways: Cdc13 forms a complex with Cdc2 for direct activation, while Cdc25 phosphatase activates by removing the inhibitory Cdc2-Y15 phosphorylation. This suggests that the mode of CDK regulation during the cell cycle switches from a mode that is predominantly dependent upon cyclin accumulation to a mode that is additionally subject to the futile cycle of the activating Cdc25 phosphatase counteracting the inhibitory action of Wee1 protein kinase.

## Discussion

In this paper we have determined how the levels of potential mitotic regulators change during the cell cycle in both the whole cell and the nucleus of fission yeast, with the aim of informing our understanding of how cells sense their size and regulate the dynamics of CDK activation at the onset of mitosis. We measured the levels of 30 proteins previously identified as potential mitotic regulators (23, 24) throughout the cell cycle together with 8 more proteins implicated in mitotic control, and a further 7 involved in the G1-S phase transition. Surprisingly, of the 38 G2 to mitosis regulators, 36 remained constant in concentration within the whole cell throughout the cell cycle. Therefore if they were to have a influence on cell cycle progression through CDK regulation this would have to be due to dynamic changes in their activity status, for example by changing phosphorylation levels or through changes in their spatial distribution. Only two mitotic regulators, the mitotic B-type cyclin Cdc13 and the activating Cdc25 phosphatase, change in whole cell levels with both peaking at the onset of mitosis.

The behaviour of the 7 proteins involved in the G1 to S-phase transition was different, with 4 changing dynamically in the whole cell during the cell cycle peaking at G1-S. We propose that changes in the level of protein concentration is more a a feature at the G1 to S-phase transition than G2 to mitosis. These could be involved in cell size sensing at the G1 to S-phase transition because there is a cell size control acting at this checkpoint in fission yeast, although it is normally cryptic and does not normally influence cell size at mitosis and cell division (40, 41). In budding yeast, several models for G1 to S-phase cell size sensing invoke the need for S-phase regulators to attain a critical concentration before S-phase can commence (12, 42, 43).

Over three quarters of the cell cycle proteins investigated were primarily located in the nucleus during interphase and 29 of them accumulated in the nucleus to a higher level than the cytoplasm. This nuclear enrichment could be due to critical cell cycle events occurring in the nucleus in a closed mitosis, but also given the nuclear localisation of DNA these proteins could also be candidates for being involved in ploidy sensing mechanisms. Seven proteins were also associated with the spindle pole body, including core CDK regulators (Cdc2, Cdc13, Clp1, Suc1, Wee1, Cig1, Cig2 and Cdr2), providing support for the long held view that the centrosome plays a critical role in cell cycle control (44, 45). Nine of the 29 nuclear-located proteins undergo significant changes during the cell cycle, six of which are core CDK regulators (Cdc2, Cdc13, Cdc25, Suc1, Wee1 and Cig2). This finding supports work from mammalian systems that cell cycle regulated nuclear import plays an important role in allowing the cell to reach the point of mitotic onset by gradually bringing about the nuclear accumulation of mitotic CDK regulators (46). Consistent with nuclear transport playing a regulatory role is the fact that 6 of the 17 genes identified in the haploinsufficient screen are involved in nuclear transport (23).

The control of CDK activity required for mitotic onset is highly dependent upon both the level of the Cdc2 activating mitotic B-type cyclin Cdc13, and the regulatory feedback loop consisting of the counteracting phosphatase activator Cdc25 and the protein kinase inhibitor Wee1 (27, 47). This regulatory loop determines the extent of the inhibitory Cdc2 Y15 phosphorylation. We have shown that the whole cell concentration of Wee1 remains reasonably constant, but Cdc25 levels oscillate significantly through the cell cycle. This suggests that the activating potential of the Cdc25/Wee1 regulatory loop follows the Cdc25 concentration profile by being high at the beginning of the cell cycle, at a time when cyclin-CDK nuclear levels are low, falling to a low by mid-G2 and then rising towards mitosis onset. When Wee1 is inactivated by a temperature shift of a Wee1^ts^ mutant, cells in the second half of G2 are immediately advanced into mitosis (26). This indicates that the Cdc25/Wee1 regulatory loop is restraining mitotic onset during the second half of G2, even though there is sufficient CDK formed by the Cdc13/Cdc2 complex in mid-G2 for mitotic onset to take place.

Cdc13 concentration rises dramatically throughout the cell cycle, much more than any other protein investigated in this study. This level rises 4-5× in the whole cell and 8-10x in the nucleus. At the point in mid-G2 when the Cdc25/Wee1 regulatory loop begins to rise, Cdc13 concentration is about two thirds of the maximum reached at the onset of mitosis. We conclude that this level of cyclin is sufficient to generate enough CDK activity to undergo mitosis, but is restrained by the Cdc25/Wee1 regulatory loop until late in G2. The difference in Cdc13 concentration from mid G2 to mitotic onset is about one third, which could correspond to a ‘CDK buffer zone’ where cells undergo mitosis with a higher level of CDK activity than is strictly necessary for the completion of mitosis (17, 48).

We propose that the increasing level of Cdc13 is primarily responsible for cell size sensing at the G2 to mitosis transition (17) and that this is the major mechansism by which eukaryotic cells maintain cell size homeostasis at cell division. As B-type cyclin increases in concentration throughout the cell cycle and peaks at mitosis in all eukaryotes investigated thus far, this could be a universal cell size sensing mechanism, with the advantage over cell geometry sensing mechanisms (13–16) of being independent of a fixed cell shape, and therefore having the potential to be applicable to all cell types including metazoans.

The observations we have reported here also have implications for the dynamics of CDK regulation at the onset of mitosis. We have proposed that the increasing level of the B-type cyclin Cdc13 is primarily responsible for cell size sensing at the G2 to mitosis transition, with a possible further contribution provided by the increasing level of Cdc25 during the later part of G2 (33). Our analysis suggests that the minimum level of Cdc13 required for mitosis and cell division is established in mid-G2 when the cell attains a threshold size. However the linear increase in Cdc13 level from mid G2 to mitosis likely results in a change in CDK activity which is too gradual at the end of the cell cycle to bring about an orderly sharp transition into mitosis. We propose that the cell solves this problem by a two-step process. The first step is based on B-type cyclin Cdc13 accumulation that can accurately monitor cell size whilst Cdc25 levels are reducing and low. The second step, activated in mid-G2, is based upon the Cdc25/Wee1 regulatory loop. The increasing Cdc25 level sets up a futile cycle introducing bistability which results in a rapid rise in CDK activity and a sharp orderly progression through the multiple events of mitosis, thereby reducing the variability in size at division. Accumulation in the nucleus in a size dependent manner could also amplify any potential cell size sensing mechanism (46, 49). In all eukaryotes studied so far, mitotic B-type cyclin rises significantly throughout the cell cycle and a Cdc25-like phosphatase with a Wee1 protein kinase regulates CDK activity through control of tyrosine phosphorylation of the CDK protein kinase (50–52). Therefore this two-step model of mitotic B-type cyclin accumulation through the cell cycle sensing cell size, coupled with a subsequent Cdc25/Wee1 futile cycle introducing bistability and rapid CDK activation, may be a universal feature of eukaryotic cell cycle control given the conservation of the control elements throughout eukaryotes.

## Acknowledgments

We would like to thank the Silke Hauf and Yasushi Hiraoka labs for sharing of *S. pombe* strains; Sahand Jamal Rahi for training of *YeaZ* for fission yeast cell segmentation; James Patterson (previous Nurse lab) for construction and sharing of plasmids; members of the Nurse lab: Thomas Hammond, Emma Roberts & Theresa Zeisner and Tiffany Mak (previous member) for critical reading of the manuscript and comments. S.C. and P.N. were supported by the Francis Crick Institute which receives its core funding from Cancer Research UK (FC01121), the UK Medical Research Council (FC01121), and the Wellcome Trust (FC01121). In addition, this work was supported by the Wellcome Trust grant to P.N. [Grant numbers: 214183 & 093917]. G.D. acknowledges UCL, the Wellcome Trust (203276/Z/16/Z), and the European Molecular Biology Laboratory for support. P.R. acknowledges the support of the Biotechnology and Biological Sciences Research Council under grant BB/P026818/1 and the Engineering and Physical Science Research Council under grant EP/N013506/1.

## Author Contributions

S.C. designed and implemented all experiments, generated strains, acquired, analysed and interpreted data, and co-drafted the manuscript. P.N. conceived and supervised the project, provided advice on experimental design and analysis, and co-drafted the manuscript. G.D. advised on experimental analyses and wrote MATLAB scripts for the heatmap production and widefield image analyses. P.R. perfomed the derivative analysis.

---

**Supplementary Figure 1.**
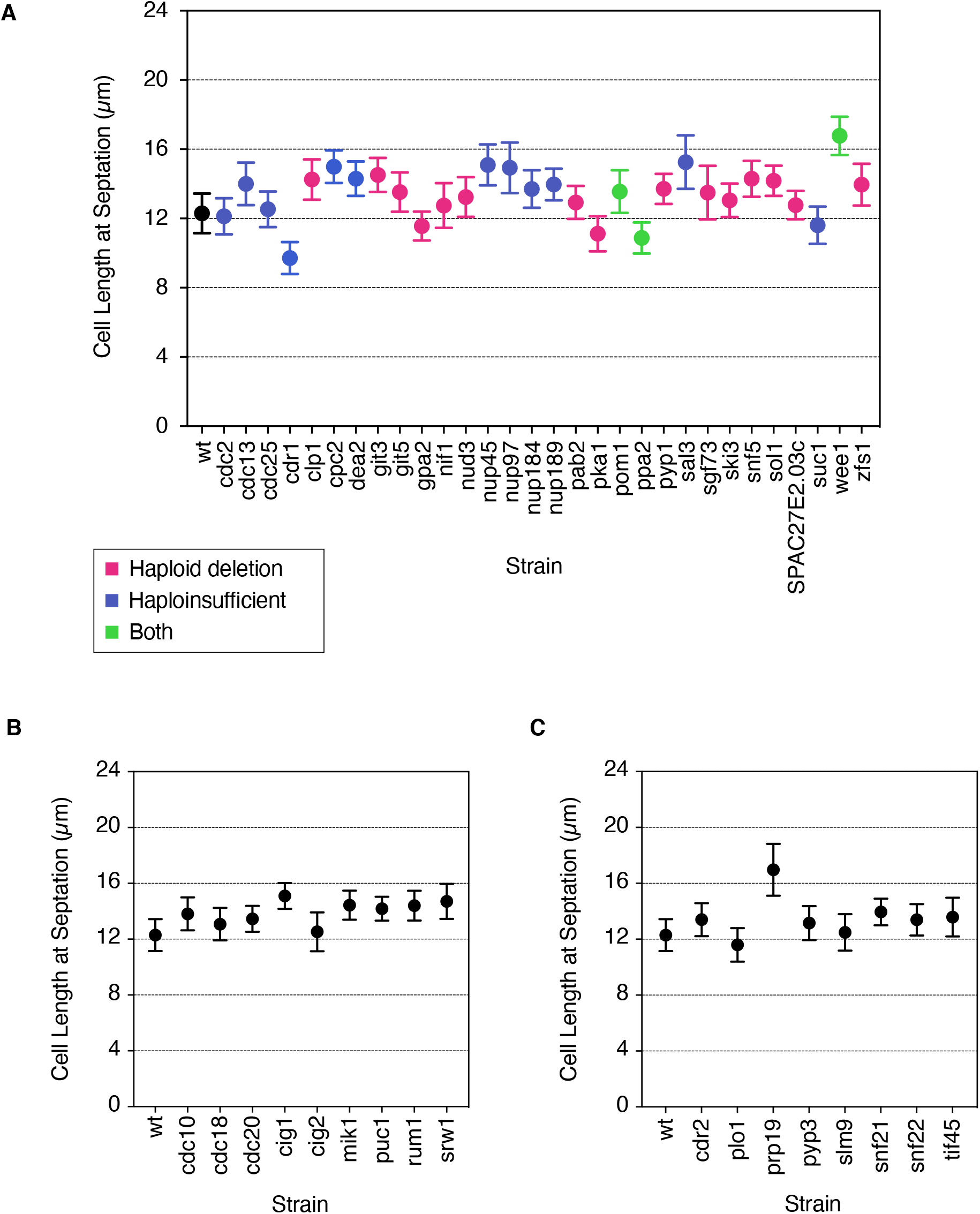
Cell length at septation for all widefield microscopy imaged strains in this study. **A**, Plot showing the quantification of cell length at septation for fluorescently tagged mitotic regulators. Colour indicates whether the tagged protein was included in this analysis due to its presence in our laboratory’s previous haploid deletion screen (pink) (24), heterozygous deletion haploinsufficiency screen (blue) (23), or both (green). Dots indicate mean, Error bars indicate S.D. n ranges from 100 - 757 cells per strain. All strains tagged with mNeonGreen except for Cdc13 (internal sfGFP), Wee1 (N-terminal GFP) and Pka1 (C-terminal GFP). **B**-**C**, as for **A** except for genes associated with the G1 to S-phase transition (**B**) or for other genes of interest involved in mitotic control (**C**). n = **B**, 81 - 608 cells per strain and **C**, 70 - 336 cells per strain. **B**-**C**, All strains are tagged with mNeonGreen.

**Supplementary Figure 2.**
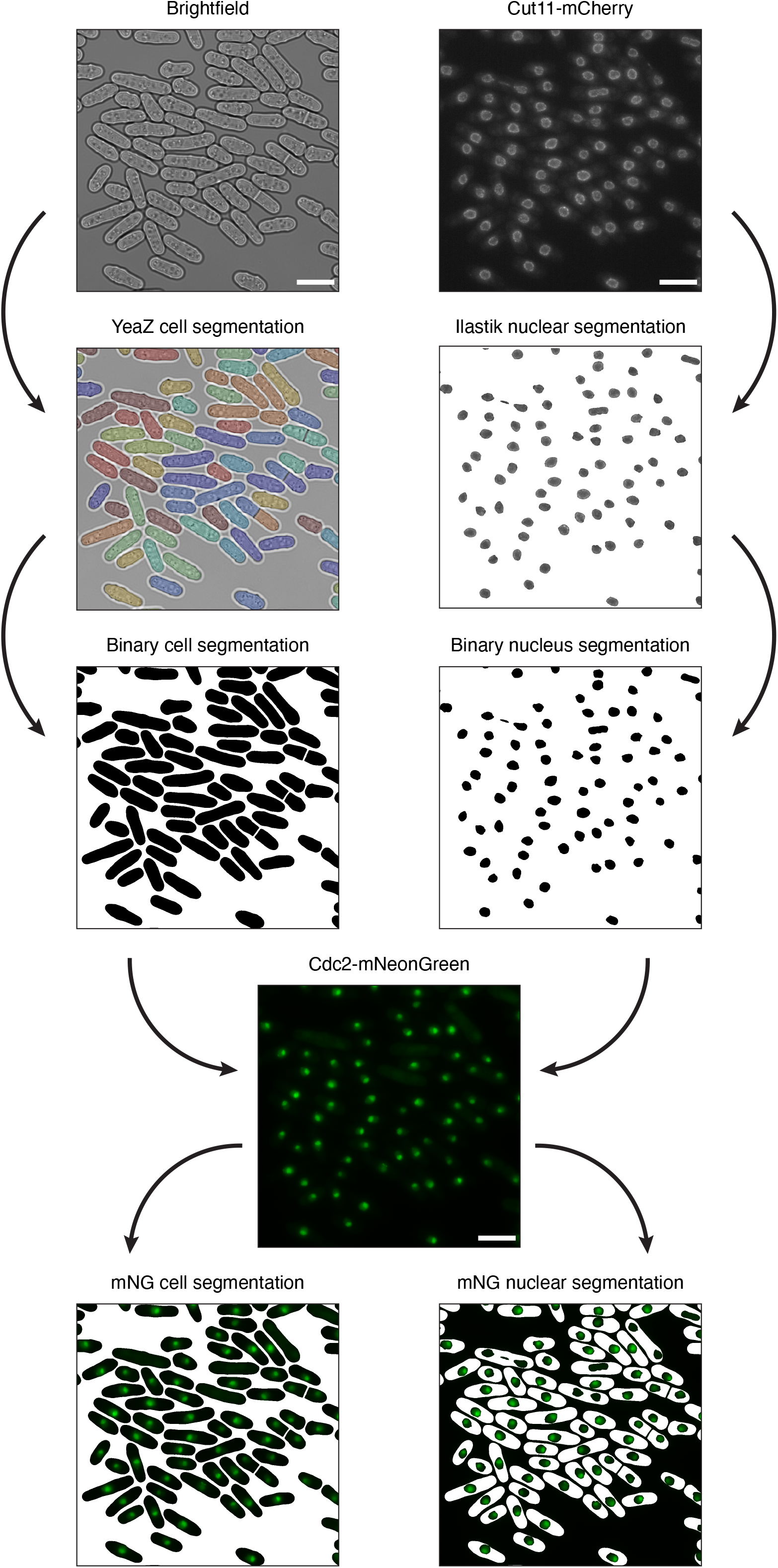
Schematic of the whole cell and nuclear segmentation pipeline using *YeaZ* and *Ilastik* for widefield imaging. Along the left, brightfield images of whole cells are segmented with *YeaZ*. Cells with septa are split into two individual cells to allow for clearer analysis of localisation and level changes occurring at the G1 to S-phase transition. Cell segmentation is exported as a binary image. Along the right, Cut11-mCh (nuclear marker) is segmented with *Ilastik* and converted to a binary image. *YeaZ* and *Ilastik* binary masks are combined and overlaid onto the tagged protein-of-interest (in this example Cdc2-mNG) to allow for whole cell measurements (bottom left), or nuclear cell-associated measurements (bottom right).

**Supplementary Figure 3.**
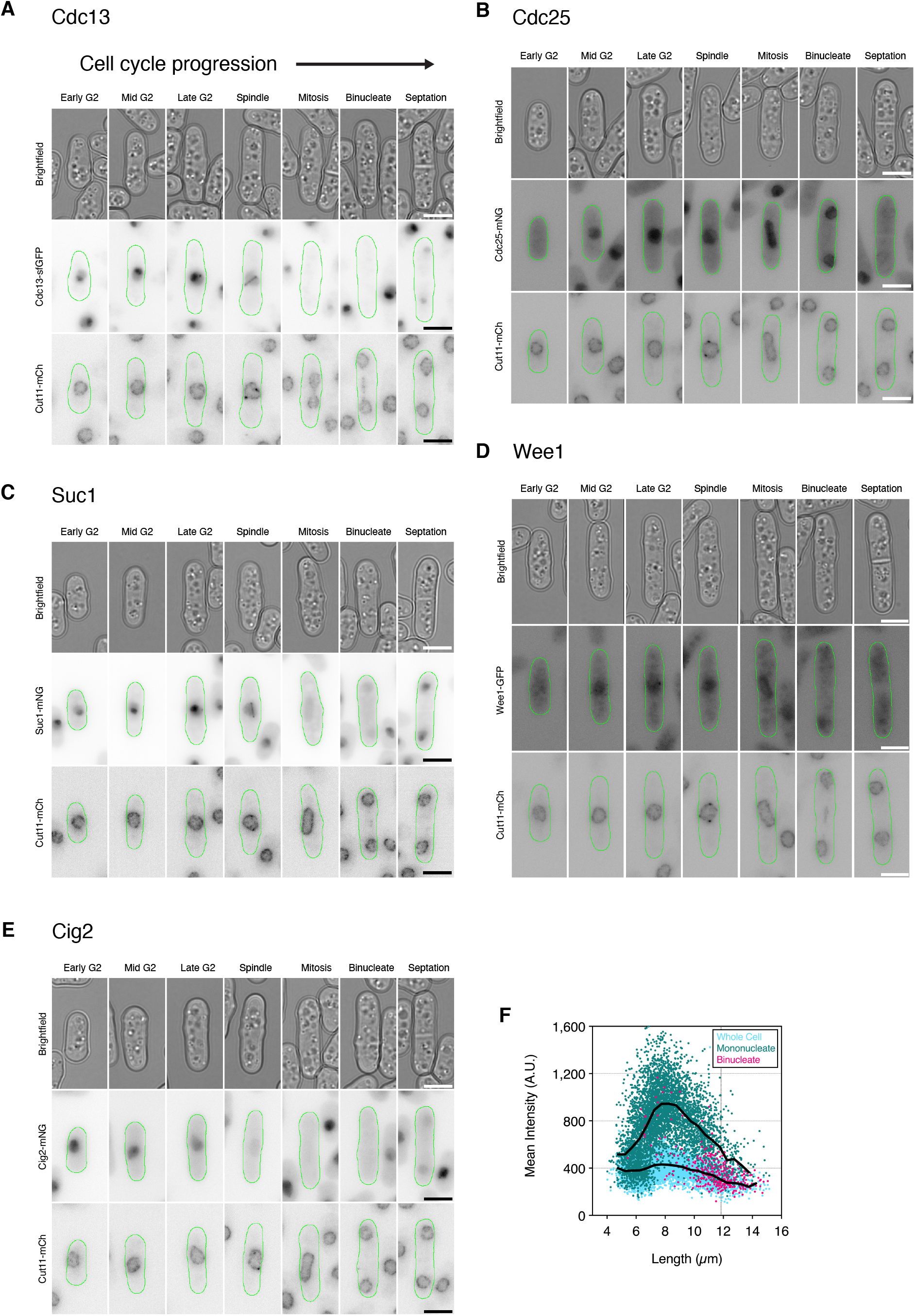
Cell cycle montages showing nuclear accumulation of core mitotic regulators. Montages showing representative cells (green outlines), selected from asynchronous populations at progressive stages of the fission yeast cell cycle from early G2 through to septation for Cdc13-sfGFP (**A**), Cdc25-mNG (**B**), Suc1-mNG (**C**), Wee1-GFP (**D**) and Cig2-mNG (**E**). Top, Brightfield images; Middle, Inverted fluorescence images; Bottom, Inverted fluorescence images of Cut11-mCh (nuclear marker). Fluorescence images for mNG and GFP are maximum intensity projections normalised min-to-max for all pixels within each montage. Cut11-mCh images are maximum intensity projections normalised from 10,000 to 50,000-pixel values (64-bit images). Scale bars = 5 μm. **F**, Plot showing mean cellular fluorescence intensity (light blue), mean nuclear fluorescence for mononucleate cells (green) and binucleates (pink) plotted against cell length for Cig2-mNG. Black lines represent connected mean values calculated at 0.5 μm bins for whole cell values (bottom), and nuclear mononucleate values (top). Vertical dotted line indicates cell length at septation. n = 5,221 cells.

## Methods

### Lead Contact and Materials Availability

Further information and requests for resources and reagents should be directed to Paul Nurse.

All strains, plasmids and reagents generated in this study are available without restriction.

### *Schizosaccharomyces pombe* culture

Cells were cultured by standard growth methods as previously described (53) first on solid YE4S agar and then in liquid Edinburgh Minimal Media supplemented with 20.0 g/L Dextrose Anhydrous (Fisher Scientific BP350-1) and 5.0 g/L Ammonium Chloride (Sigma A9434) (supplemented with adenine, leucine, histidine and uracil to a concentration of 0.15 g/L for autotrophic markers if required) added post-autoclaving and filtered (0.22 μm). Cells were grown at 25°C and maintained in exponential growth between 2 × 10^6^ and 1 × 10^7^ cells/ml (ideally imaged at ~5 × 10^6^ cells/ml).

### Strains

See Table S1 for complete list and full genotypes of *S. pombe* strains used in this study. Strains generated specifically for this study were constructed using standard methods (53, 54). C-terminal gene tagging was performed as previously described (55) using standard primers (Listed in Table S2) designed from the Bähler lab web-interface (bahlerlab.info/resources) for pFA6a vector templates carrying kanamycin or GFP-hygromycin resistance casettes using the lithium acetate method of transformation. pFA6a-Kan vectors were modified by James Patterson for mNeonGreen tagging which carry a non-standard linker (GATTCTGCTGGATCAGCTGGC) upstream of mNeonGreen with the standard forward linker (CGGATCCCCGGGTTAATTAA) replaced.

### Fluorescence Microscopy

For widefield microscopy, 1.5 ml of 0.4-0.5 OD^595^ culture was centrifuged at 3,000 rpm for 30 s, with the supernatant removed and the process repeated. 1.75 μl of pelleted cells were plated on a microscope slide, spread and flattened with addition of a coverslip. Live cells were imaged within 10 mins of preparation. All microscopy was performed at 25°C using a Nikon Ti2 Inverted microscope with a 100X Plan Apochromat oil immersion objective (NA 1.45), Perfect Focus System, a Prime sCMOS camera (Photometrics) and Okolab environmental chamber. The microscope was controlled with Micro-Manager v2.0 software (56). Fluorescence excitation was performed with a SpectraX LED light engine (Lumercor) fitted with standard filters: 470/24 for imaging mNG/sfGFP/GFP and 575/25 for mCherry with a dual-edge Chroma 59022bs, ET - EGFP/mCherry dichroic beamsplitter. Emission filters used were Chroma, ET - EGFP single-band bandpass filter (ET525_50m) for mNG/sfGFP/GFP and Semrock, 641/75 nm Brightline single-band bandpass filter (FF02_641_75) for mCherry.

### Widefield Image Processing

Basic image processing was carried out with FIJI software (57). Whole cell segmentation was performed using the convolutional neural network *YeaZ* (34) trained for fission yeast segmentation on brightfield image slices 1 μm below the focal plane. Using FIJI, maximum intensity projections were made of sfGFP/GFP/mNG/mCherry images covering 2 μm around the focal plane. The machine learning tool *Ilastik* (38) was trained and used to segment maximum projection mCherry images to create nuclear masks. For background correction, controls without the fluorescent tag of interest for each experiment were imaged to give a measure of mean autofluorescence. This value was subtracted from whole cell and nuclear concentration measurements.

### Imaging flow cytometry and post-processing

Cells were imaged for brightfield and fluorescence (488 laser at 400 mW) with an Amnis Imagestream X Mk II Imaging Flow Cytometer with a 60X objective lens. Cells were concentrated from asynchronous exponentially growing cultures (OD^595^ ~0.4-0.5) by centrifugation (3000 rpm/30 s), resuspended in >50 μl of media and waterbath sonicated for 15 s. Prior to acquisition, cells were gated based upon BF Gradient RMS values (value of cell focus) of 65 - 78, and Area/Aspect Ratio values consistent with single cells. Approximately 250,000 gated cells were acquired in 10 - 15 min experiments with an acquisition rate of 300 - 500 gated cells/s.

Post-acquisition processing was undertaken with Amnis IDEAS 6.2 data exploration and analysis software. Cell segmentation masks were created from BF images: (Erode(MO1, 3) named Pombe_Mask and overlaid onto fluorescent images (Ch02).

Cells were further gated based on:

R1: Width_Pombe X coordinates: 3.75 - 6.75
R2: Thickness_Max_Pombe of R1 X coordinates: 3.75 - 6.75
R3: Thickness_Min_Pombe of R2 X coordinates: 3.1 - 5.5
R4: Gradient_RMS_MO1_Ch01 of R3 X coordinates: 65 - 78
R5: Intensity_Pombe_Ch02: removal of extremes on a strain-by-strain basis

Background correction was based upon imaging of a non-fluorescent control on each experimental day. Mean auto-fluorescence per length bin was calculated as a linear regression and used to calculate the level of background to be subtracted from total, mean and max pixel fluorescent values.

### Derivative analyses

To calculate the d(intensity)/d(length) values at the required lengths we first calculated the density of the experimentally measured points in the two-dimensional intensity vs length space. We used the *kde2d* Matlab routine which uses a second order Gaussian kernel with an automatic bandwidth selection method (58). The peak density at each length is then determined to give the intensity value which is then differentiated with respect to length.

### Calculation of the number of nuclei and nuclear intensity

Using a previous approach (8, 17) we extracted the individual cell tiff images for the brightfield and each fluorescence channels from the .cif file generated by the ImageStream X instrument. A cell mask is generated from the brightfield image using a simple grayscale Otsu threshold for this image. The medial line for the cell mask is defined and the nuclear marker fluorescence intensity measured at each point along this line. A simple peak detection algorithm is used to detect if the cell is mono or binucleated. The fluorescence intensity level for each nucleus is simply determined by measuring the pixel value from the fluorescence intensity at this position along the medial line.

### Graphs and Statistics

A custom MATLAB script was used for whole cell, nuclear and top percentage background-subtracted fluorescence measurements for widefield imaging (script author GD - this study). This script counted the number of nuclei per cell with cells having >2 nuclear objects considered binucleate (Ilastik segmentation often found pieces of nuclear bridge). All plots and statistical analyses were performed with Graphpad Prism 9 except for heatmaps (Fig. 1), which were produced by a custom MATLAB script of background-subtracted whole cell mean data collected from imaging flow cytometry. Statistical analyses used and n numbers are stated in the figure legends. Source data is provided with this paper.

## Code availability

Custom MATLAB scripts used in this study are freely available on Github in a public repository at GitHub.com/scottcurran10/fission-yeast-cell-cycle. The use of this code is governed by an MIT license.

---

**Table S1.**
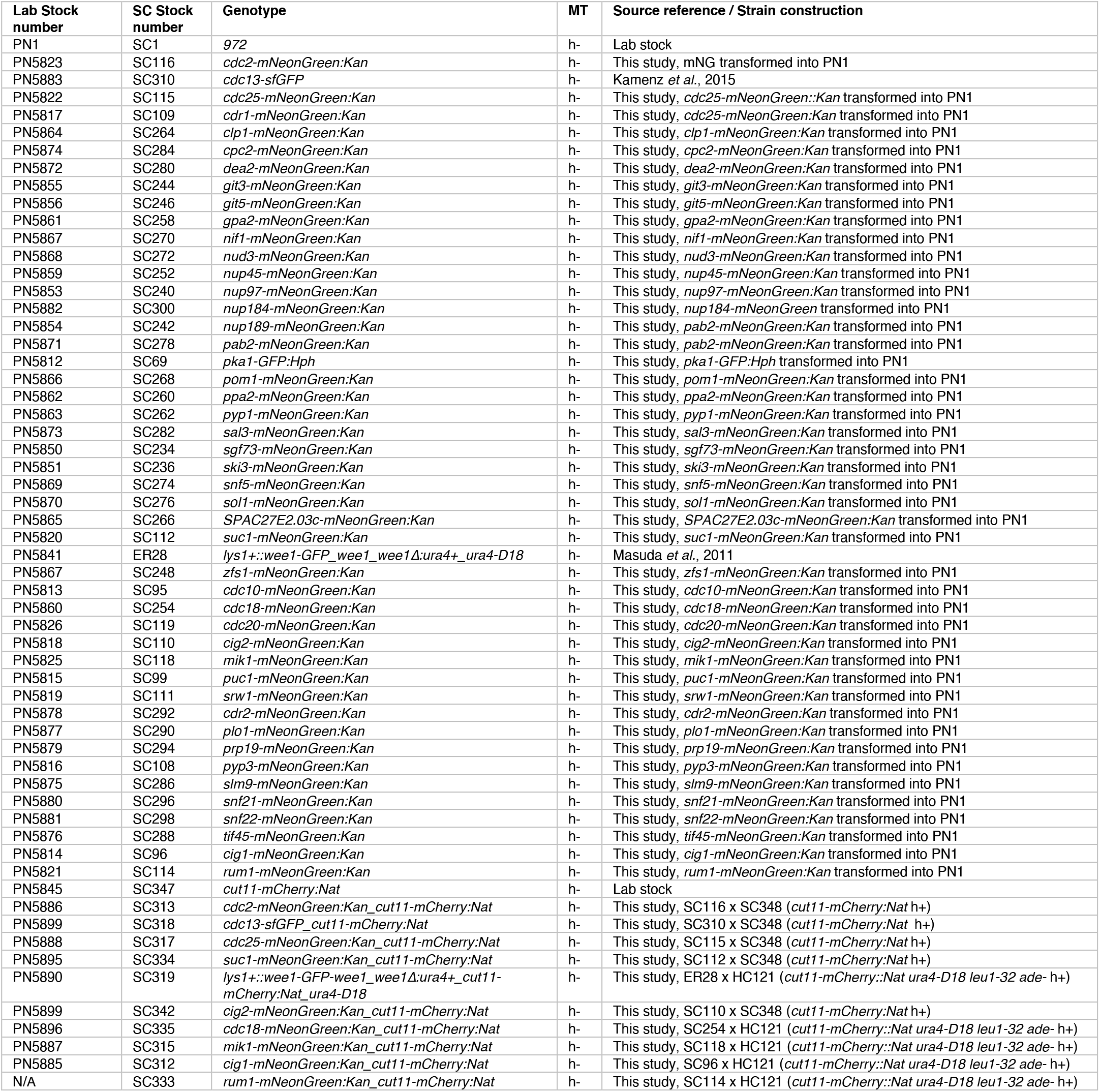
Complete list of *S. pombe* strains used in this study. List of all the *Schizosaccharomyces pombe* strains used in this study along with their full genotypes, mating type, Nurse Lab stock number (PN…), S Curran stock number (SC…) and relevant source references and strain construction notes.

**Table S2.**
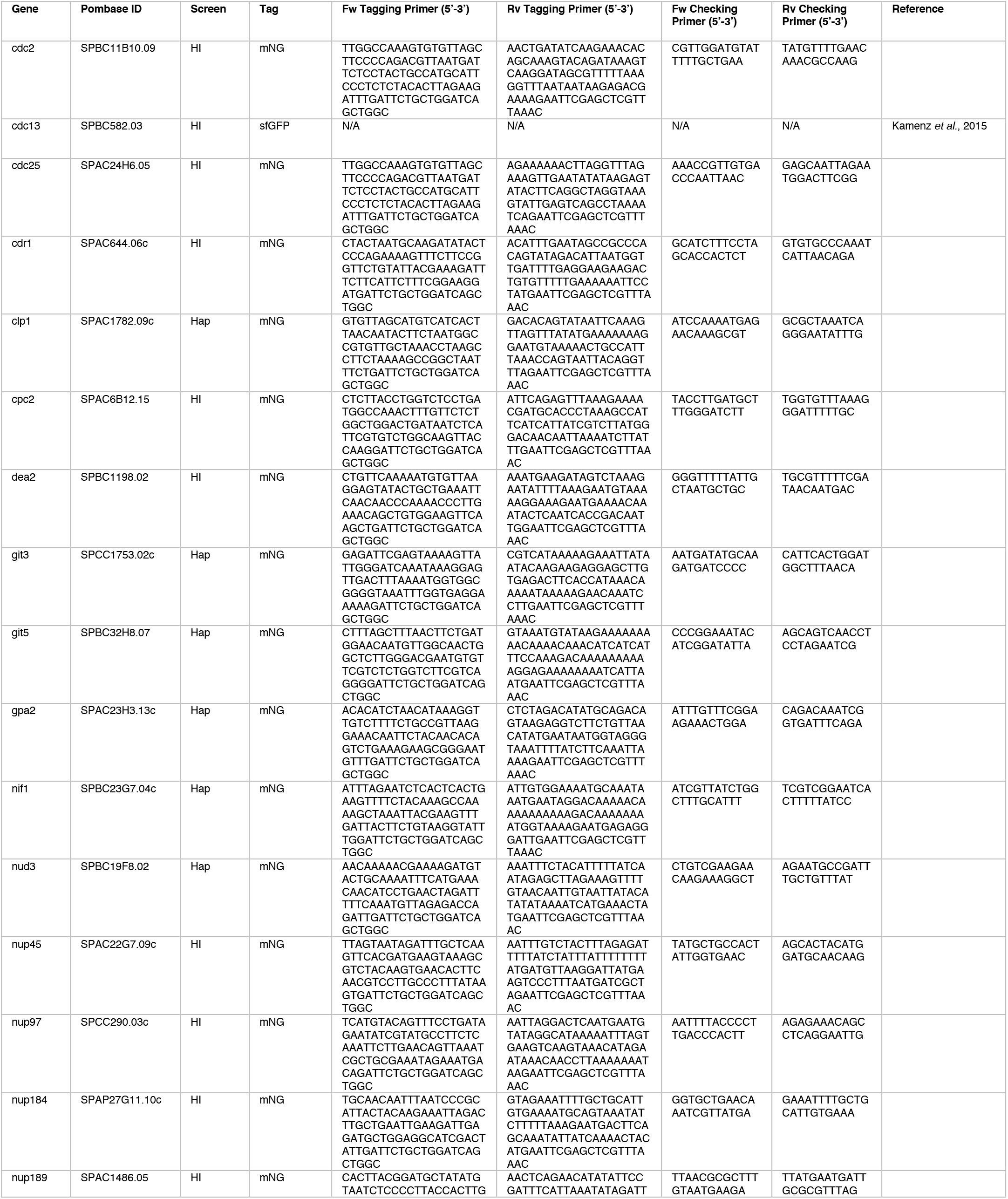
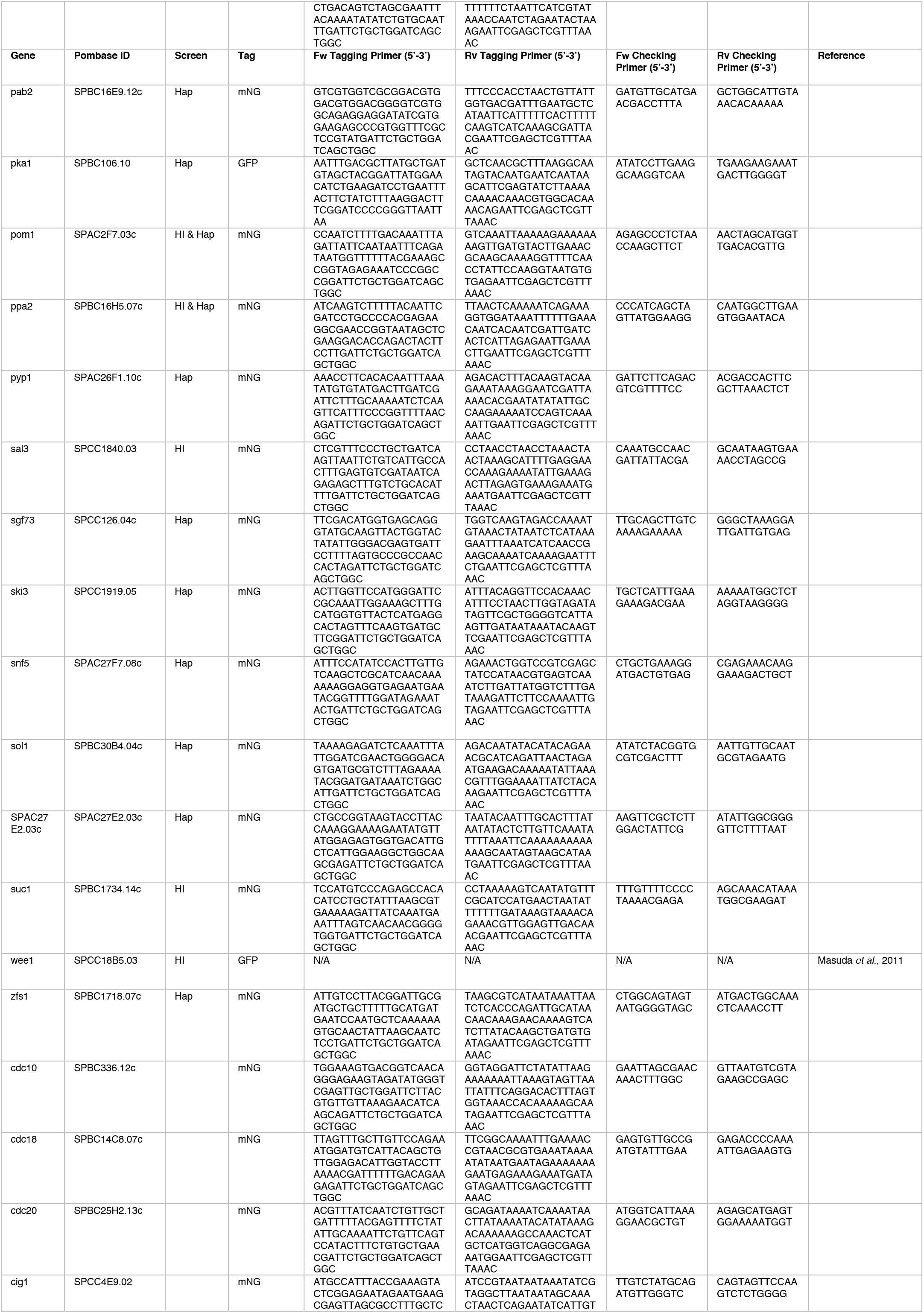
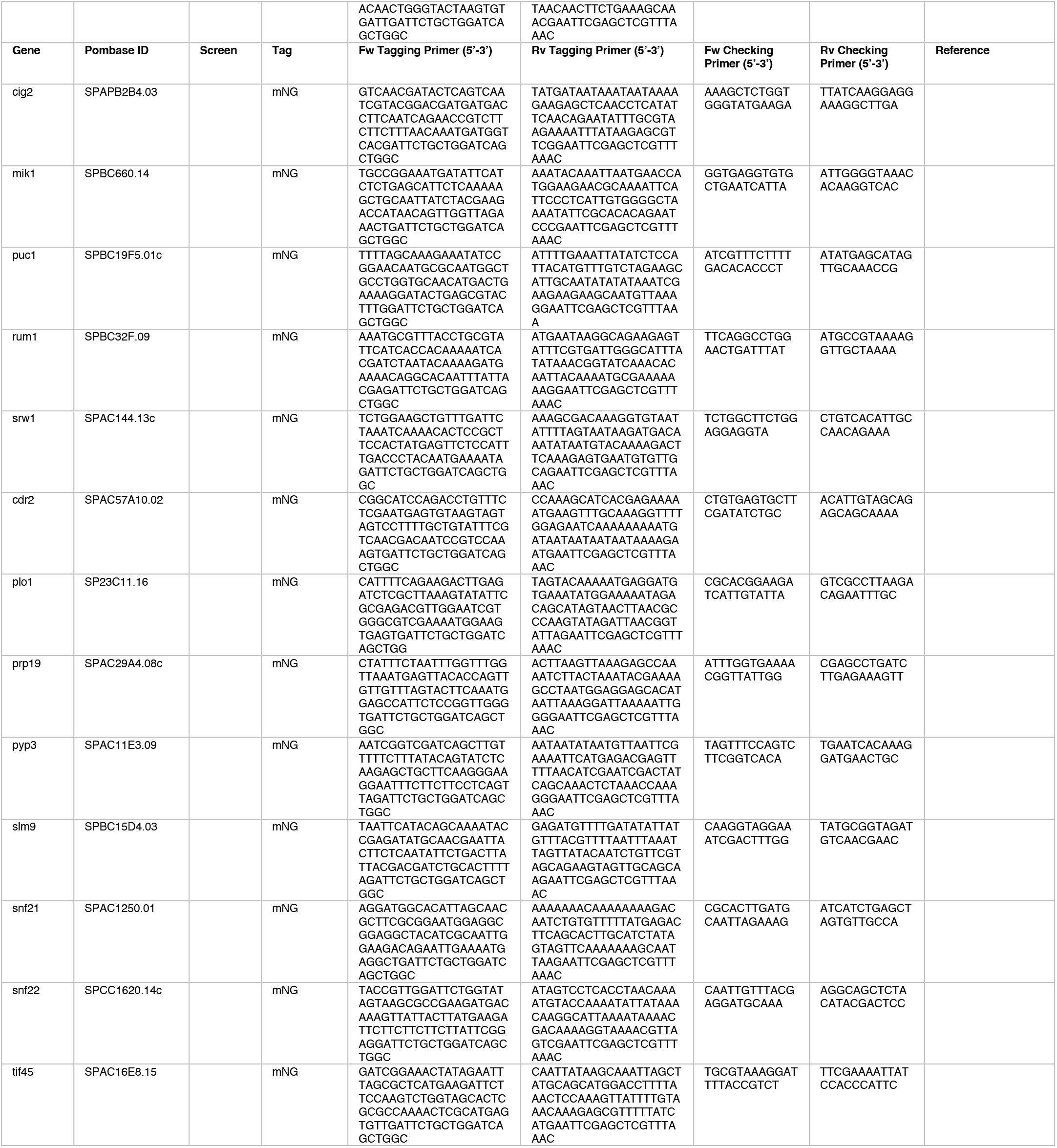
Primers used for C-terminal tagging of proteins analysed in this study. List of all *Schizosaccharomyces pombe* genes fluorescently tagged in this study, with tagging primers for pFA6A C-terminal tagging of full length proteins and checking primers.

