## Extended Supplement for "A quantitative and spatial analysis of cell cycle regulators during the fission yeast cycle"

| Page | Gene | Tag | Systematic ID | Localisation | Key |
|---|---|---|---|---|---|
| 1 | cdc2 | mNG | SPBC11B10.09 | Cytoplasm, Nucleus (increases through G2), SPB, Spindle | ◊◉✭☐▲ |
| 2 | cdc13 | sfGFP | SPBC582.03 | Nucleus (increases through G2), SPB, Spindle | ◊◉✭☐▲ |
| 3 | cdc25 | mNG | SPAC24H6.05 | Cytoplasm, Nucleus (decreases early G2, increases mid-late G2) | ◊◉✭☐ |
| 4 | cdr1 | mNG | SPAC644.06c | Cytoplasm, Medial cortical nodes | |
| 5 | clp1 | mNG | SPAC1782.09c | Nucleolus, SPB, Spindle | ◊◉✭▲ |
| 6 | cpc2 | mNG | SPAC6B12.15 | Cytoplasm | |
| 7 | dea2 | mNG | SPBC1198.02 | Cytoplasm, Nucleus | ◊◉ |
| 8 | git3 | mNG | SPCC1753.02c | Cytoplasm, Cortical Membrane, Septum | |
| 9 | git5 | mNG | SPBC32H8.07 | Cytoplasm, Cortical Membrane, Septum | |
| 10 | gpa2 | mNG | SPAC23H3.13c | Cytoplasm, Cortical Membrane, Septum, Nucleus | ◊◉ |
| 11 | nif1 | mNG | SPBC23G7.04c | Cytoplasm, Septum, Cell tips | |
| 12 | nud3 | mNG | SPBC19F8.02 | Cytoplasm, Nucleus | ◊ |
| 13 | nup45 | mNG | SPAC22G7.09c | Nucleus, Nuclear periphery, Nuclear bridge | ◊◉ |
| 14 | nup97 | mNG | SPCC290.03c | Nucleus, Nuclear periphery, Nuclear bridge | ◊◉ |
| 15 | nup184 | mNG | SPAP27G11.10c | Nucleus, Nuclear periphery, Nuclear bridge | ◊◉ |
| 16 | nup189 | mNG | SPAC1486.05 | Nucleus, Nuclear periphery, Nuclear bridge | ◊◉ |
| 17 | pab2 | mNG | SPBC16E9.12c | Cytoplasm, Nucleus (puncta), Nuclear puncta | ◊◉ |
| 18 | pka1 | GFP | SPBC106.10 | Cytoplasm, Nucleus | ◊◉ |
| 19 | pom1 | mNG | SPAC2F7.03c | Cell tips, Septum, Cytoplasm | |
| 20 | ppa2 | mNG | SPBC16H5.07c | Cytoplasm, Nucleus | ◊◉ |
| 21 | pyp1 | mNG | SPAC26F1.10c | Cytoplasm, Nucleus | ◊ |
| 22 | sal3 | mNG | SPCC1840.03 | Cytoplasm, Nucleus | ◊◉ |
| 23 | sgf73 | mNG | SPCC126.04c | Cytoplasm, Nucleus | ◊◉ |
| 24 | ski3 | mNG | SPCC1919.05 | Cytoplasm | |
| 25 | snf5 | mNG | SPAC27F7.08c | Nucleus | ◊◉ |
| 26 | sol1 | mNG | SPBC30B4.04c | Nucleus | ◊◉ |
| 27 | SPAC27E2.03c | mNG | SPAC27E2.03c | Cytoplasm, Nucleus | ◊ |
| 28 | suc1 | mNG | SPBC1734.14c | Cytoplasm, Nucleus (increases from mid G2), SPB, Spindle | ◊◉✭☐▲ |
| 29 | wee1 | mNG | SPCC18B5.03 | Cytoplasm, Nucleus (increases through G2) | ◊◉✭☐▲ |
| 30 | zfs1 | mNG | SPBC1718.07c | Cytoplasm | |
| 31 | cdc10 | mNG | SPBC336.12c | Cytoplasm, Nucleus (mild oscillation in mid G2) | ◊◉✭ |
| 32 | cdc18 | mNG | SPBC14C8.07c | Nucleus (transient G1 expression) | ◊◉✭ |
| 33 | cdc20 | mNG | SPBC25H2.13c | Nucleus | ◊ |
| 34 | cig1 | mNG | SPCC4E9.02 | SPB, Nucleus in anaphase | ◊◉✭▲ |
| 35 | cig2 | mNG | SPAPB2B4.03 | Cytoplasm, Nucleus (Decreases through G2), SPB | ◊◉✭▲ |
| 36 | mik1 | mNG | SPBC660.14 | Nucleus (Transient G1 expression) | ◊◉✭ |
| 37 | puc1 | mNG | SPBC19F5.01c | Cytoplasm, Nucleus | ◊ |
| 38 | rum1 | mNG | SPBC32F.09 | N/A, condition dependent expression | |
| 39 | srw1 | mNG | SPAC144.13c | Cytoplasm, Nucleus (decreases through G2) | ◊◉✭ |
| 40 | cdr2 | mNG | SPA57A10.02 | Cytoplasm, Medial cortical nodes | |
| 41 | plo1 | mNG | SP23C11.16 | Cytoplasm, Nucleus, SPB, Septum | ◊▲ |
| 42 | prp19 | mNG | SPAC29A4.08c | Nucleus | ◊◉ |
| 43 | pyp3 | mNG | SPAC11E3.09 | Cytoplasm, Nucleus | ◊ |
| 44 | slm9 | mNG | SPBC15D4.03 | Nucleus | ◊◉ |
| 45 | snf21 | mNG | SPAC1250.01 | Nucleus | ◊◉ |
| 46 | snf22 | mNG | SPCC.1620.14c | Nucleus | ◊◉ |
| 47 | tif45 | mNG | SPAC16E8.15 | Cytoplasm | |

◊ 36/47 are nuclear localised

◉ 29/36 are nuclear concentrated (higher than cytoplasm)

✭ 12/29 have changing nuclear levels

☐ 5/12 peak at G2/M transition

▲ 7/47 are SPB localised

**Figure Legend**

**Information**

- Gene standard name and fluorophore tag
- Nurse lab strain number
- Pombase Systematic ID
- Full product name according to Pombase

**Images**

- Left, Single slice (focal plane) representative brightfield image of asynchronous culture.
- Right, Maximum intensity projection of fluorescent imaging (green).

**Plots from single fluorophore imaging widefield imaging**

- Blue, Mean whole cell fluorescence intensity vs cell length
- Red, Mean of top % of pixels (stated on Y-axis for each strain) vs cell length
- Teal, Total fluorescence intensity vs cell length.

Coloured dots represent individual cell values. Circles represent mean at 0.25 $\mu$m length bins with >10 cells/bin. N numbers are stated plots. Errors bars, 95% CI. Vertical dotted line represents septation length.

**Plots of two-colour widefield imaging (where available)**

- Plots showing mean cellular fluorescence intensity (light blue), mean nuclear fluorescence for mononucleate cells (green) and binucleates (pinks) vs cell length.

Coloured dots represent individual cell values. Black lines represent connected mean values calculated at 0.5 $\mu$m bins for whole cell vales (bottom), and nuclear mononucleate values (top). Vertical dotted line indicates cell length at septation. N numbers stated on plots.

**Plots for single colour imaging flow cytometry data**

- Left, Mean whole cell fluorescence intensity vs cell length.
- Right, Max pixel fluorescence intensity vs cell length.

Black circles indicate mean of bins at 0.33 $\mu$m length intervals. Error bars = 95% CI. >500 cells/bin. N numbers stated on plots.

* Dates indicate day of experiment for future reference to aid authors and to reader for any potential request of raw data.

### cdc2-mNG

PN5823/PN5886
SPBC11B10.09
cyclin-dependent protein kinase Cdk1/Cdc2

Widefield: 19/10/18

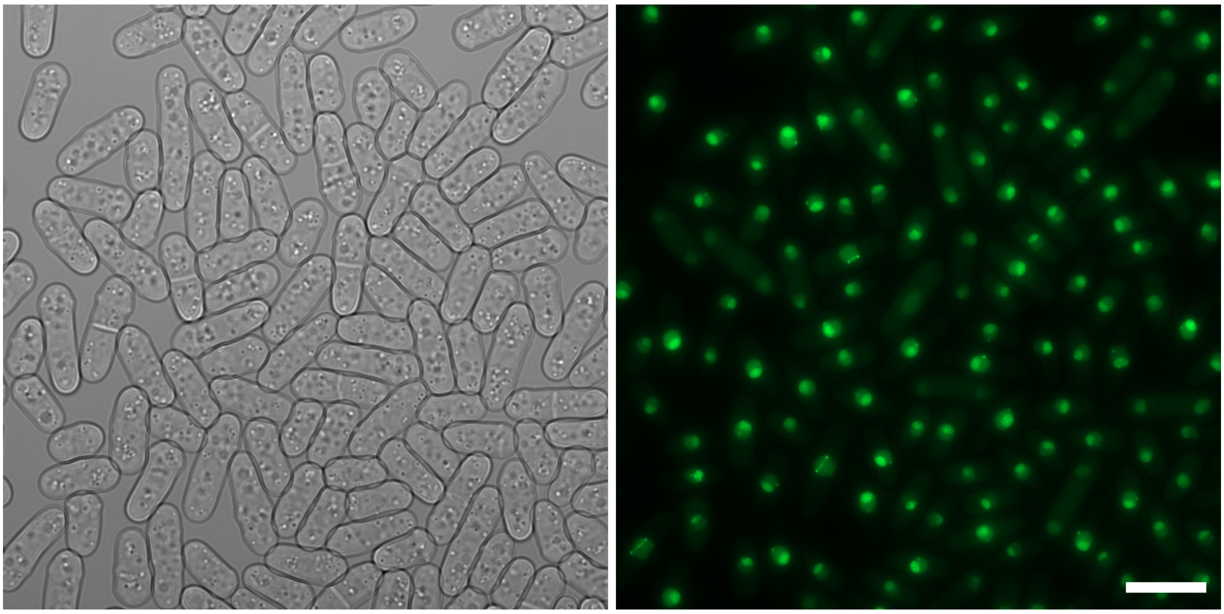

Widefield: 19/10/18

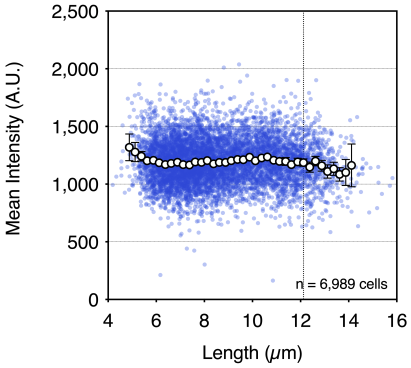


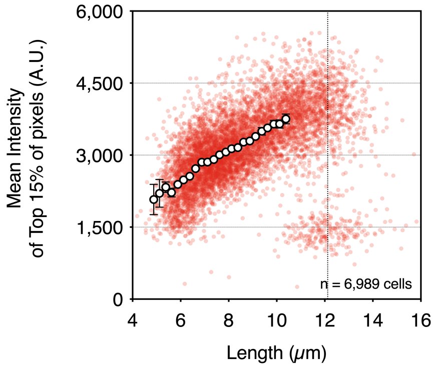


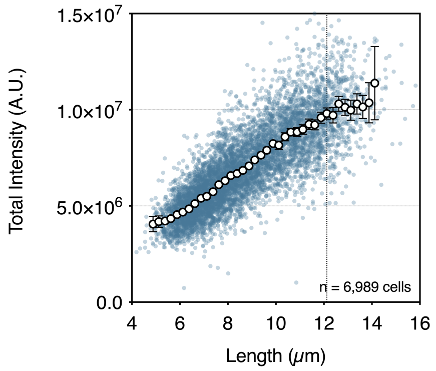


Widefield: 12/03/21

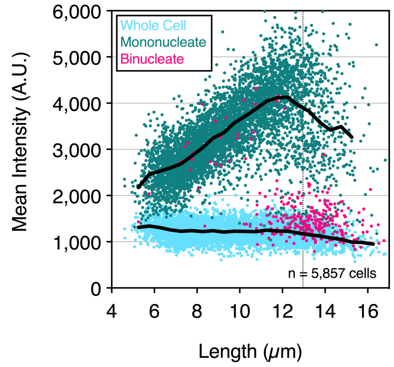


Imagestream: 10/10/18

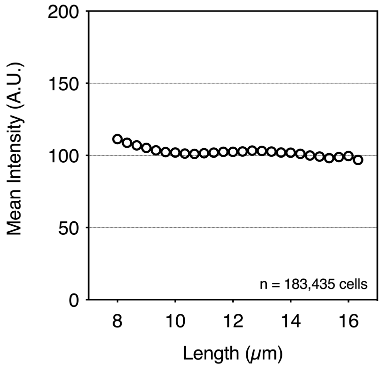


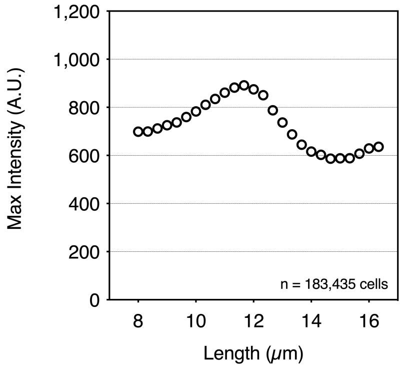

### cdc13-sfGFP

PN5883/PN5899
SPBC582.03
G2/M B-type cyclin Cdc13

Widefield: 19/10/18

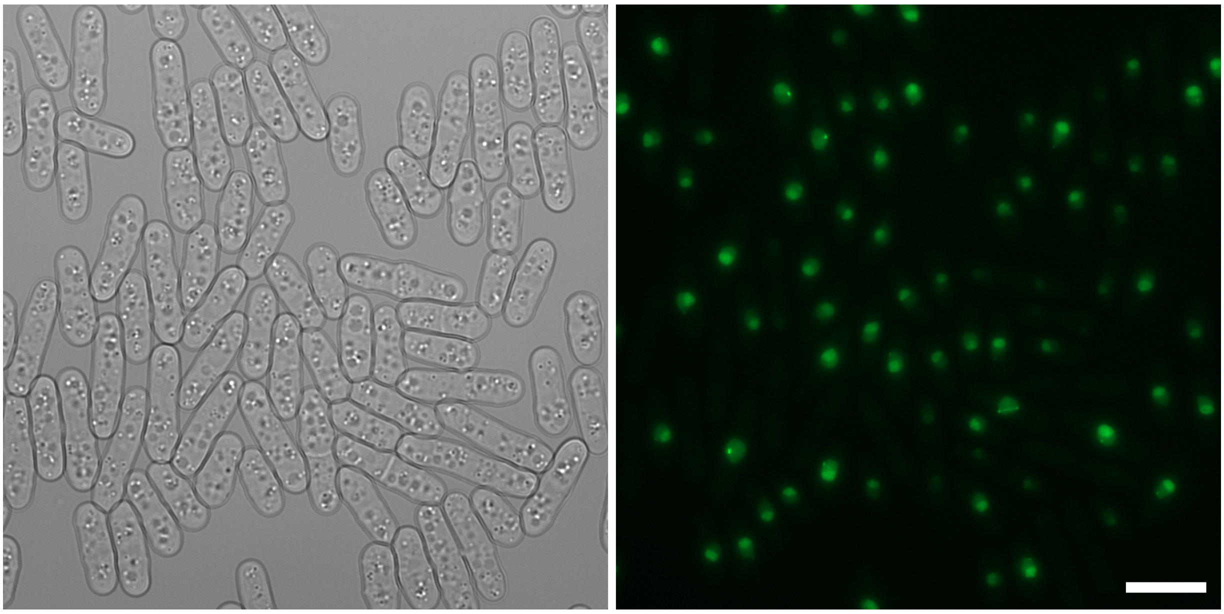

Widefield: 19/10/18

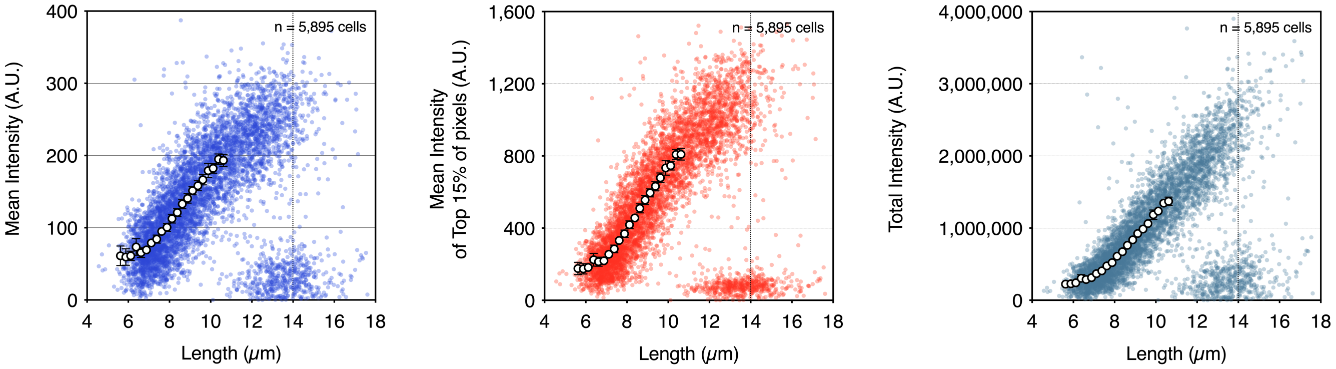


Widefield: 12/03/21

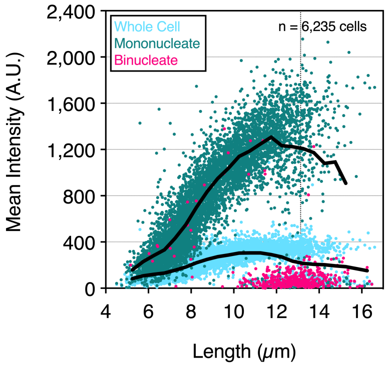


Imagestream: 10/10/18

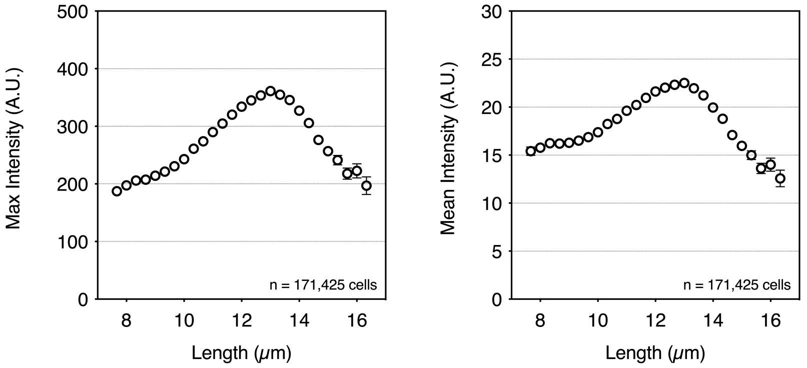

### cdc25-mNG

PN5822/PN5888
SPAC24H6.05
protein tyrosine phosphatase (M phase inducer) Cdc25

Widefield: 19/10/18

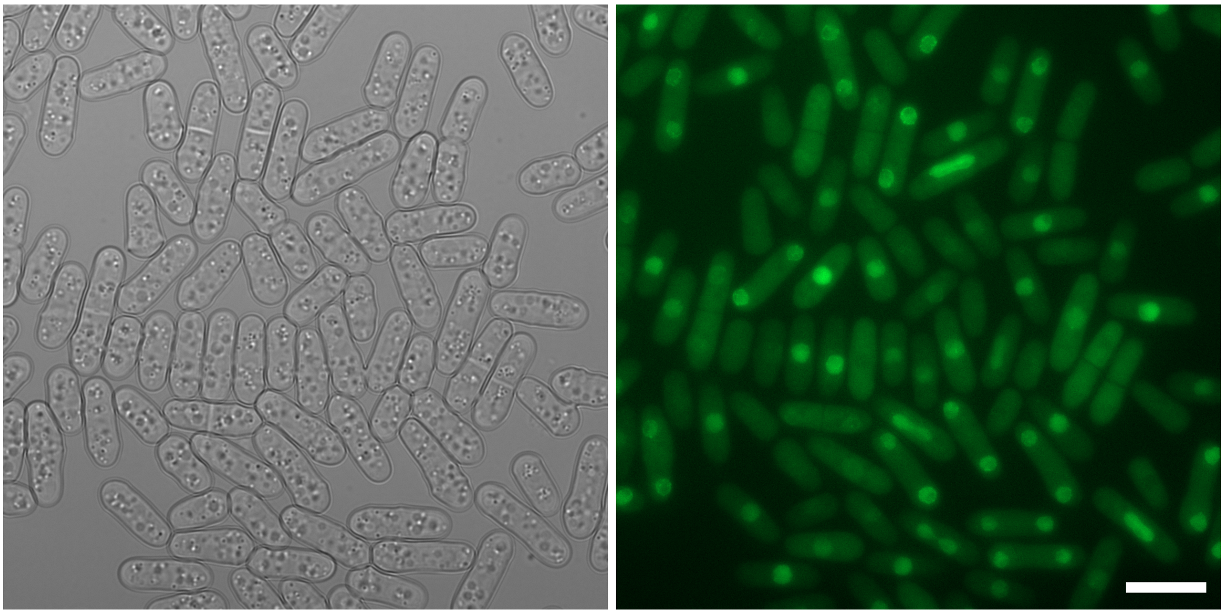

Widefield: 19/10/18

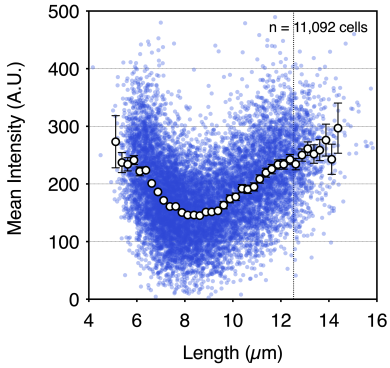


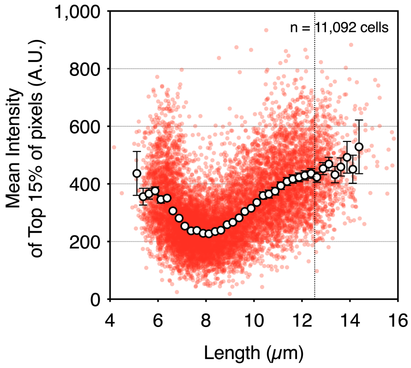


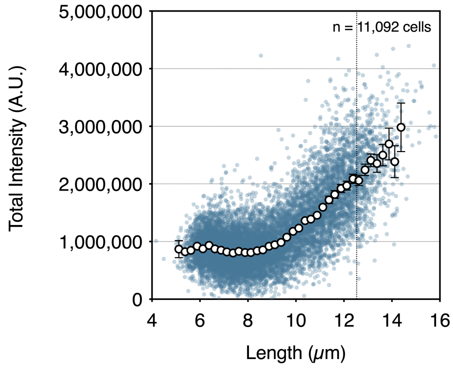


Widefield: 12/03/21

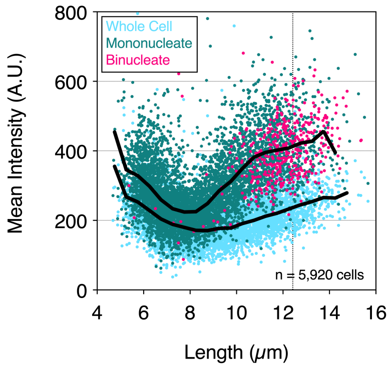


Imagestream: 10/10/18

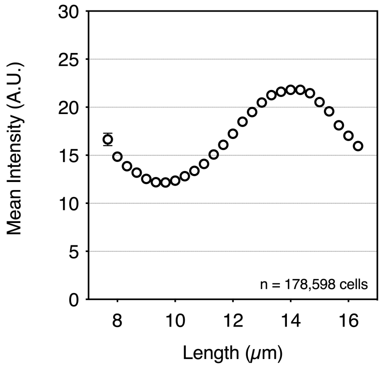


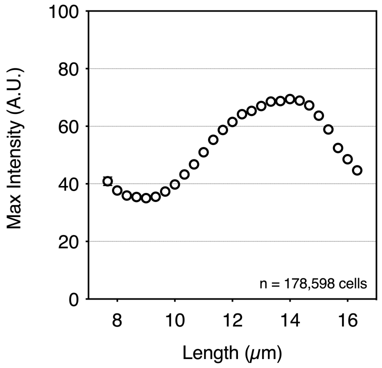

### cdr1-mNG

PN5817
SPAC644.06c
NIM1 family serine/threonine protein kinase Cdr1/Nim1

Widefield: 27/04/18

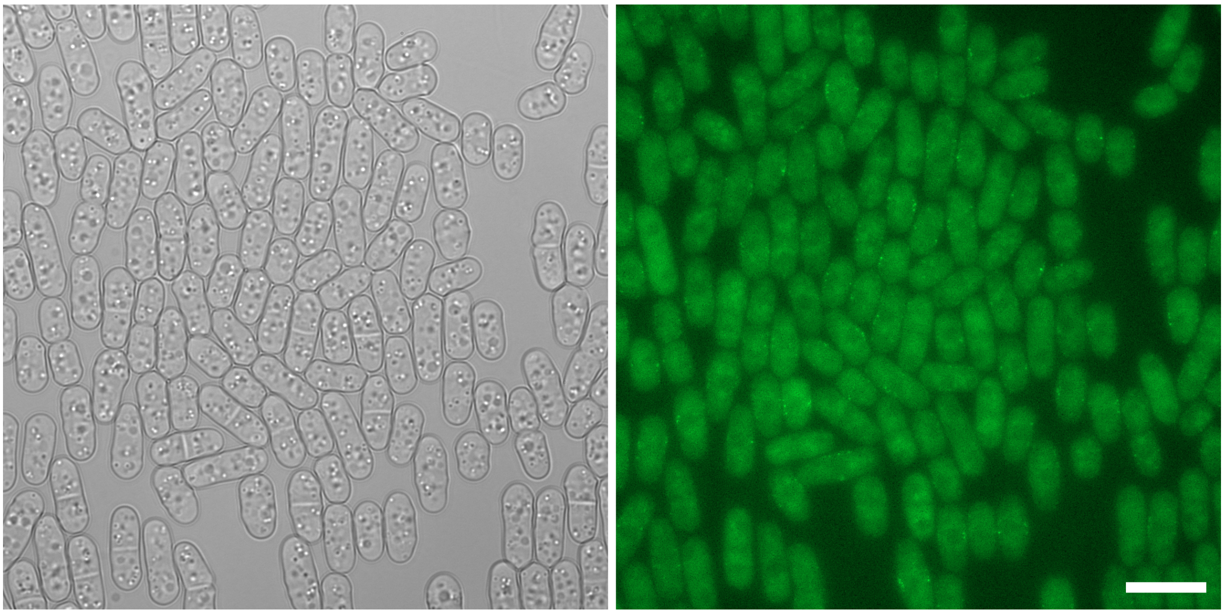

Widefield: 27/04/18

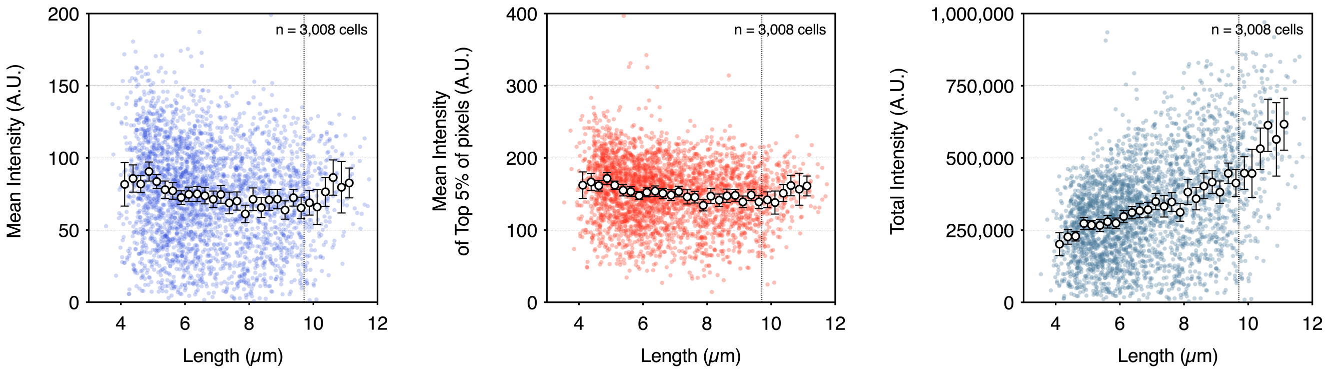


Imagestream: 09/11/18

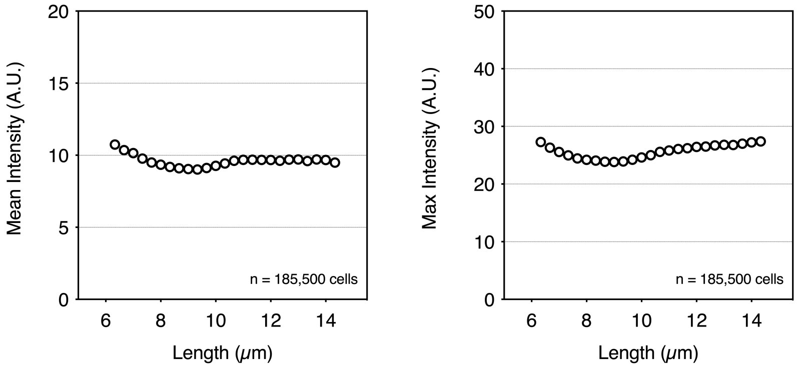

### clp1-mNG

PN5864
SPAC1782.09c
Cdc14-related protein phosphatase Clp1/Flp1

Widefield: 19/04/18

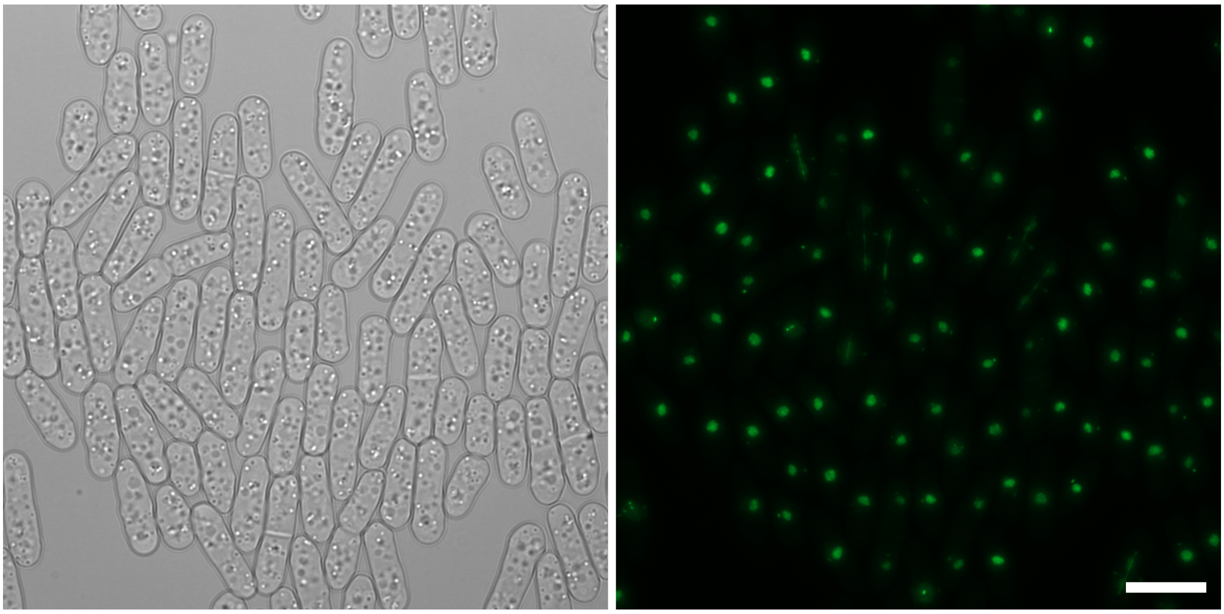

Widefield: 19/04/18

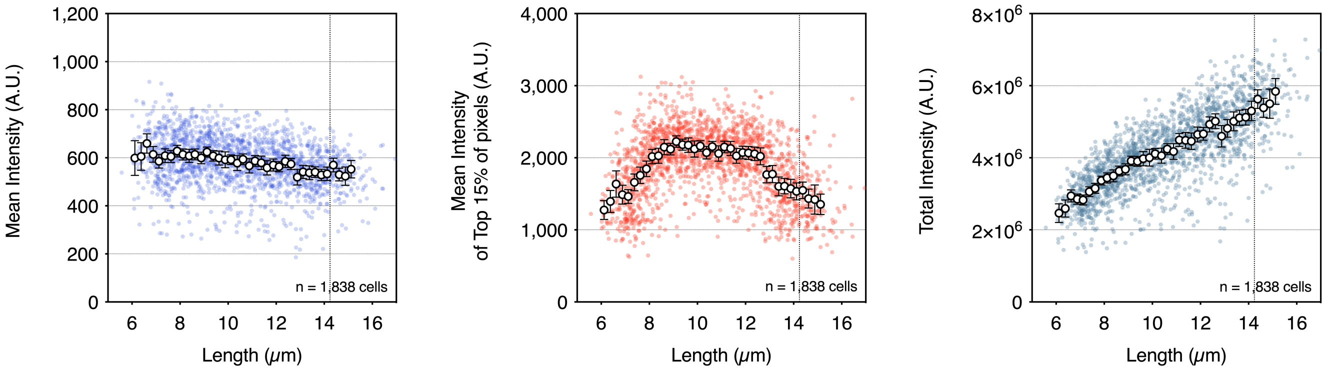


Imagestream: 17/10/18

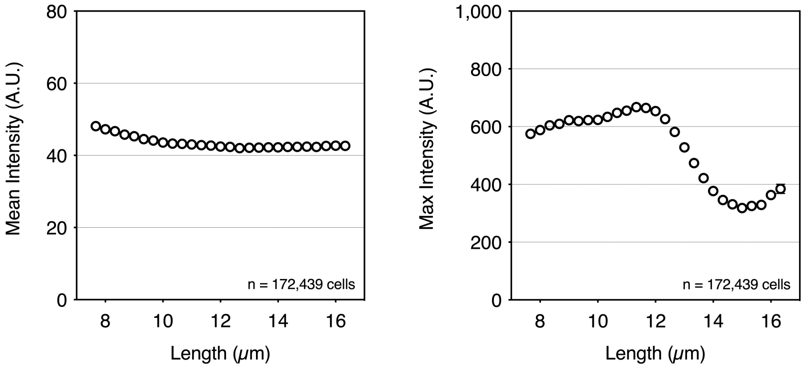

### cpc2-mNG

PN5874
SPAC6B12.15
RACK1 ortholog Cpc2

Widefield: 20/04/18

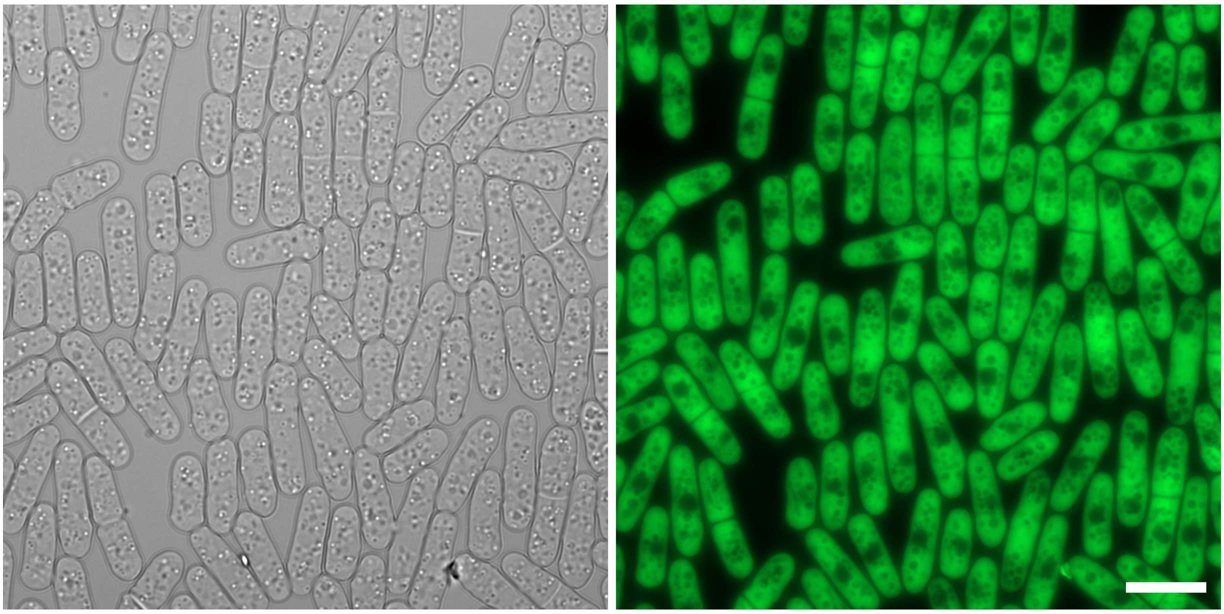

Widefield: 20/04/18

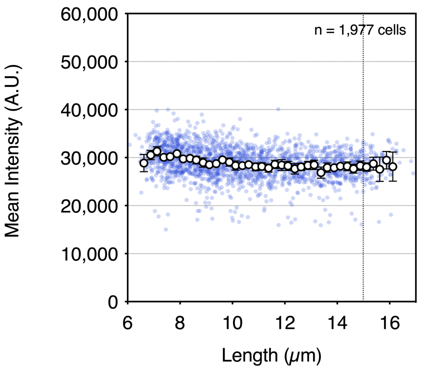


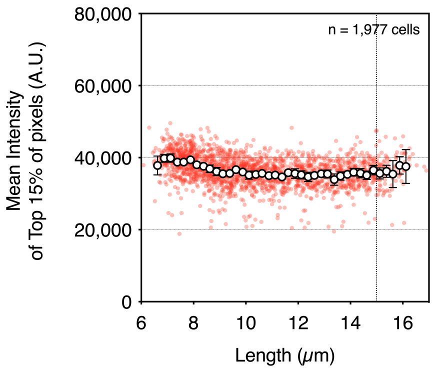


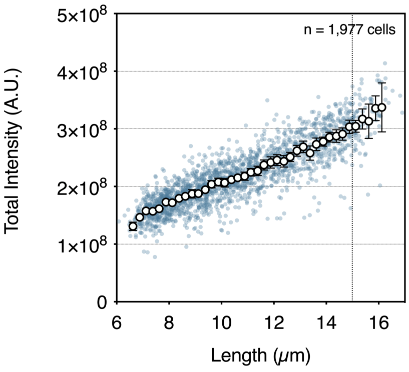


Imagestream: 10/10/18

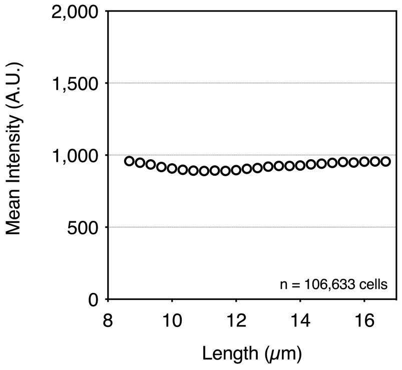


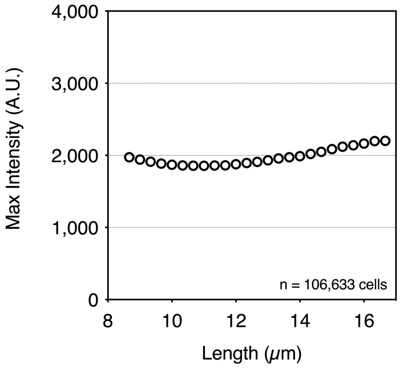

### dea2-mNG

PN5872
SPBC1198.02
adenine deaminase Dea2

Widefield: 20/04/18

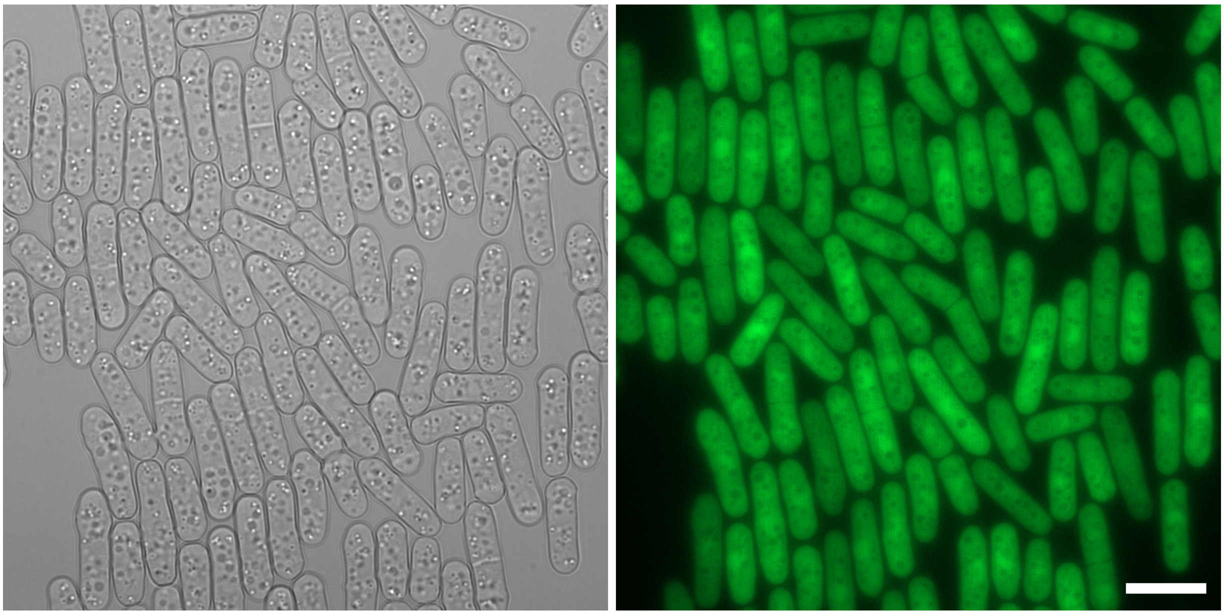

Widefield: 20/04/18

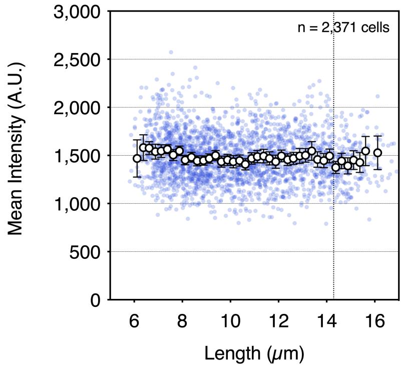


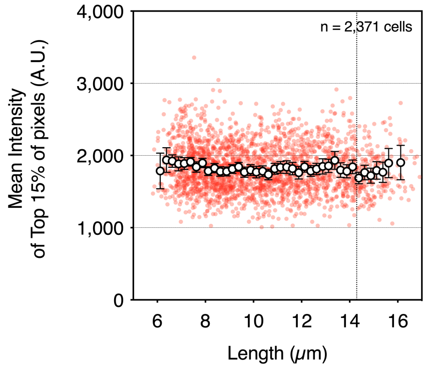


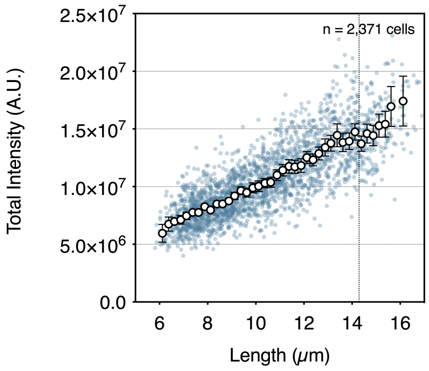


Imagestream: 12/10/18

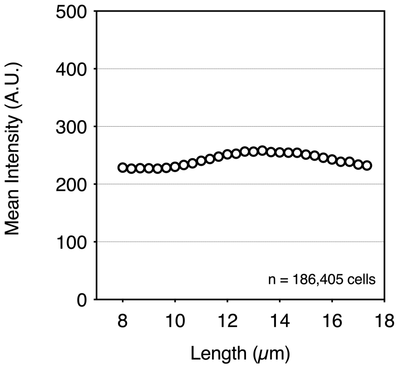


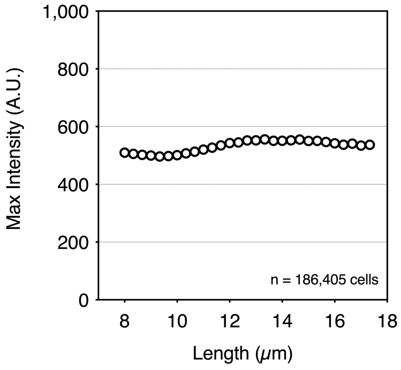

### git3-mNG

PN5855
SPCC1753.02c
G-protein coupled receptor Git3

Widefield: 02/05/18

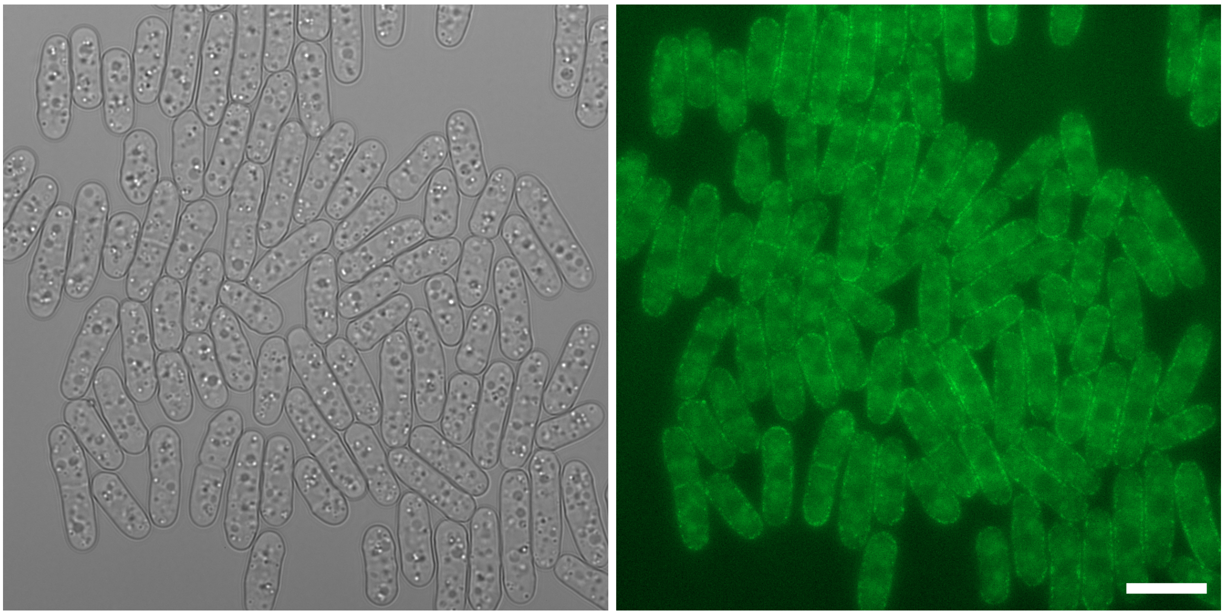

Widefield: 02/05/18

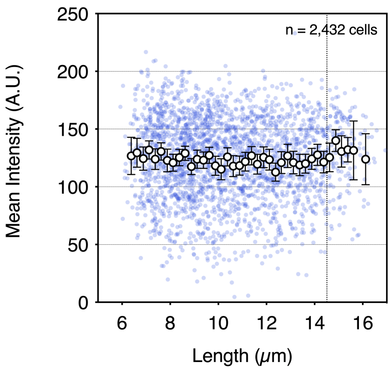


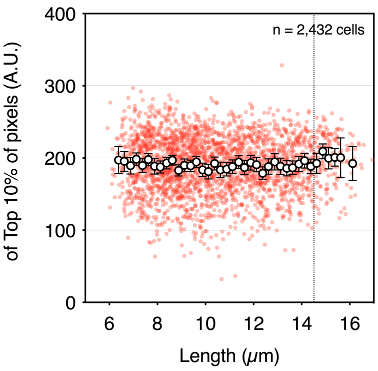


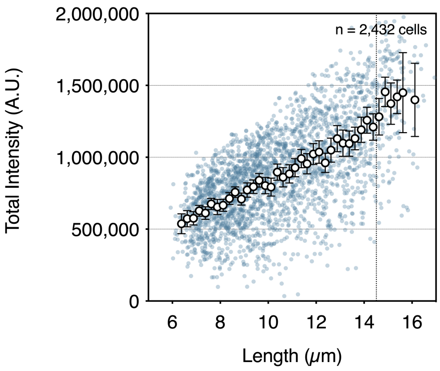


Imagestream: 10/10/18

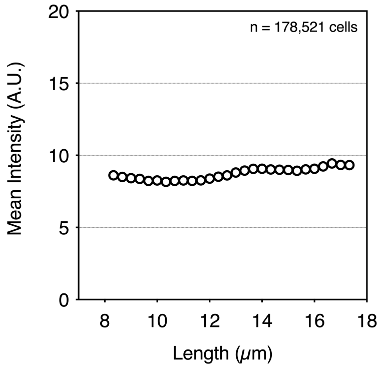


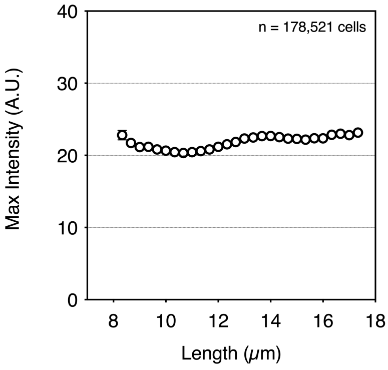

### git5-mNG

PN5856
SPBC32H8.07
heterotrimeric G protein beta (WD repeat) subunit Git5

Widefield: 27/04/18

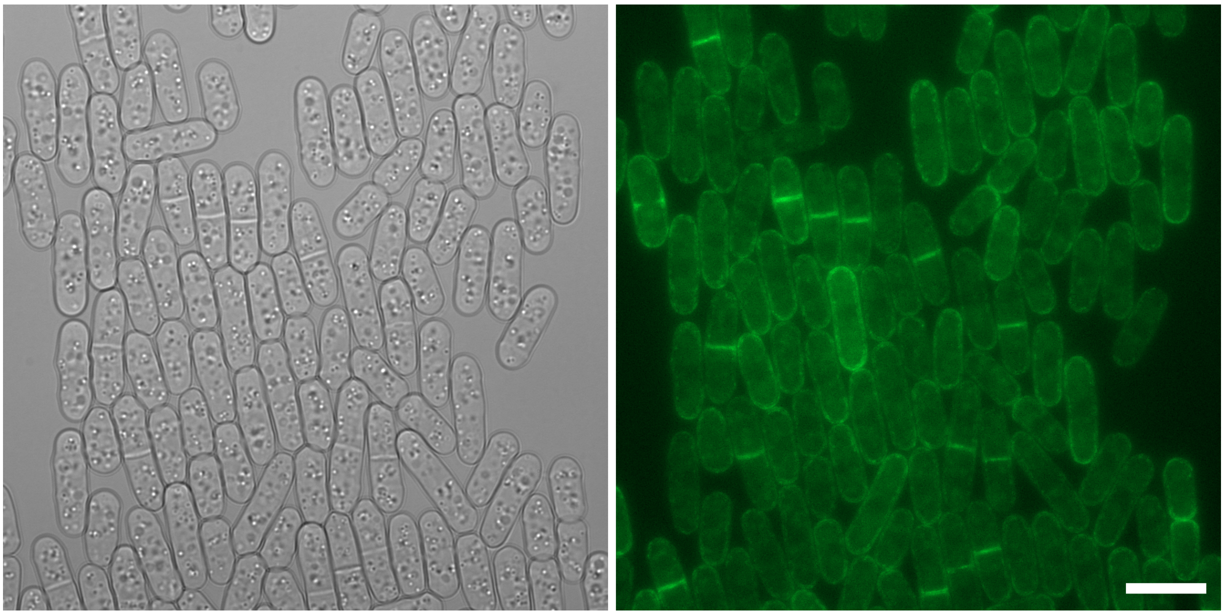

Widefield: 27/04/18

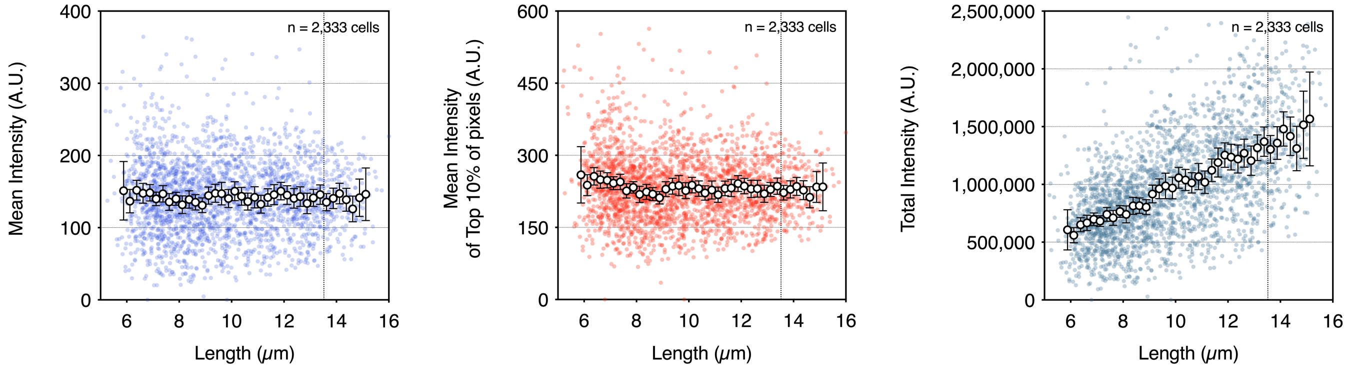


Imagestream: 11/10/18

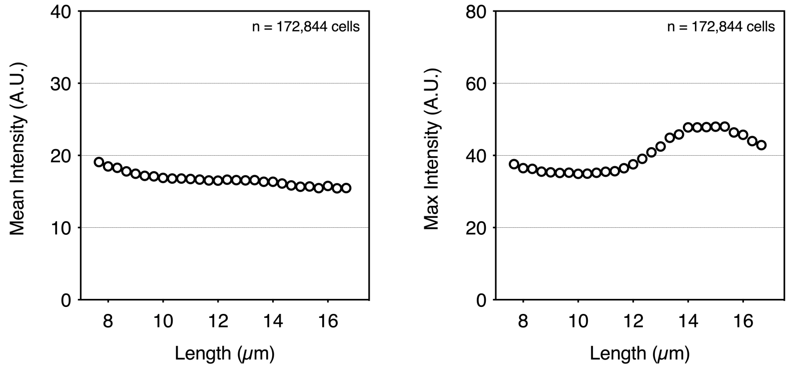

### gpa2-mNG

PN5861
SPAC23H3.13c
heterotrimeric G protein alpha-2 subunit Gpa2

Widefield: 19/10/18

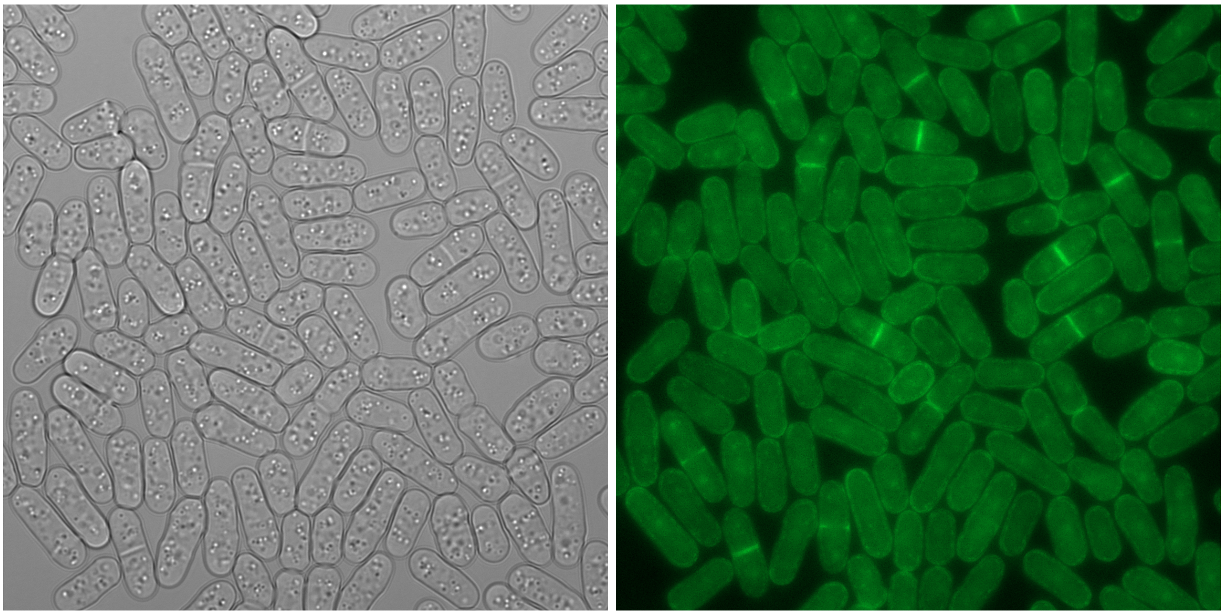

Widefield: 19/10/18

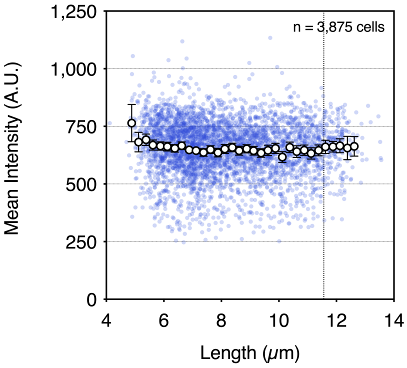


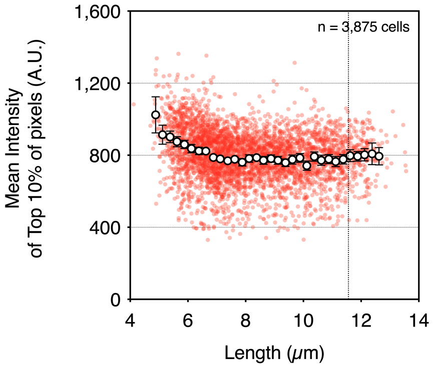


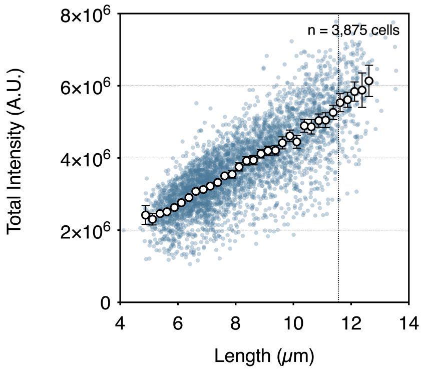


Imagestream: 12/10/18

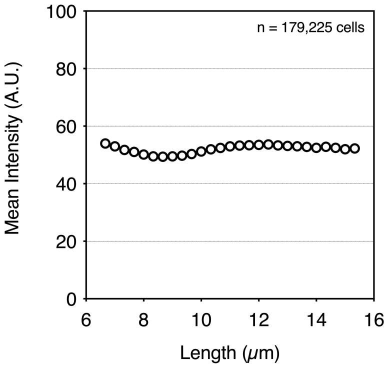


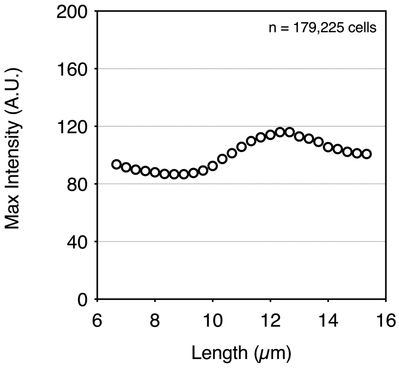

### nif1-mNG

PN5867
SPBC23G7.04c
protein kinase inhibitor, SEL1 repeat protein Nif1

Widefield: 02/05/18

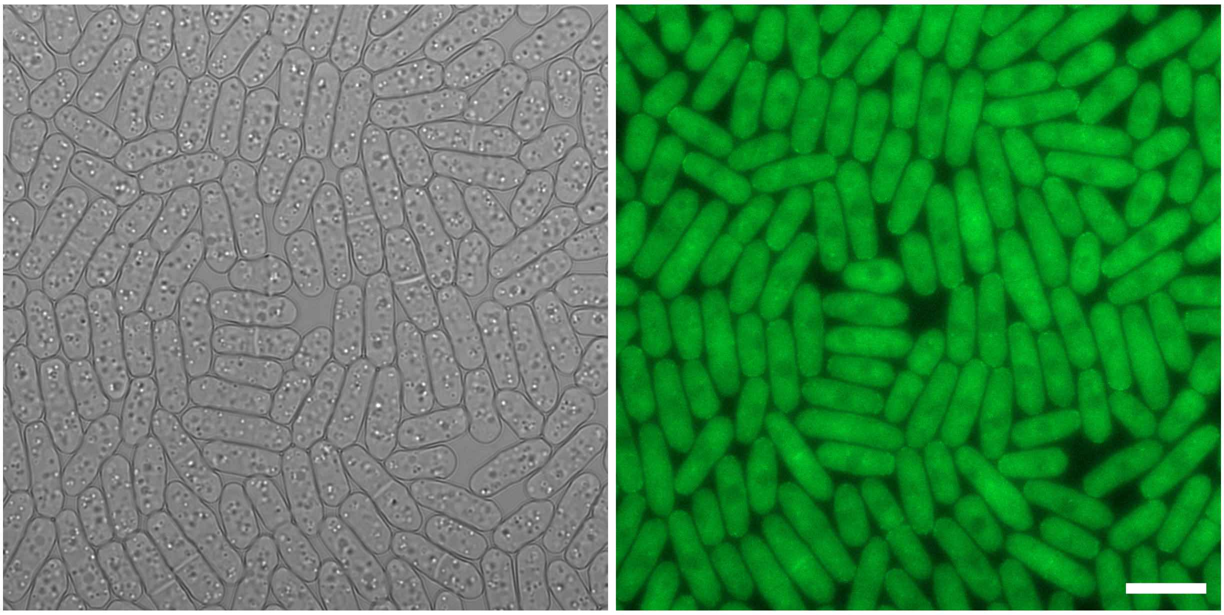

Widefield: 02/05/18

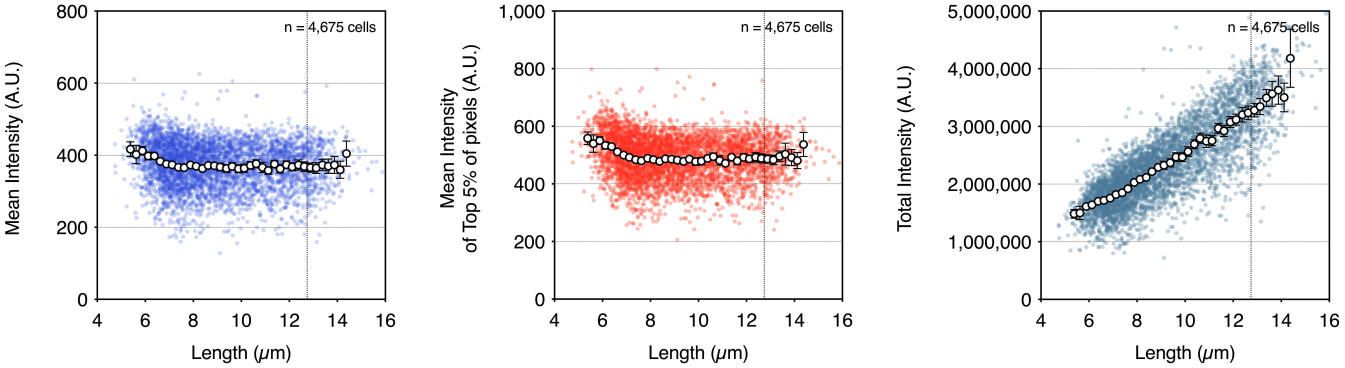


Imagestream: 12/10/18

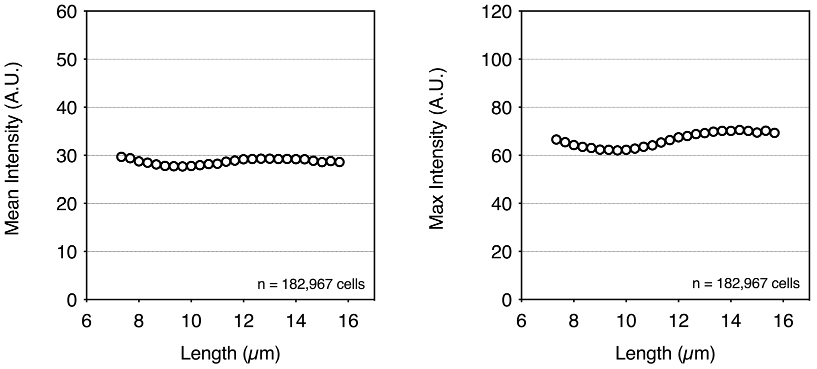

### nud3-mNG

PN5868
SPBC19F8.02
nuclear distribution protein NUDC homolog

Widefield: 27/04/18

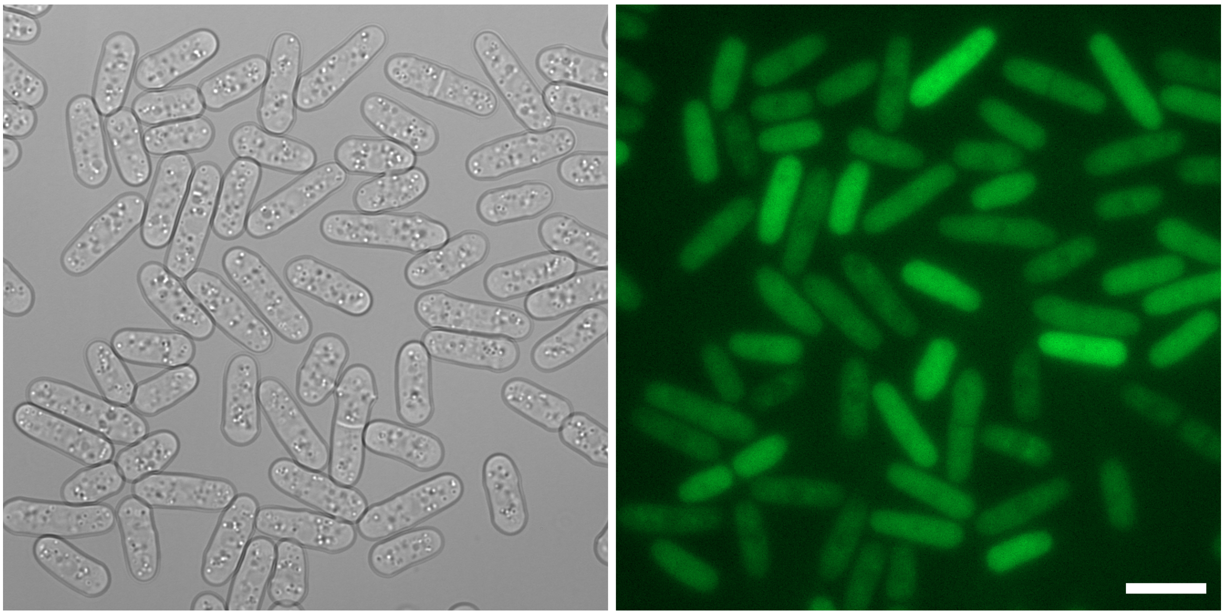

Widefield: 27/04/18

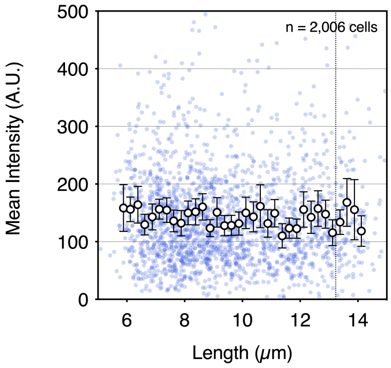


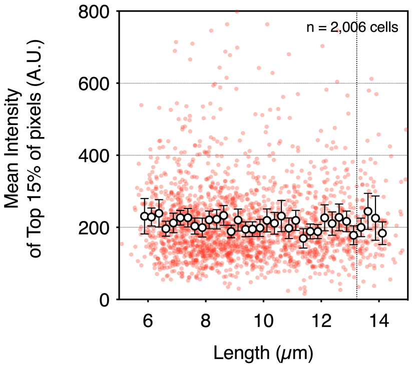


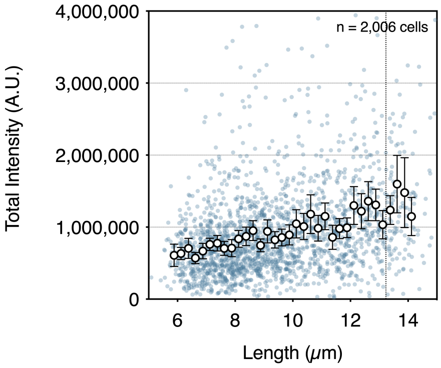


Imagestream: 10/10/18

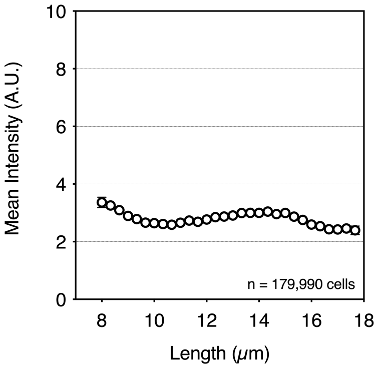


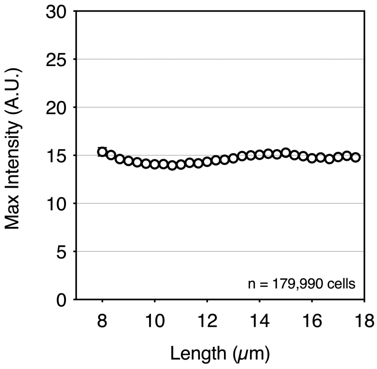

### nup45-mNG

PN5859
SPAC22G7.09c
nucleoporin Nup45

Widefield: 27/04/18

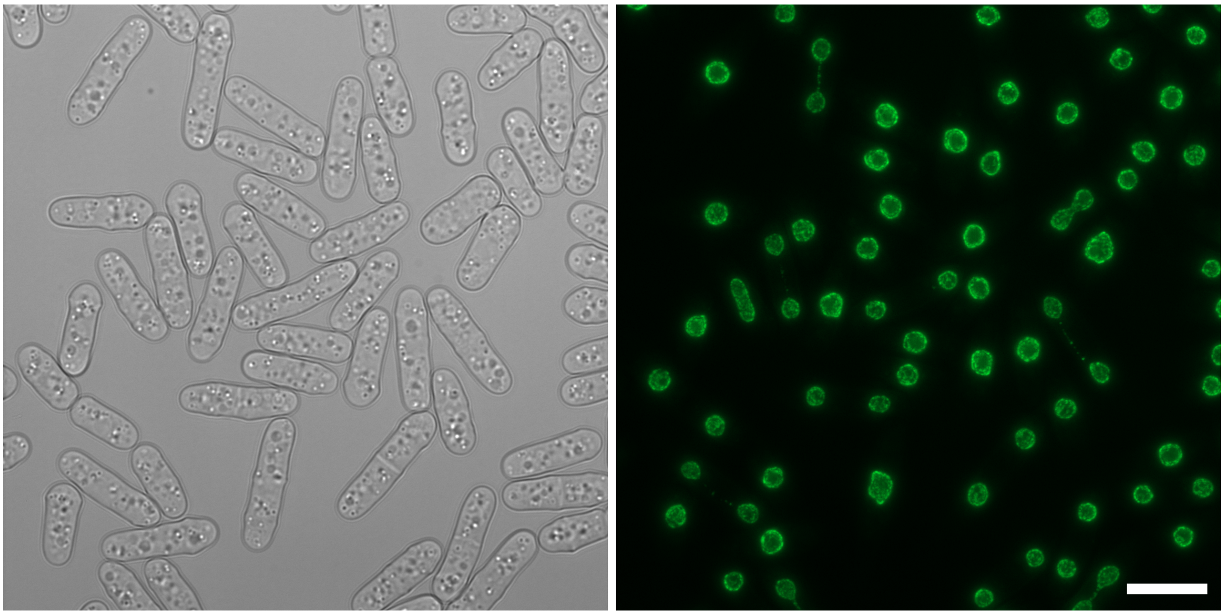

Widefield: 27/04/18

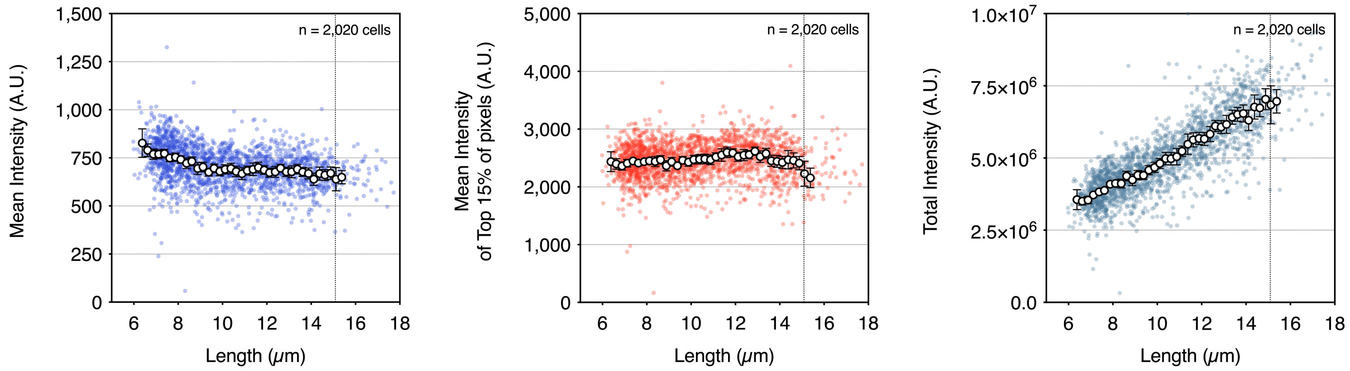


Imagestream: 10/10/18

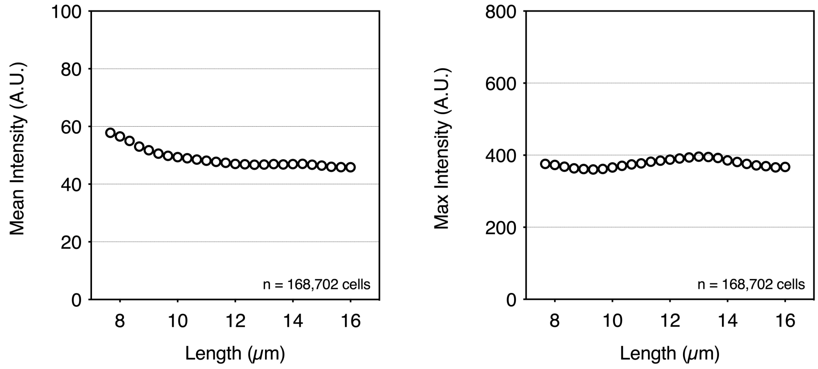

### nup97-mNG

PN5853
SPCC290.03c
nucleoporin Nic96 homology

Widefield: 27/04/18

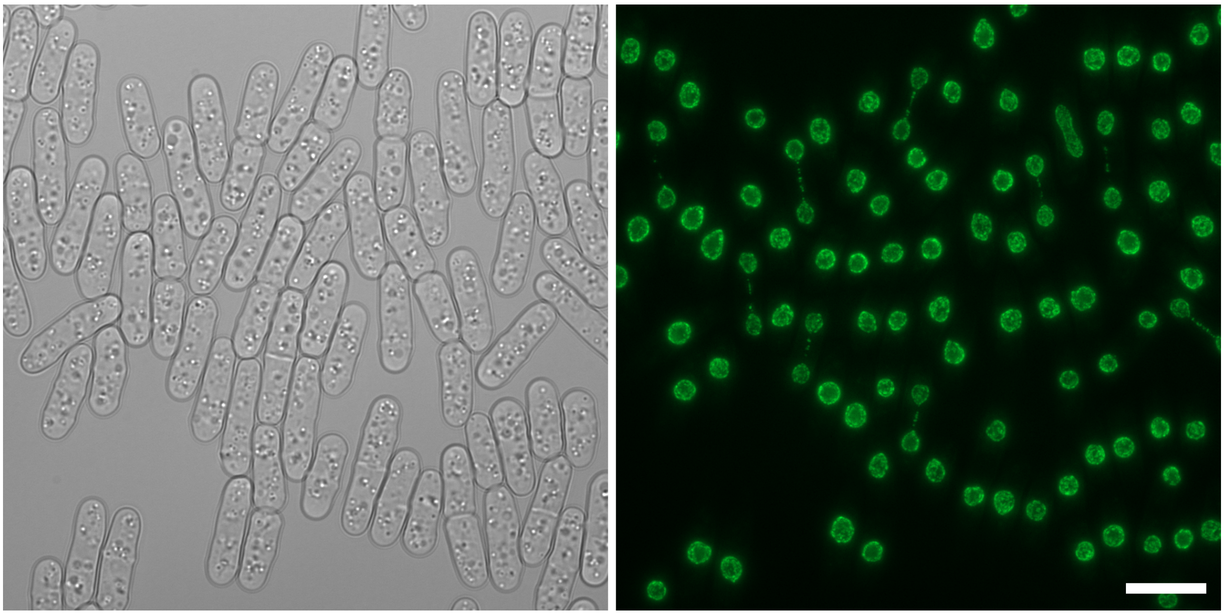

Widefield: 27/04/18

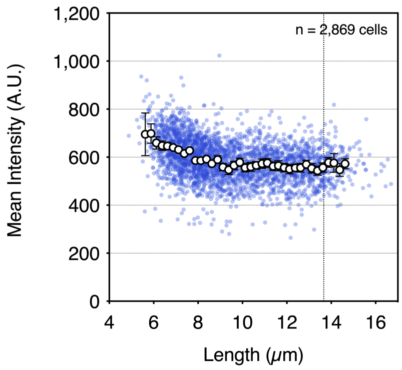


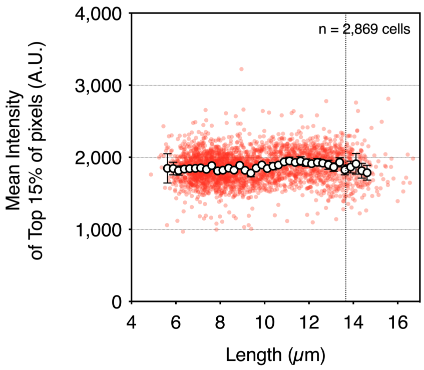


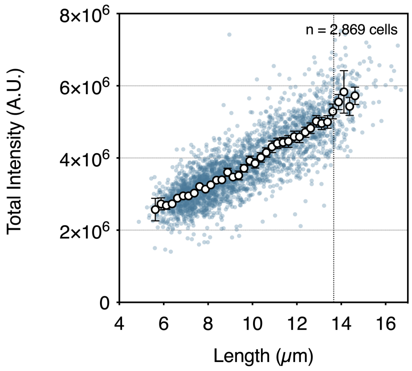


Imagestream: 12/10/18

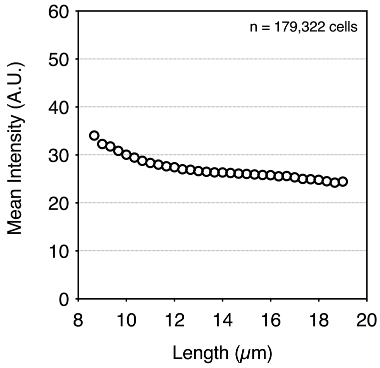


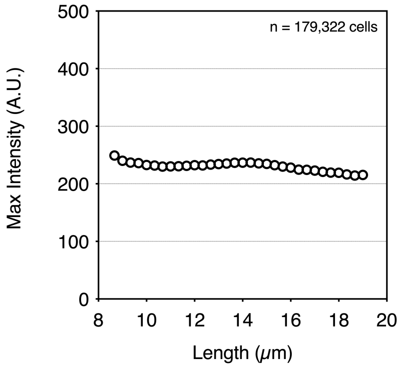

### nup184-mNG

PN5882
SPAP27G11.10c
nucleoporin Nup184

Widefield: 19/04/18

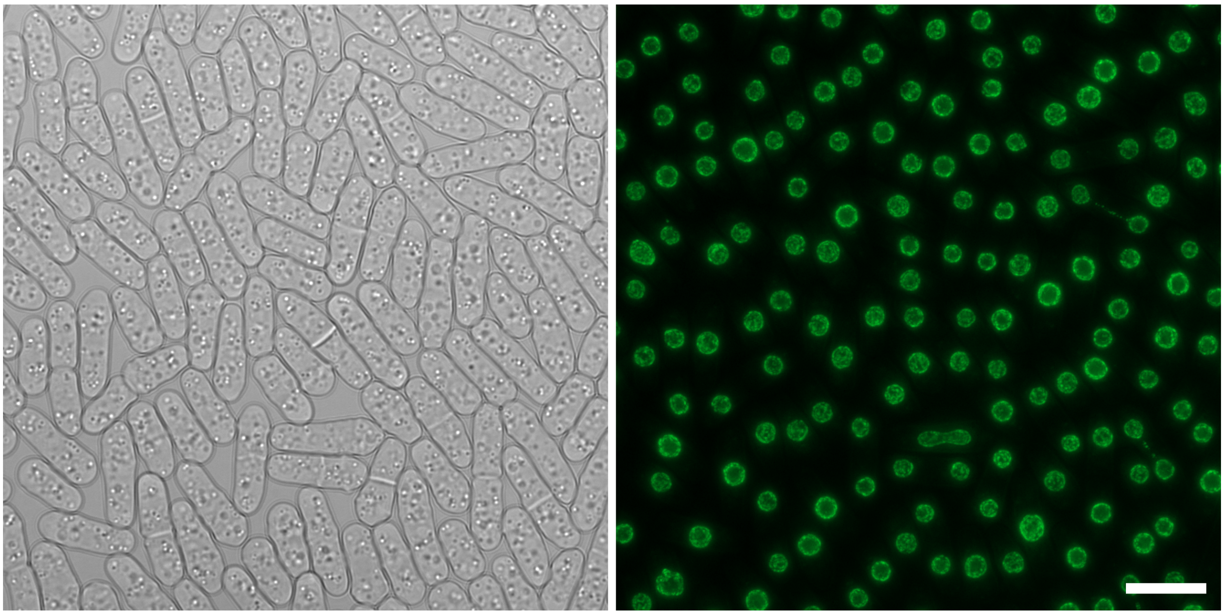

Widefield: 19/04/18

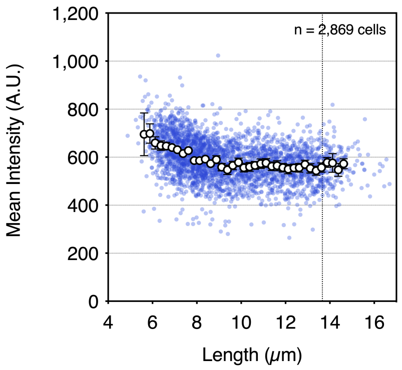


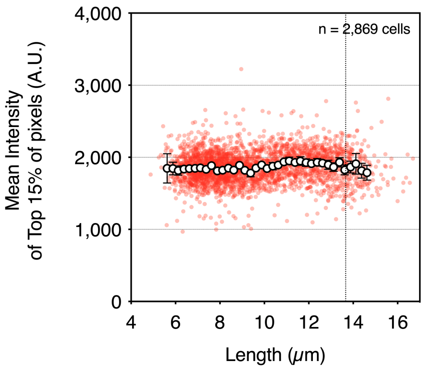


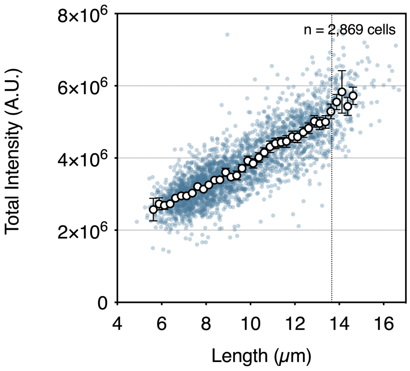


Imagestream: 12/10/18

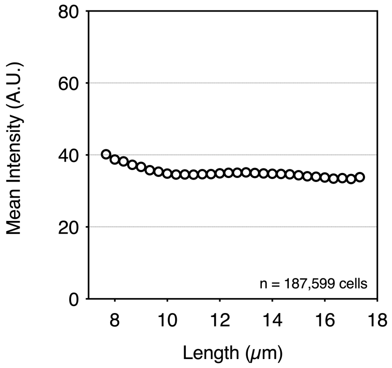


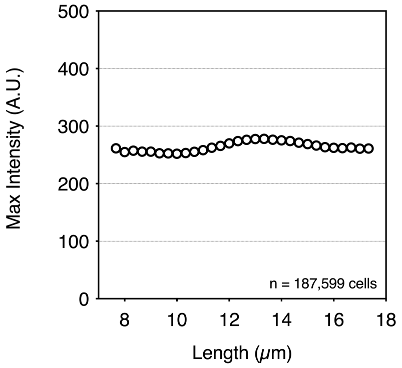

### nup189-mNG

PN5854
SPAC1486.05
nucleoporin Nup98 and Nup96

Widefield: 27/04/18

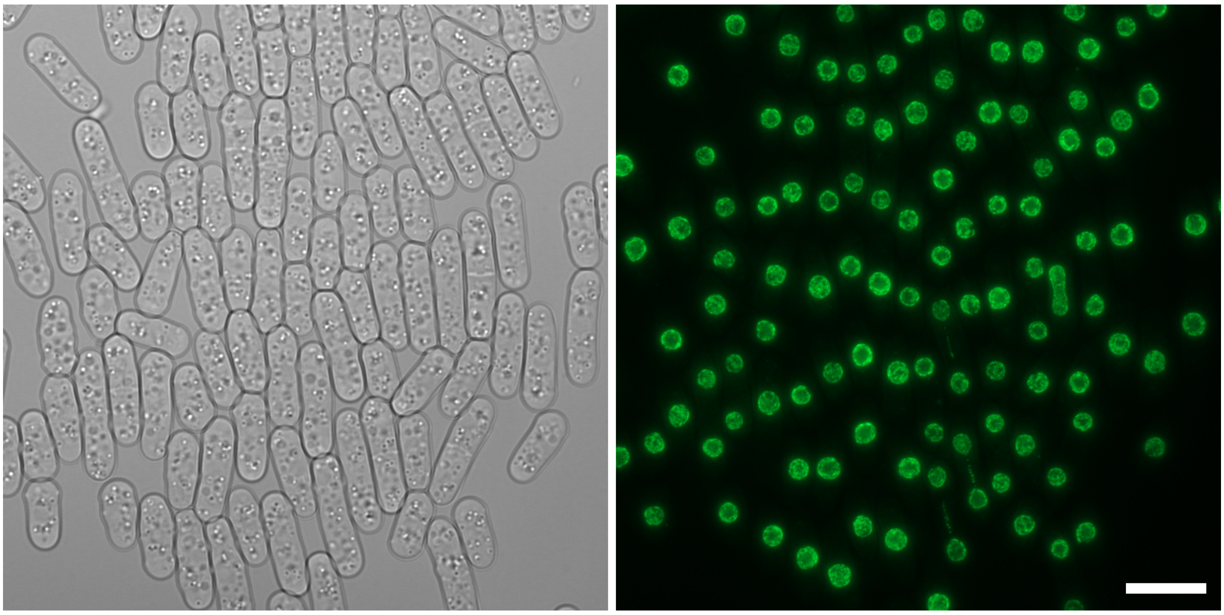

Widefield: 27/04/18

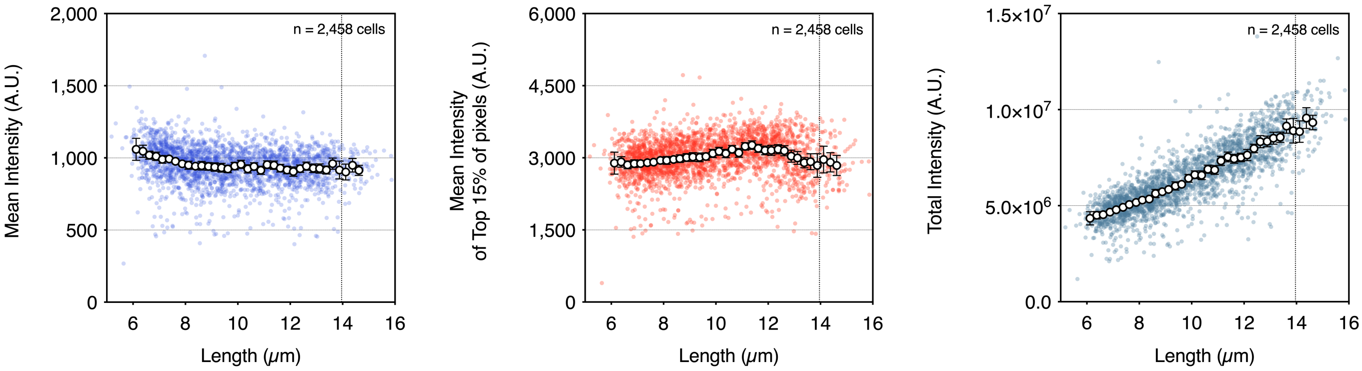


Imagestream: 10/10/18

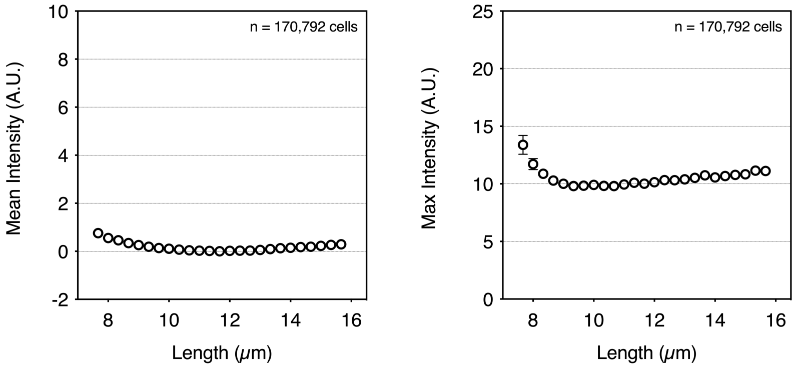

### pab2-mNG

PN5871
SPBC16E9.12c
poly(A) binding protein Pab2

Widefield: 19/04/18

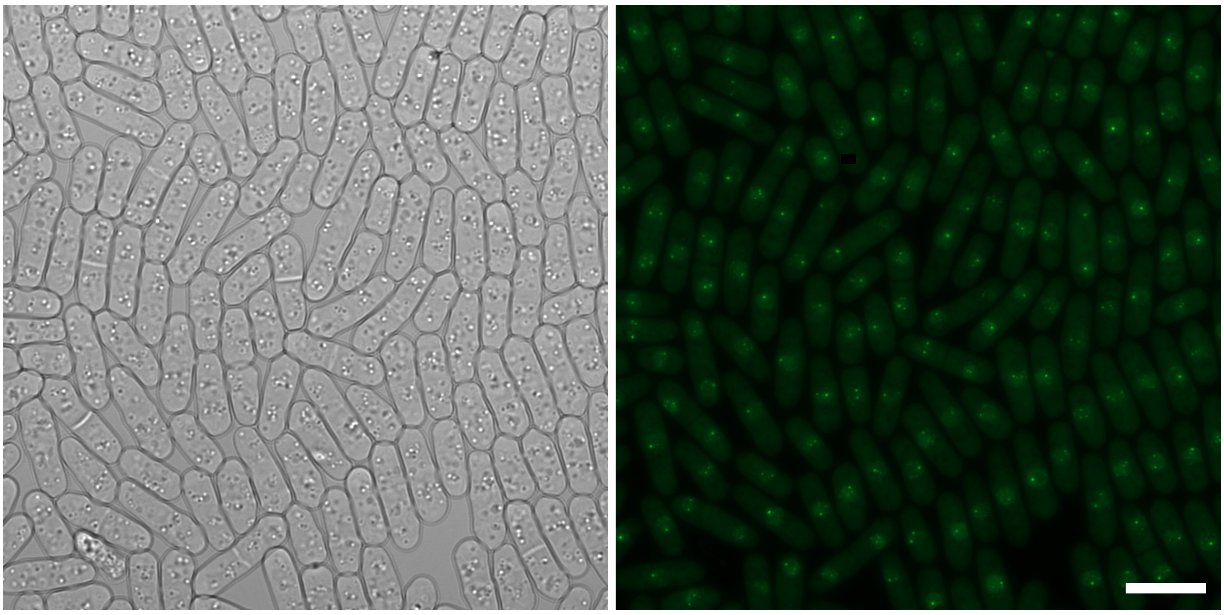

Widefield: 19/04/18

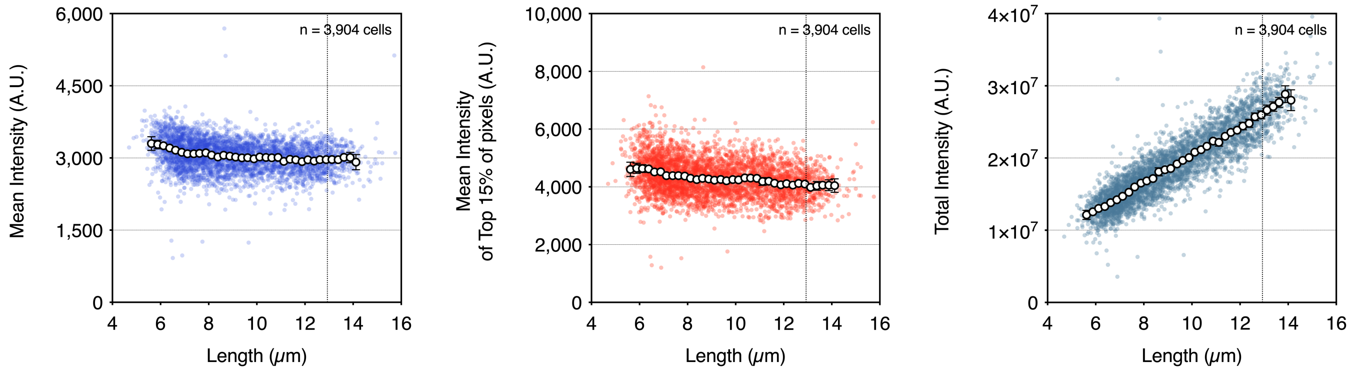


Imagestream: 17/10/18

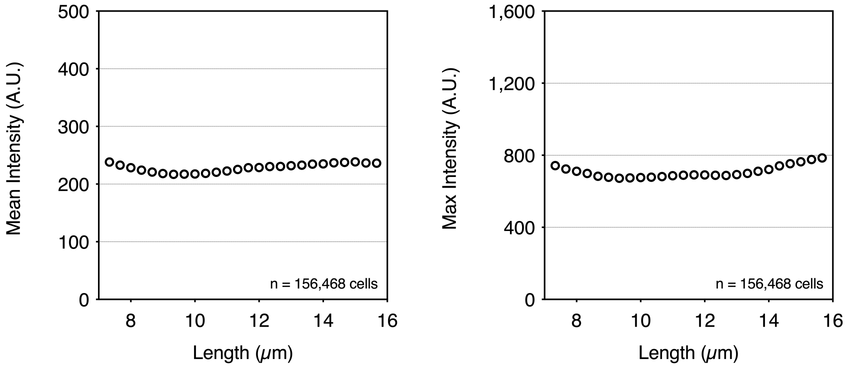

### pka1-GFP

PN5812
SPBC106.10
cAMP-dependent protein kinase catalytic subunit Pka 1

Widefield: 02/05/18

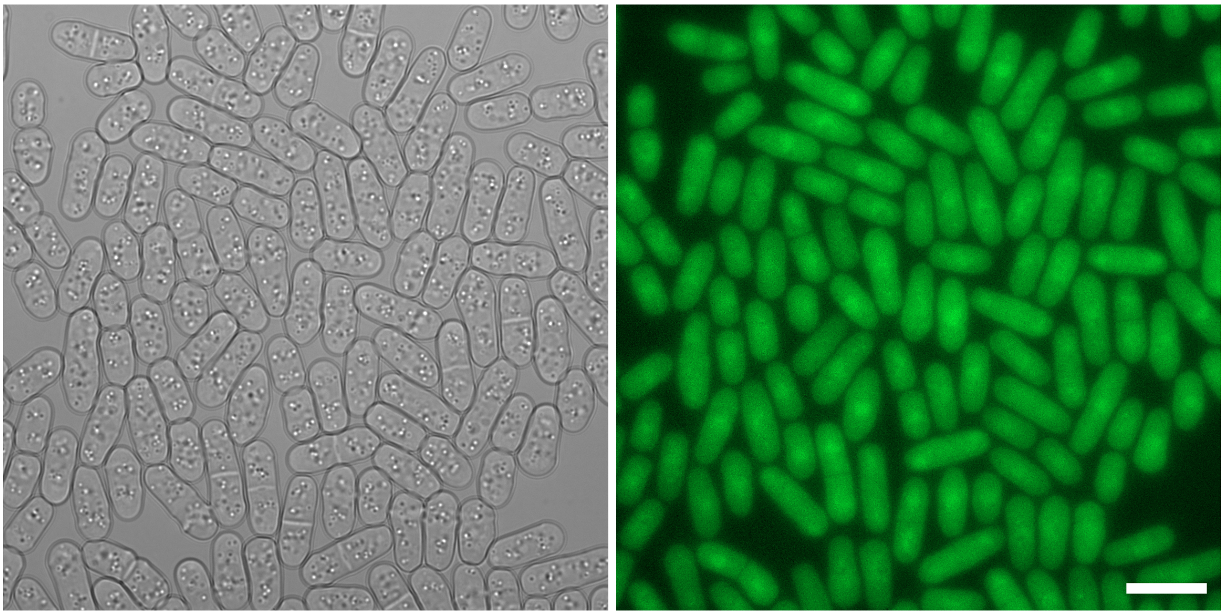

Widefield: 02/05/18

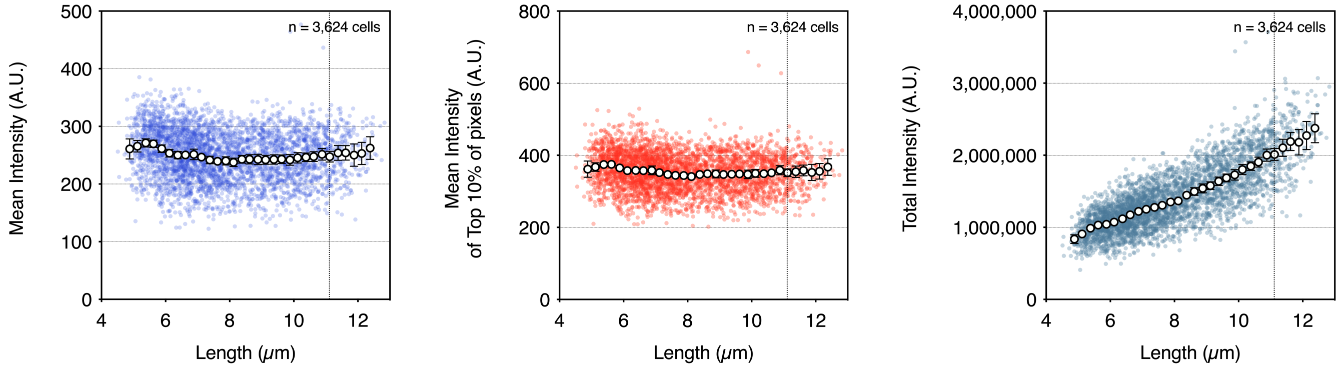


Imagestream: 10/10/18

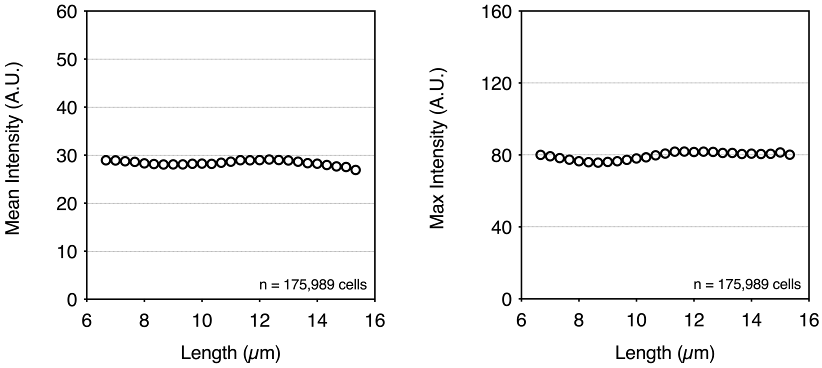

### pom1-mNG

PN5866
SPAC2F7.03c
DYRK family cell polairty protein kinase Pom1

Widefield: 19/04/18

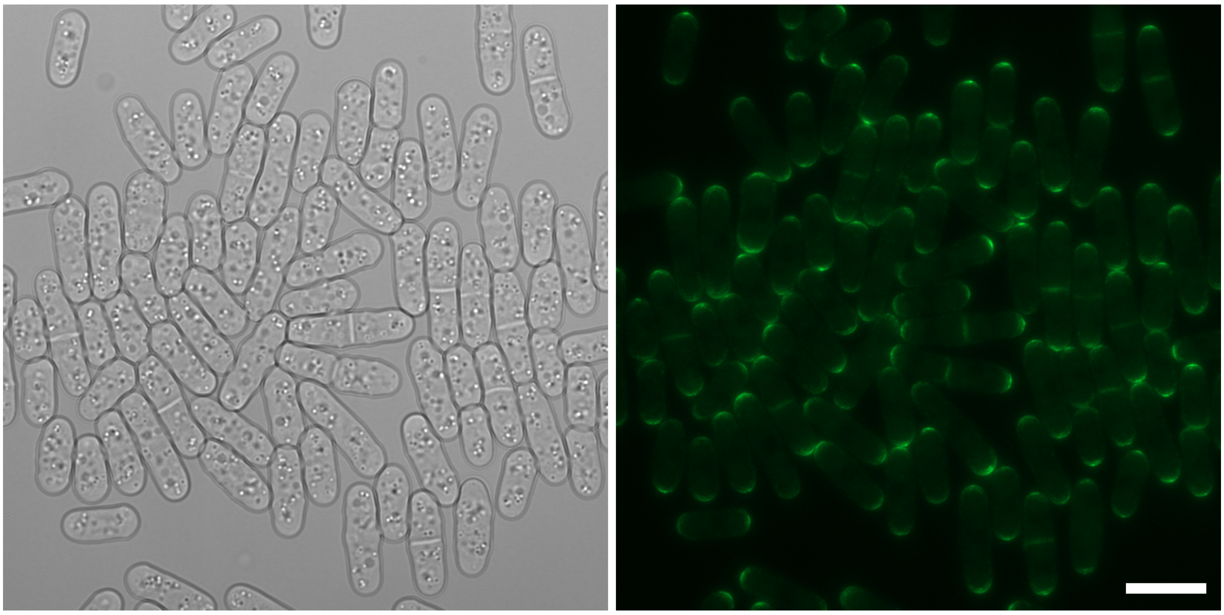

Widefield: 19/04/18

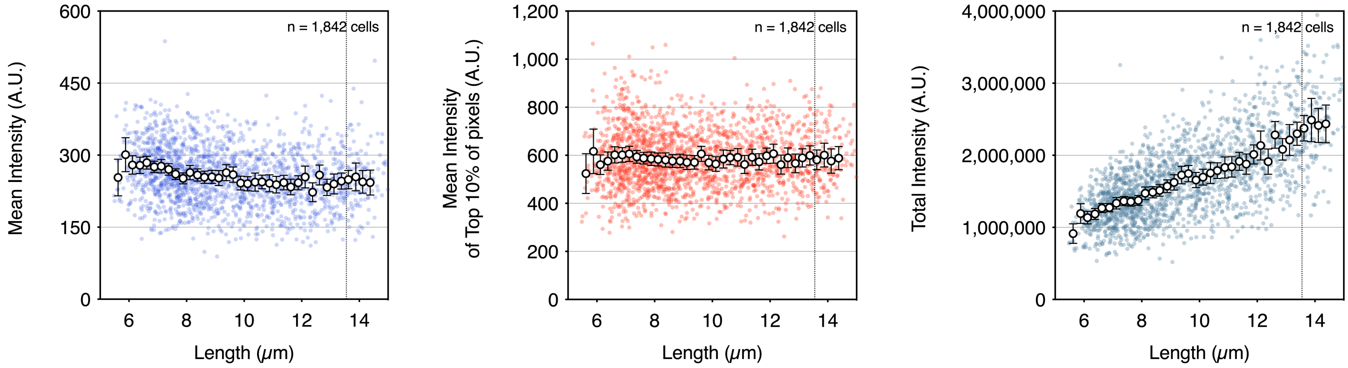


Imagestream: 17/10/18

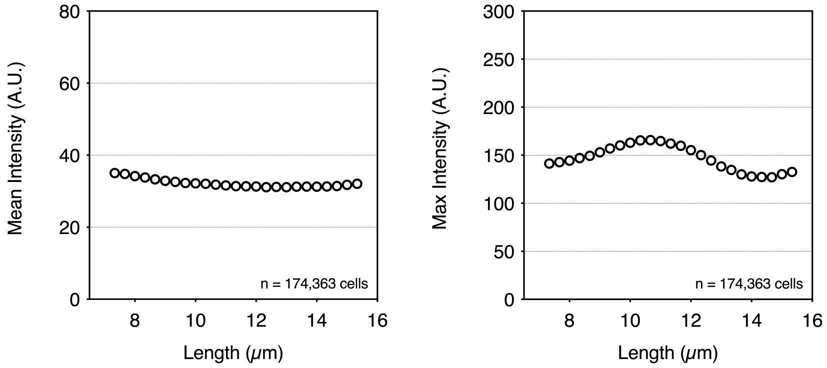

### ppa2-mNG

PN5862
SPBC16H5.07c
serine/threonine protein phosphatase Ppa2

Widefield: 19/04/18

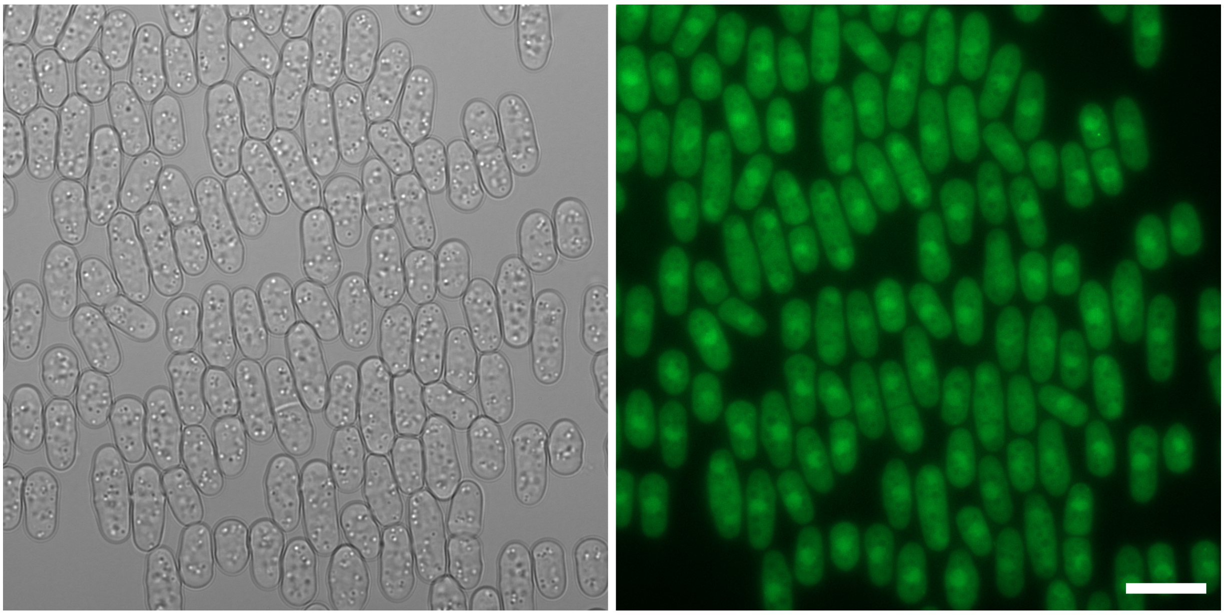

Widefield: 19/04/18

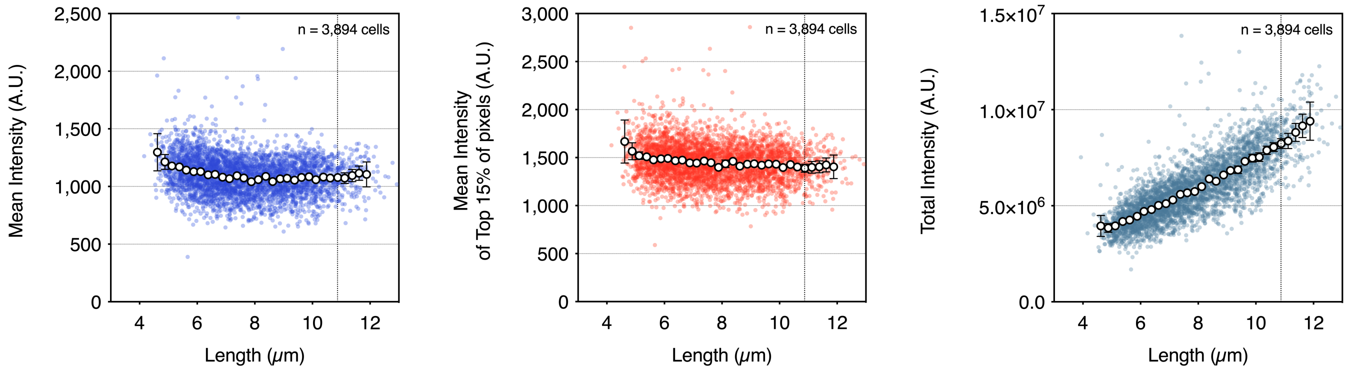


Imagestream: 17/10/18

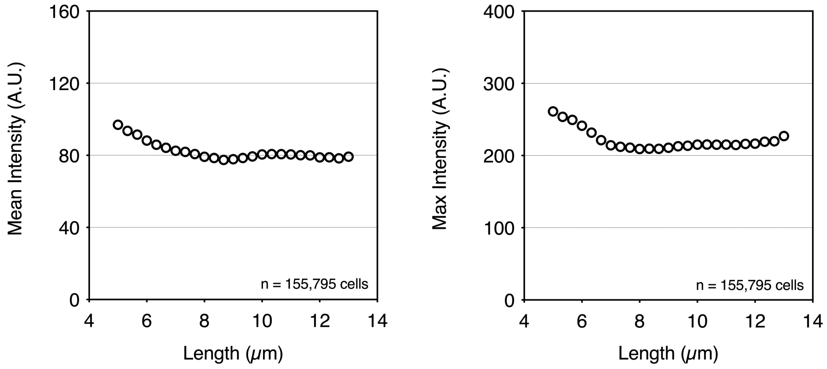

### pyp1-mNG

PN5863
SPAC26F1.10c
tyrosine phosphatase Pyp1

Widefield: 19/04/18

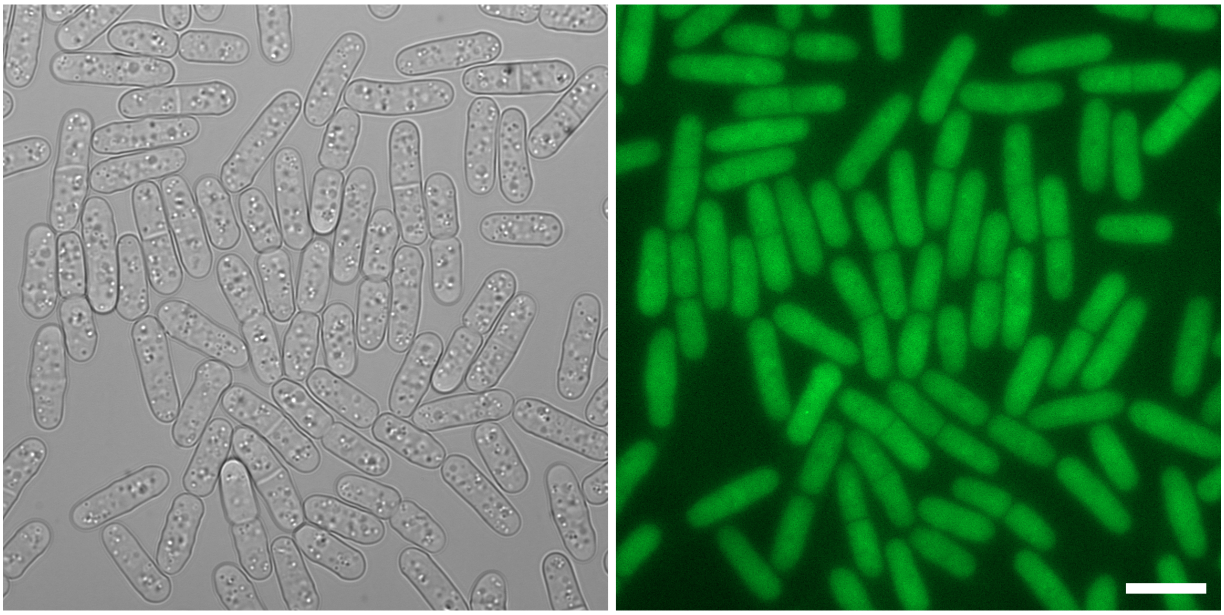

Widefield: 19/04/18

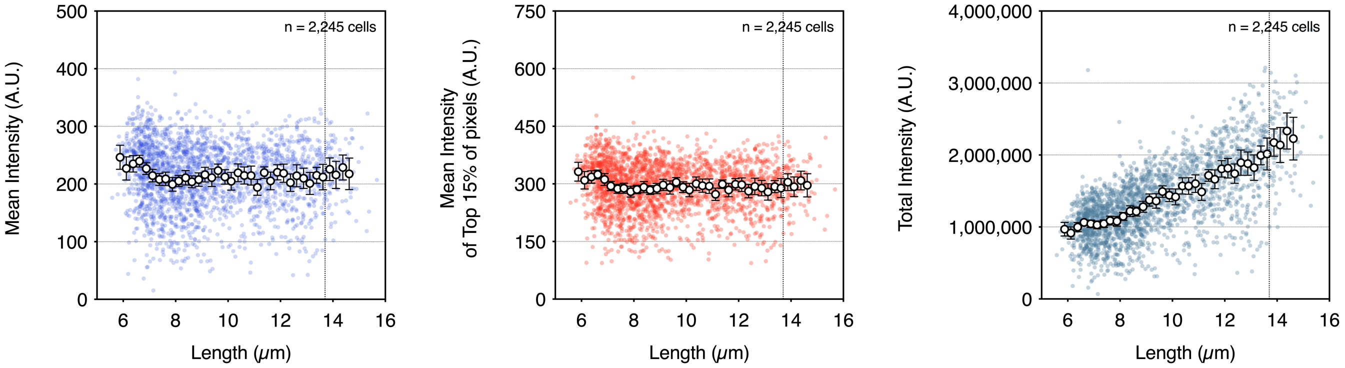


Imagestream: 11/10/18

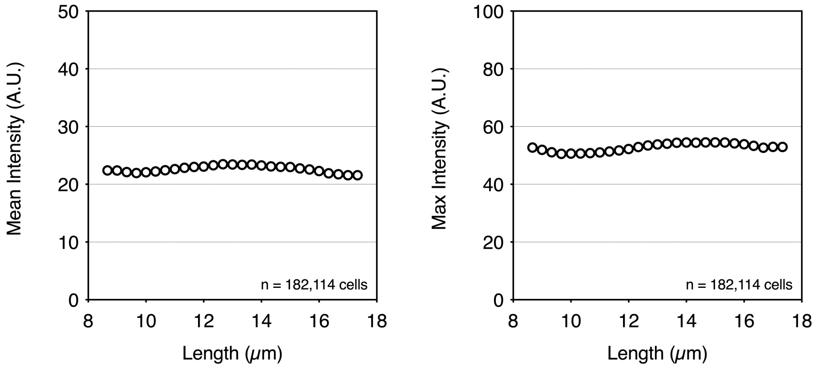

### sal3-mNG

PN5873
SPCC1840.03
karyopherin/importin beta family nuclear import signal

Widefield: 20/04/18

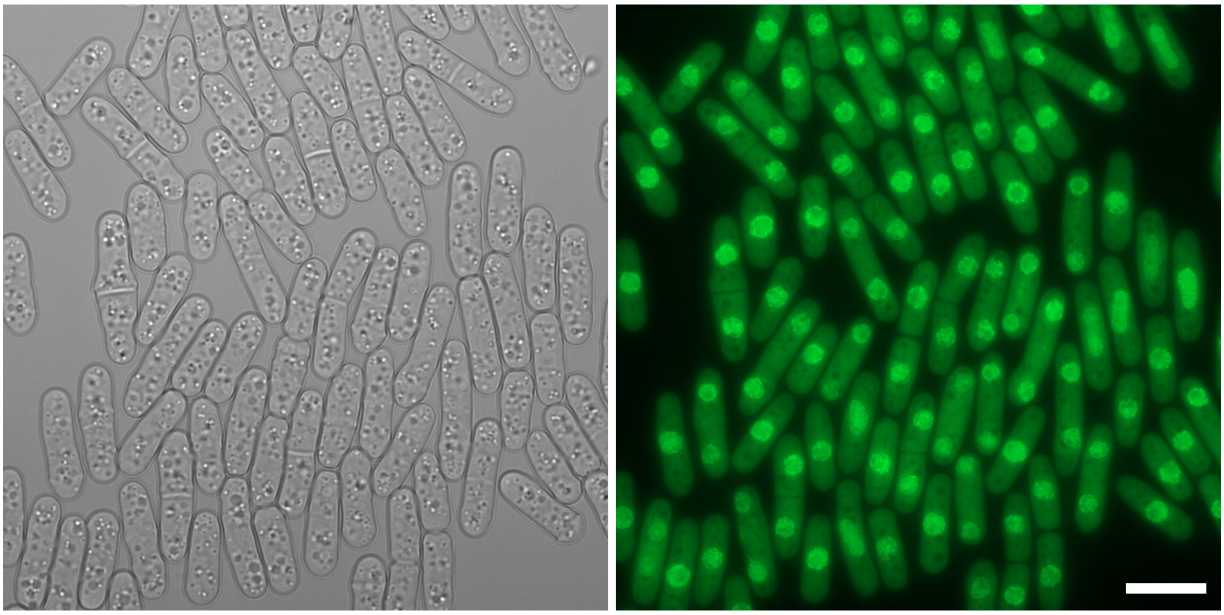

Widefield: 20/04/18

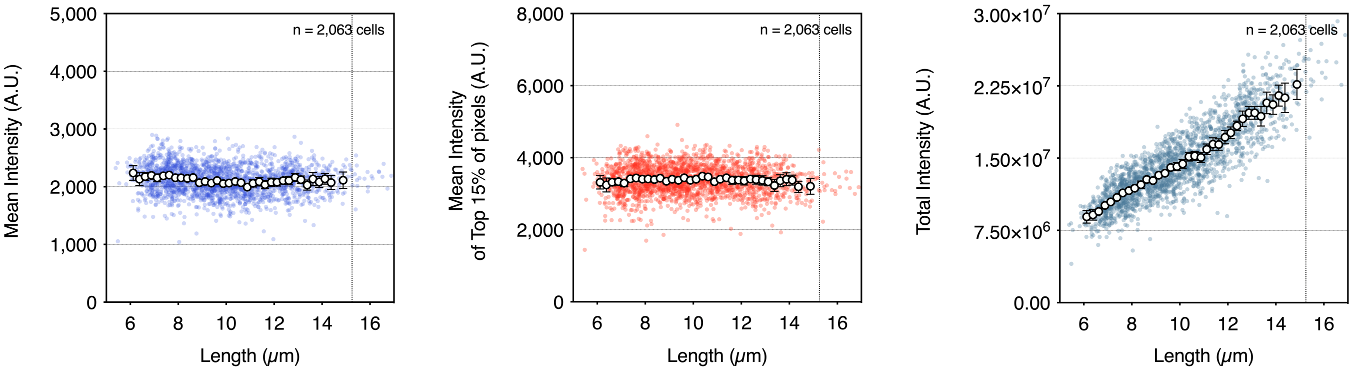


Imagestream: 10/10/18

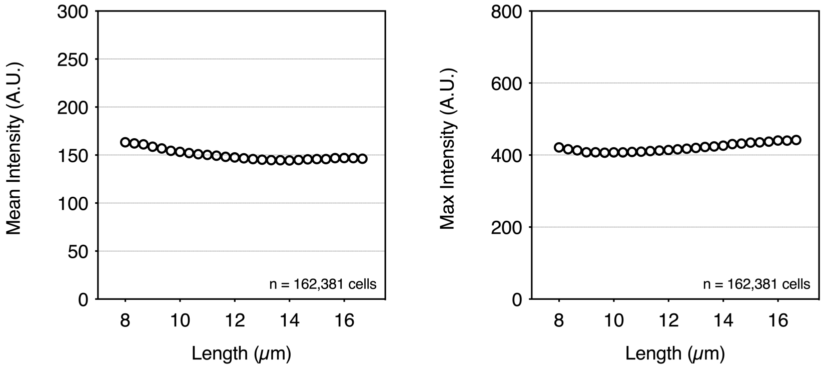

### sgf73-mNG

PN5850
SPCC126.04c
SAGA complex subunit

Widefield: 27/04/18

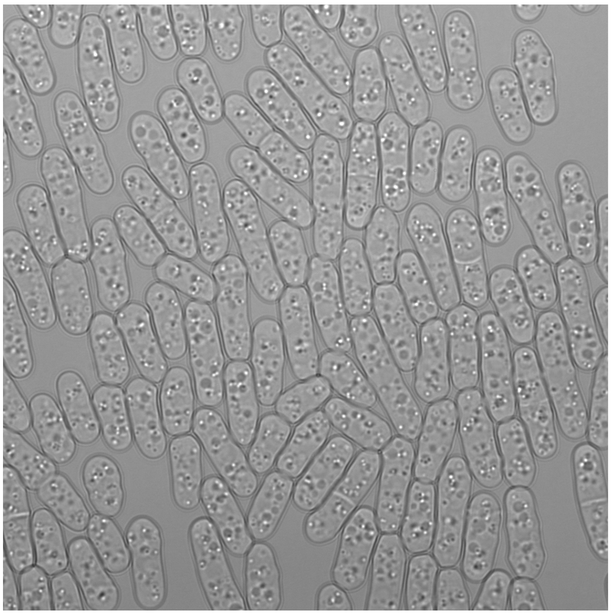

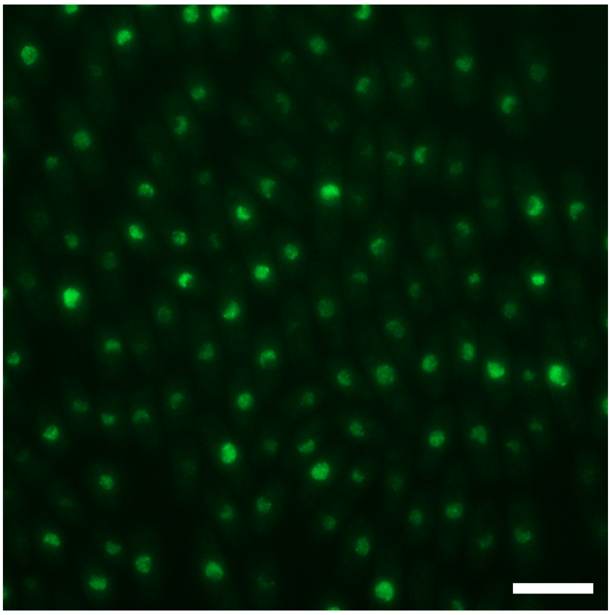

Widefield: 27/04/18

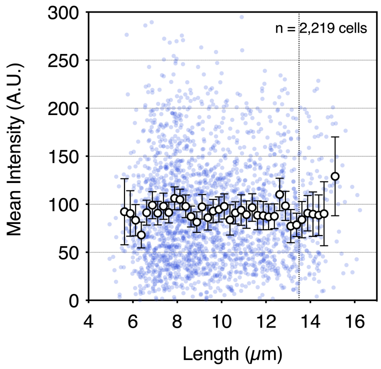


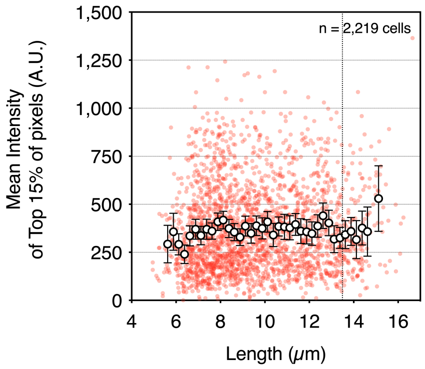


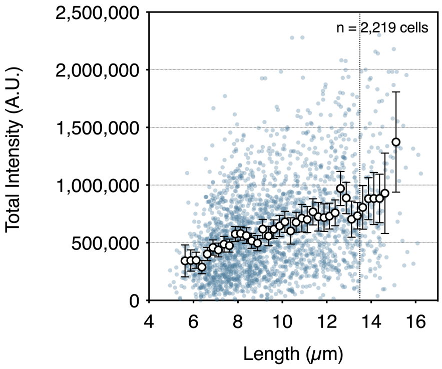


Imagestream: 17/10/18

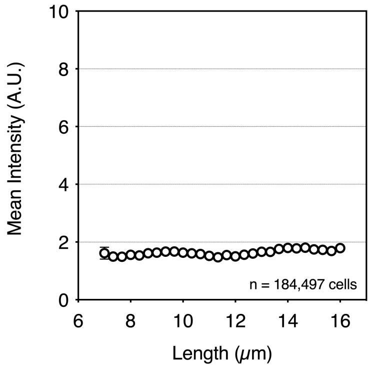


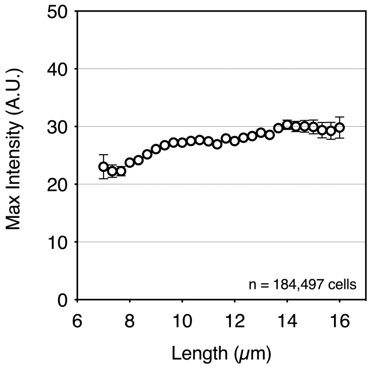

### ski3-mNG

PN5851
SPCC1919.05
Ski complex TPR repeat subunit Ski3 (predicted)

Widefield: 27/04/18

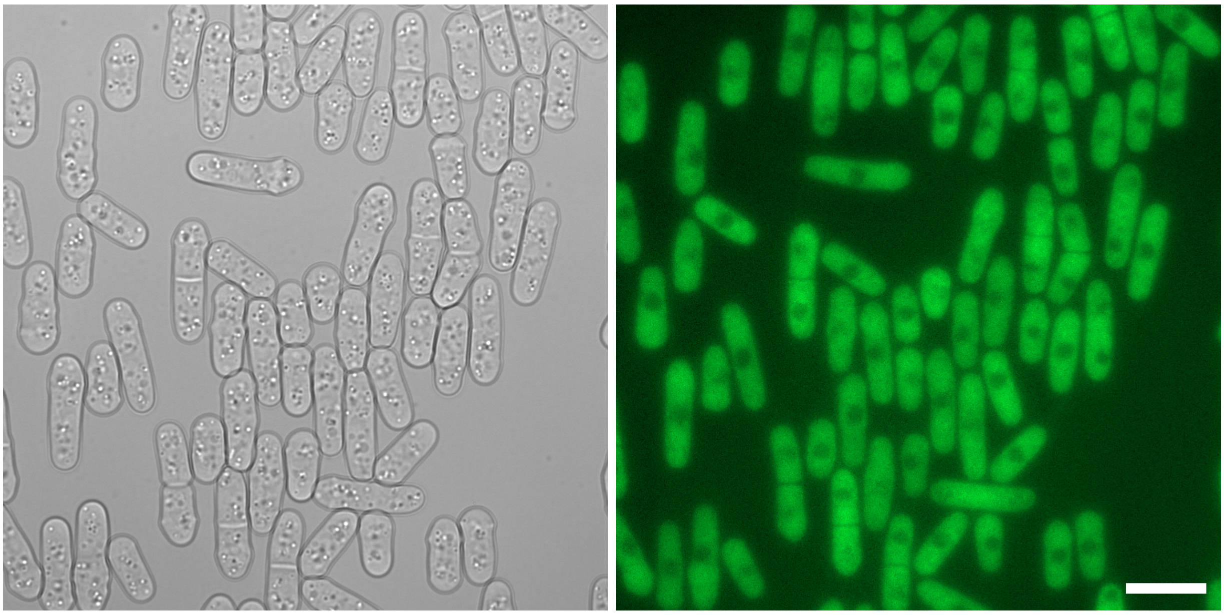

Widefield: 27/04/18

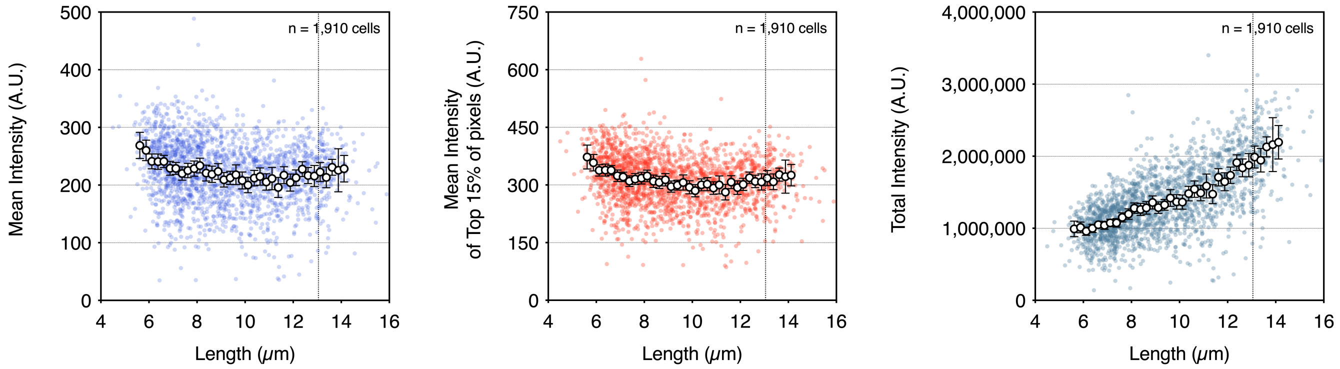


Imagestream: 17/10/18

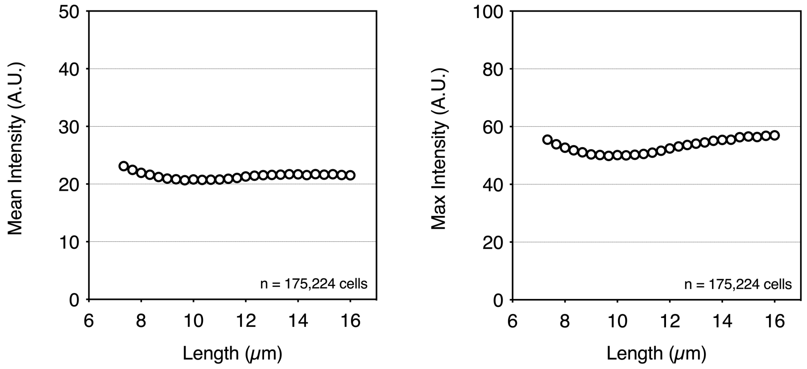

### snf5-mNG

PN5869
SPAC27F7.08c
SWI/SNF complex subunit Snf5

Widefield: 19/04/18

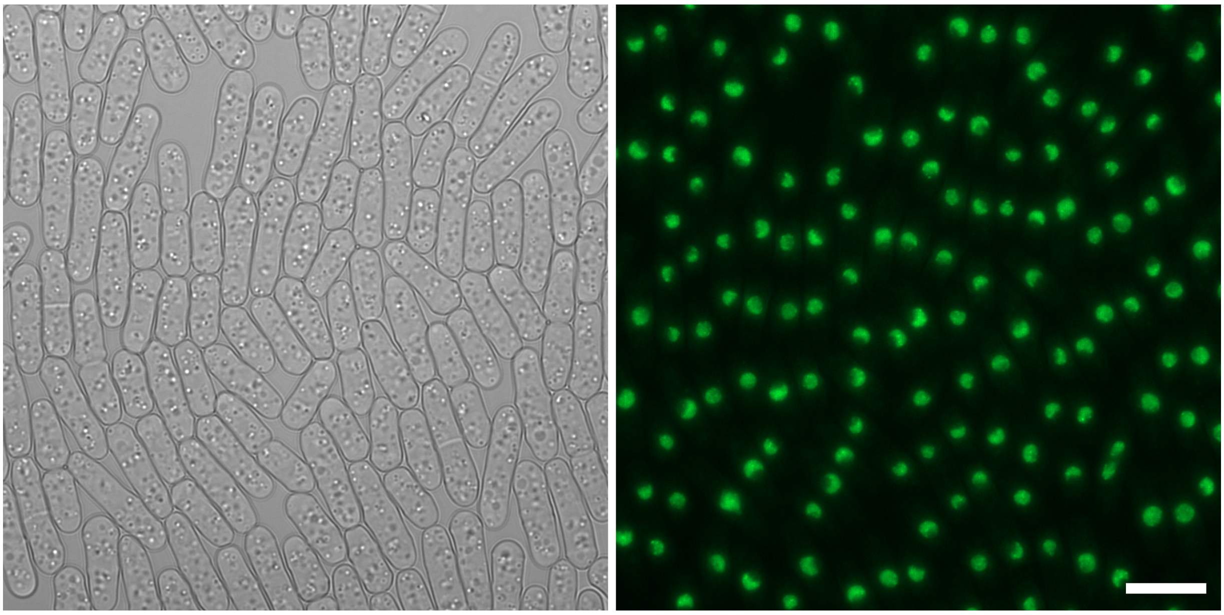

Widefield: 19/04/18

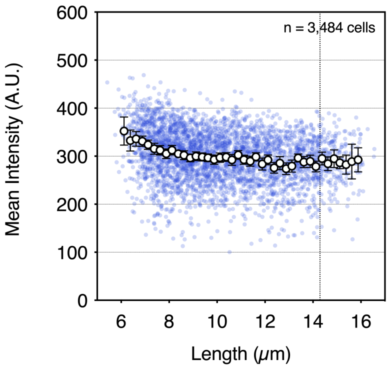


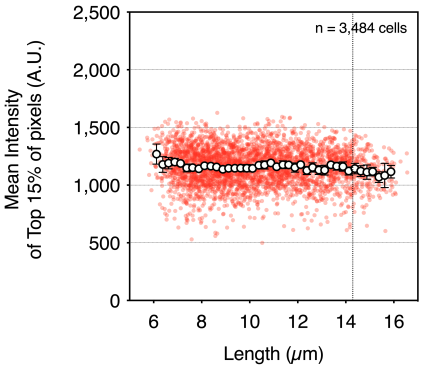


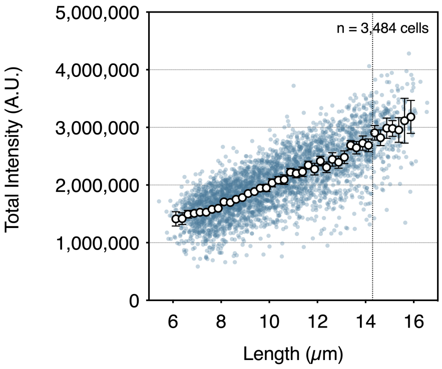


Imagestream: 12/10/18

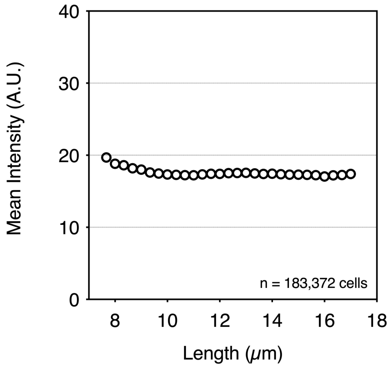


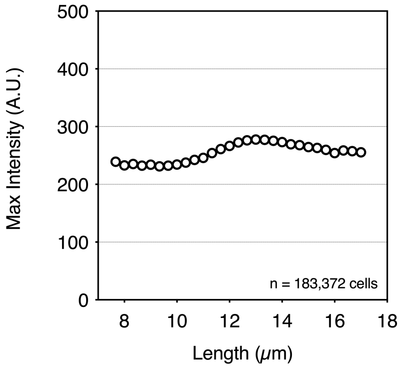

### sol1-mNG

PN5870
SPBC30B4.04c
SWI/SNF complex subunit Sol1

Widefield: 19/04/18

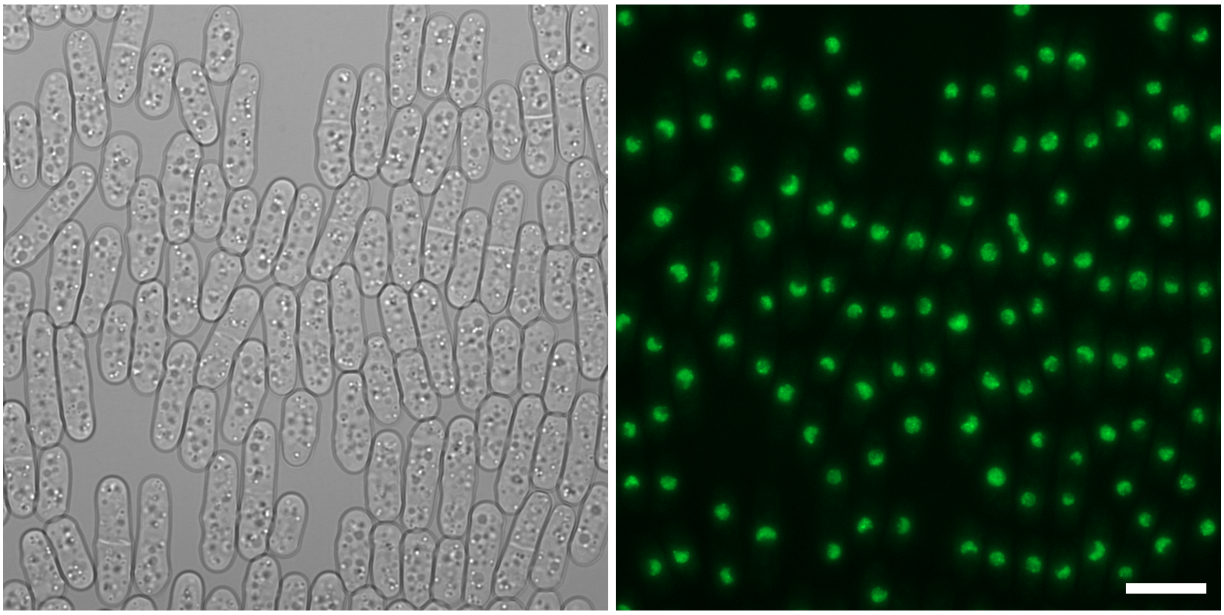

Widefield: 19/04/18

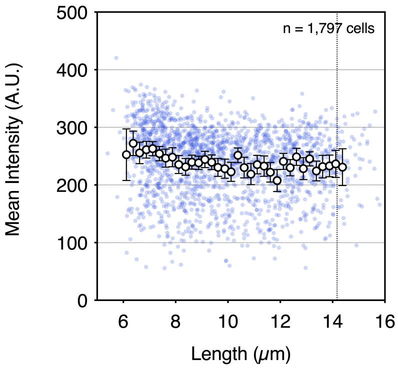


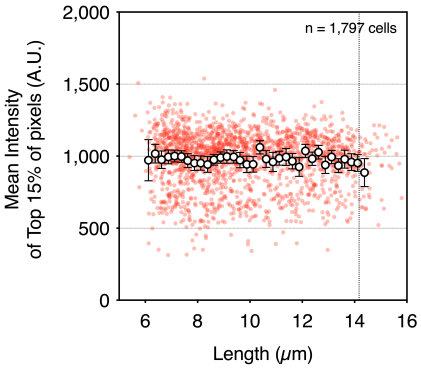


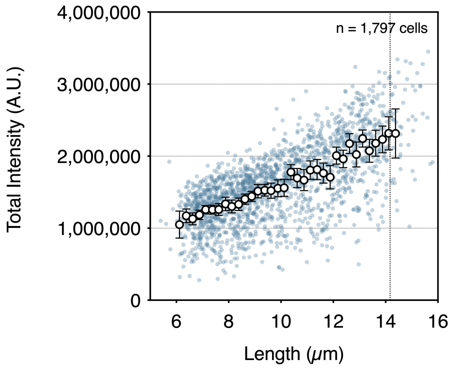


Imagestream: 13/10/18

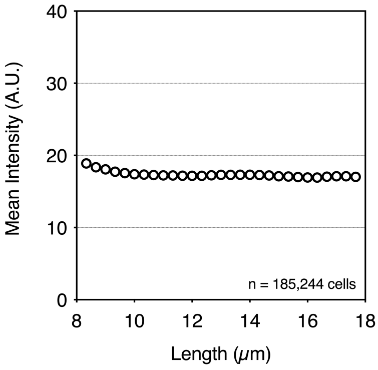


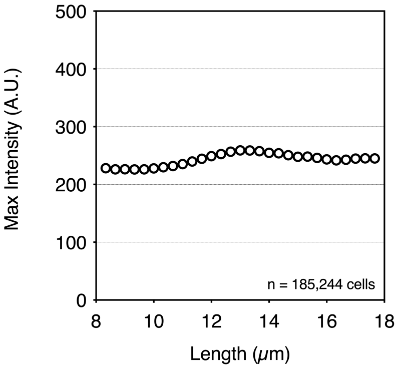

### SPAC27E2.03c-mNG

PN5865
Obg-like ATPase, human
OLA1 ortholog (predicted)

Widefield: 19/04/18

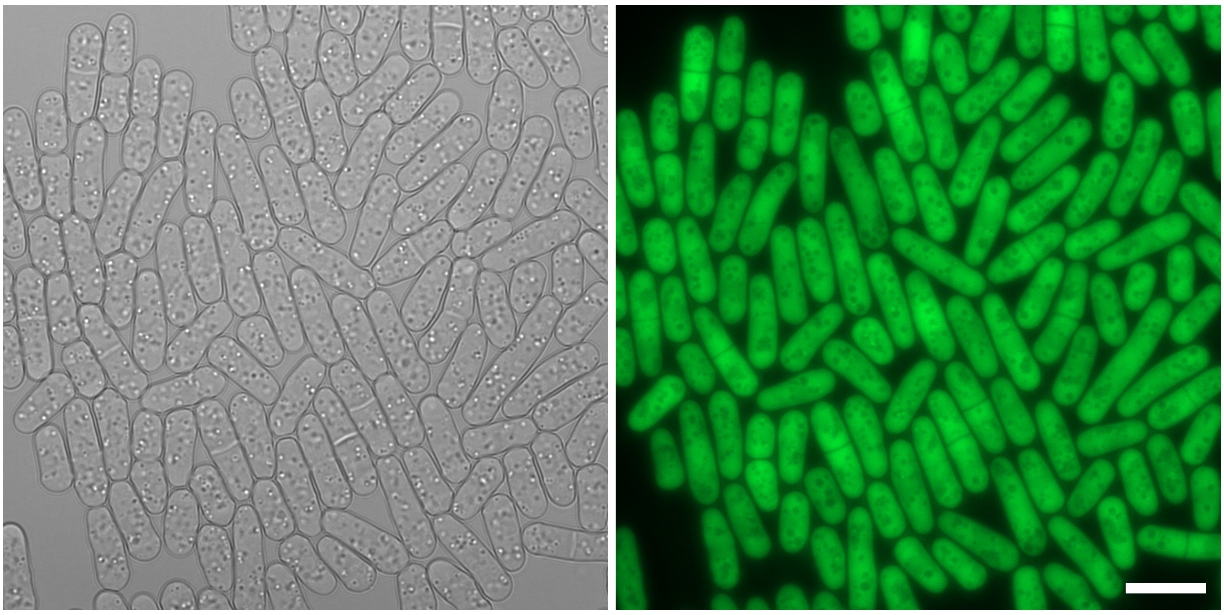

Widefield: 19/04/18

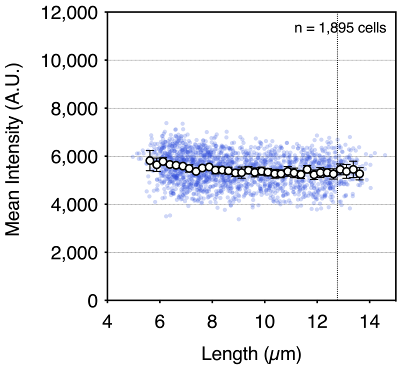


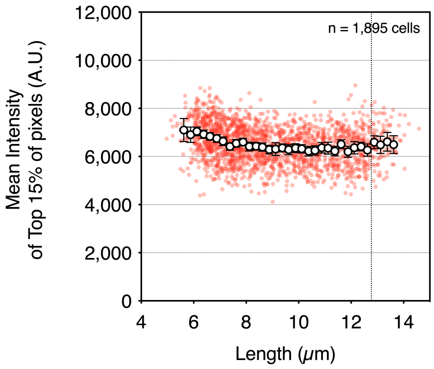


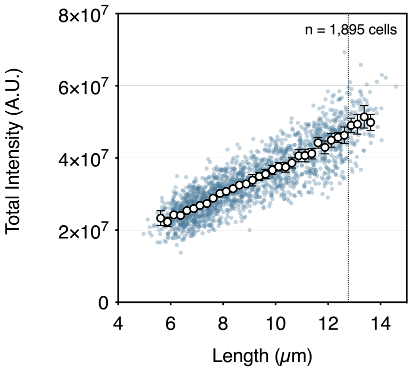


Imagestream: 17/10/18

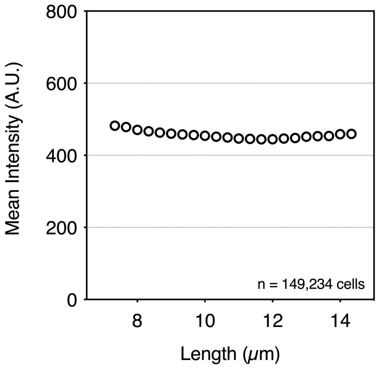


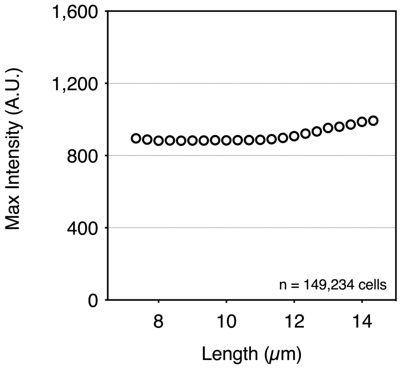

### suc1-mNG

PN5820/PN5895
SPBC1734.14c
cyclin-dependent protein kinase regulatory subunit Suc1

Widefield: 19/10/18

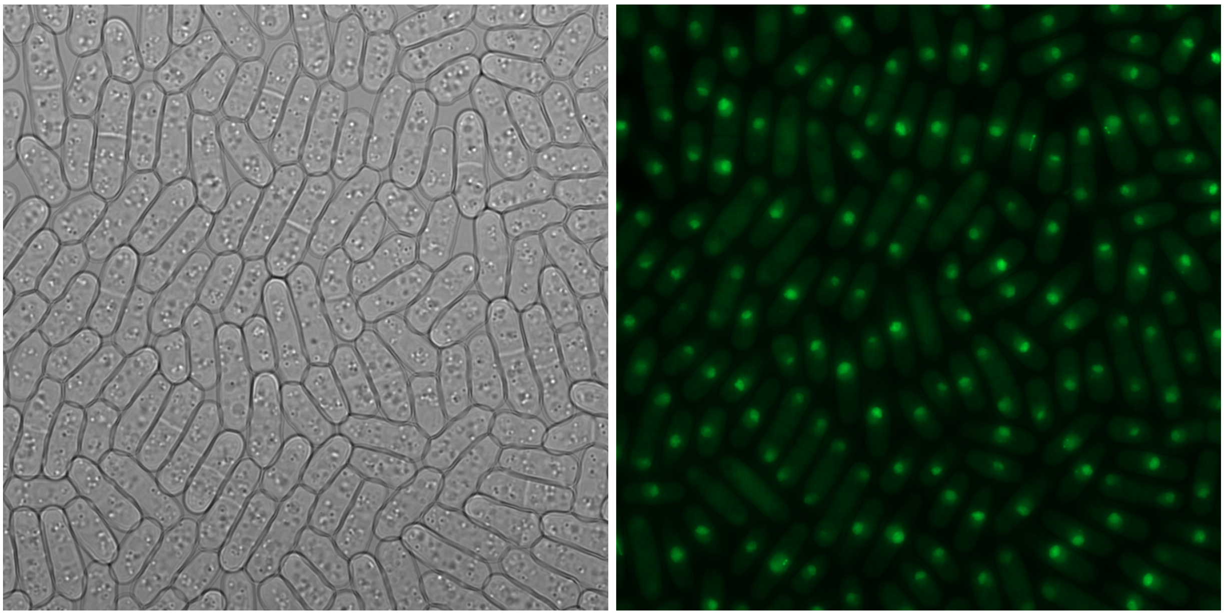

Widefield: 19/10/18

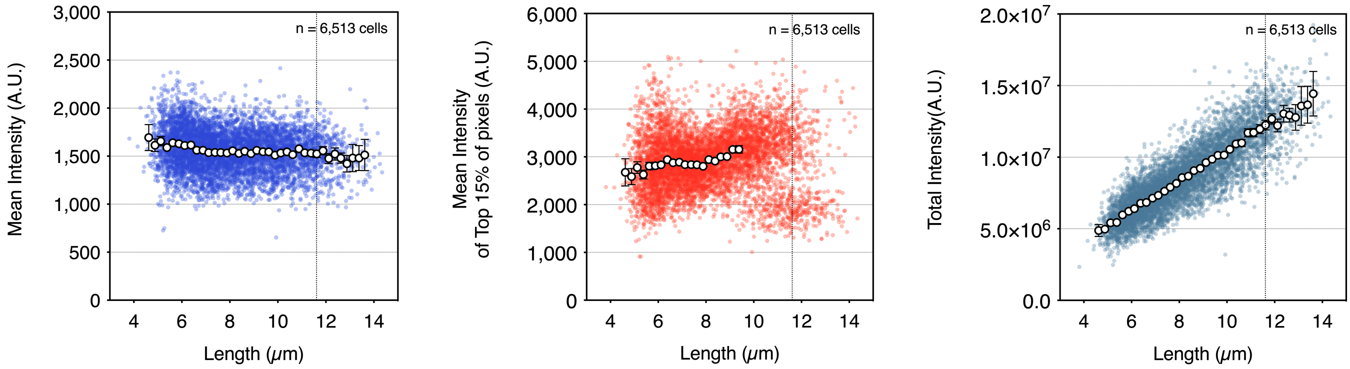


Widefield: 12/03/21

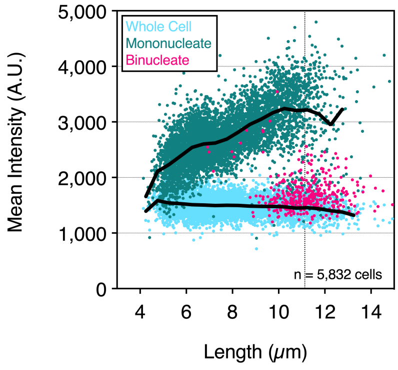


Imagestream: 10/10/18

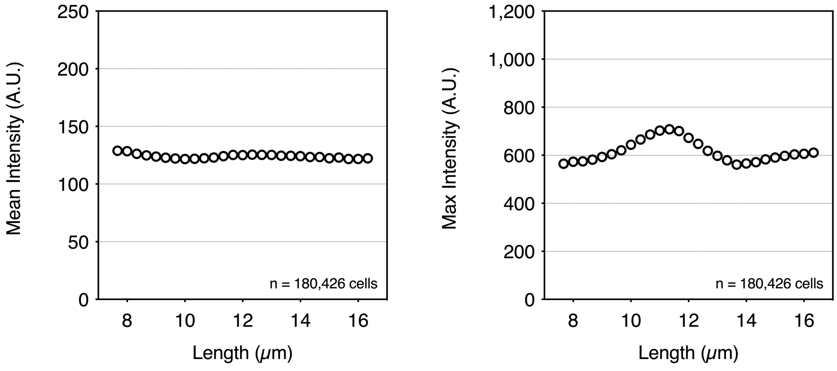

### wee1-GFP

PN5841/PN5890
SPCC18B5.03
M phase inhibitor protein kinase Wee1

Widefield: 27/04/18

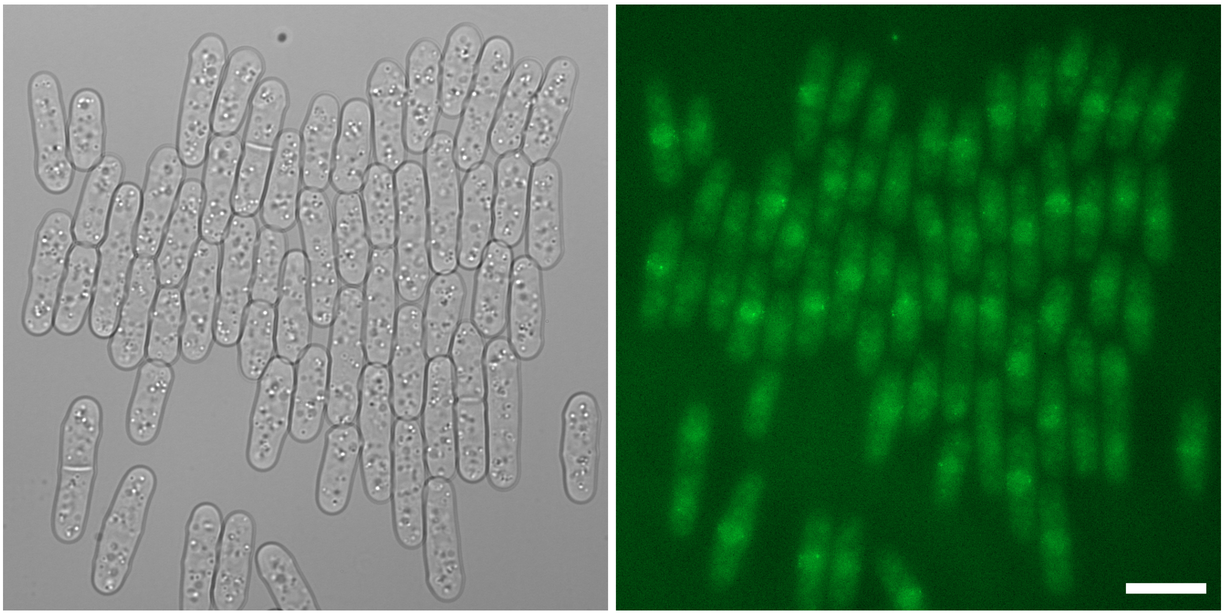

Widefield: 27/04/18

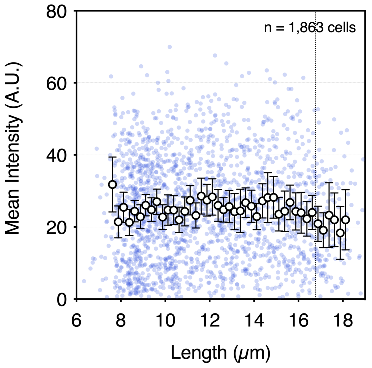


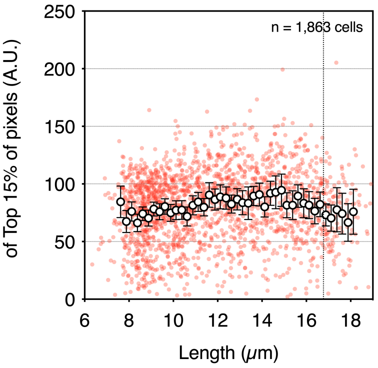


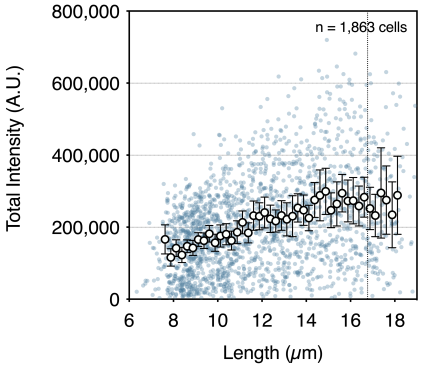


Widefield: 12/03/21

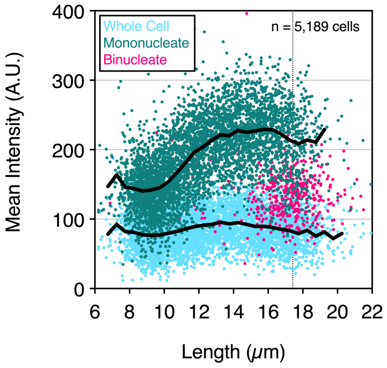


Imagestream: 10/10/18

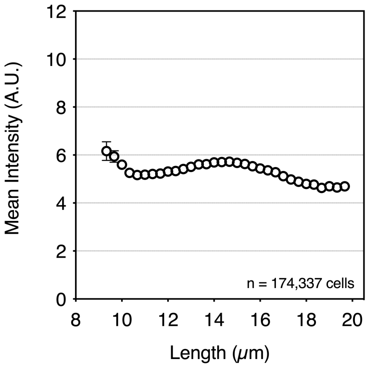


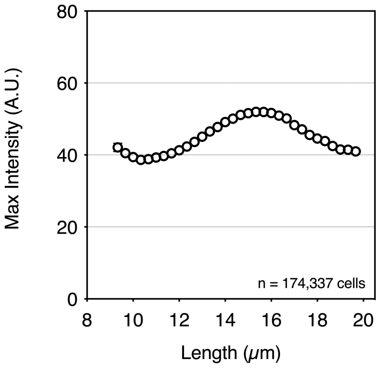

### zfs1-mNG

PN5867
SPBC1718.07c
CCH tandem zinc finger protein, human Tristetrapolin homolog Zfs1, involved in mRNA metabolism

Widefield: 27/04/18

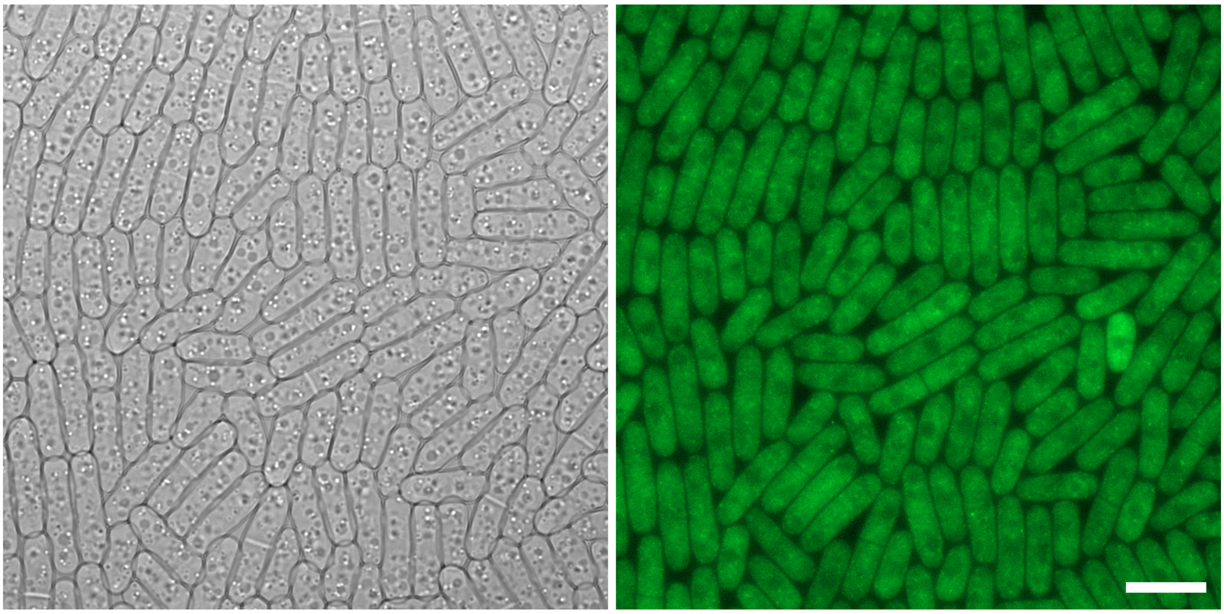

Widefield: 27/04/18

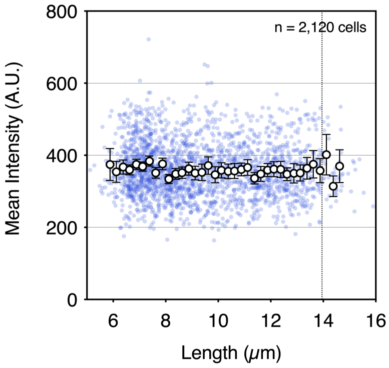


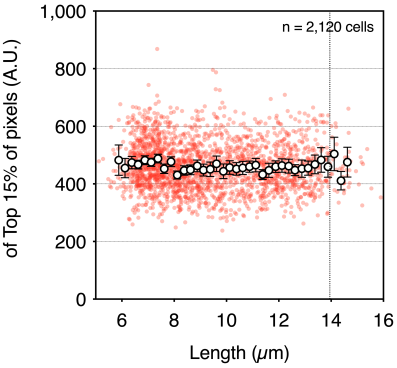


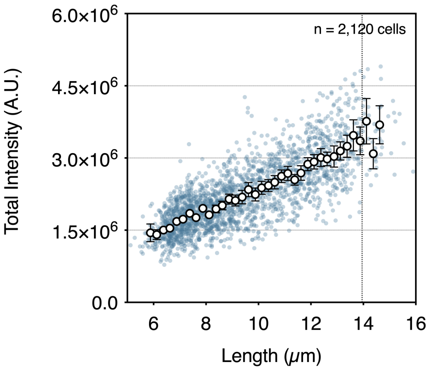


Imagestream: 10/10/18

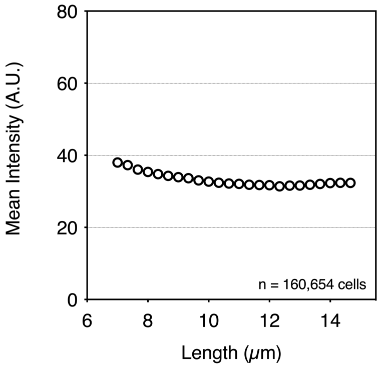


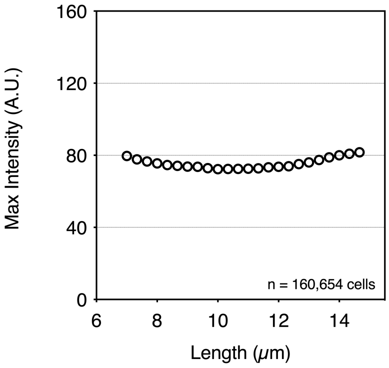

### cdc10-mNG

PN5813
SPBC336.12c
MBF transcription factor complex subunit Cdc10

Widefield: 27/04/18

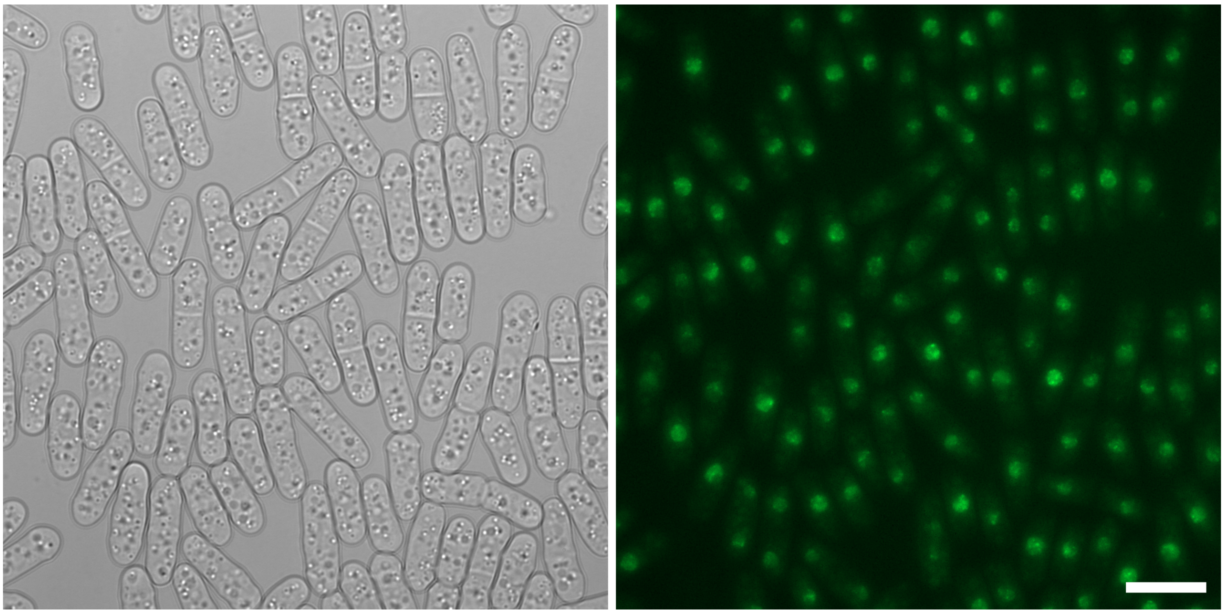

Widefield: 27/04/18

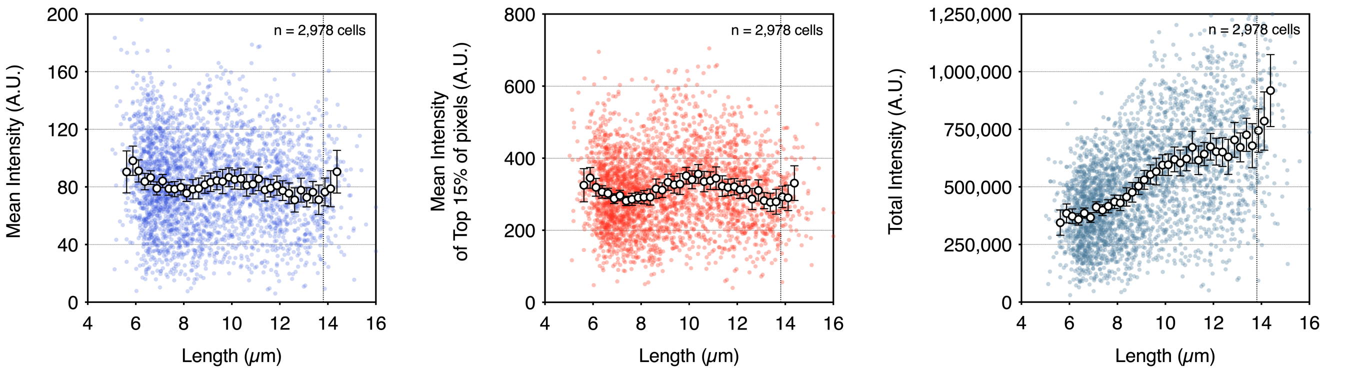


Imagestream: 10/10/18

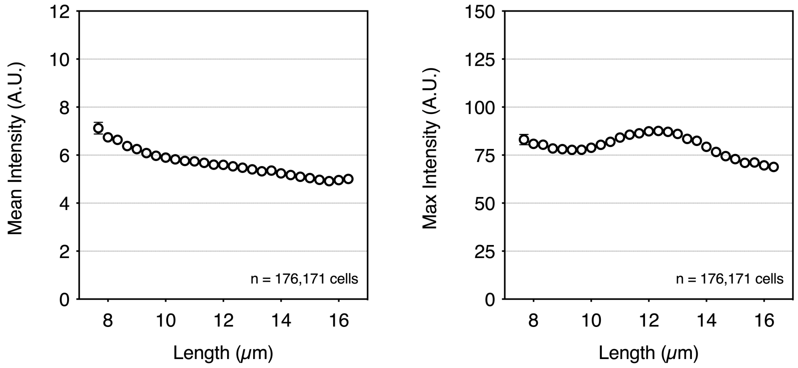

### cdc18-mNG

PN5860/PN5896
SPBC14C8.07c
MCM loader

Widefield: 19/10/18

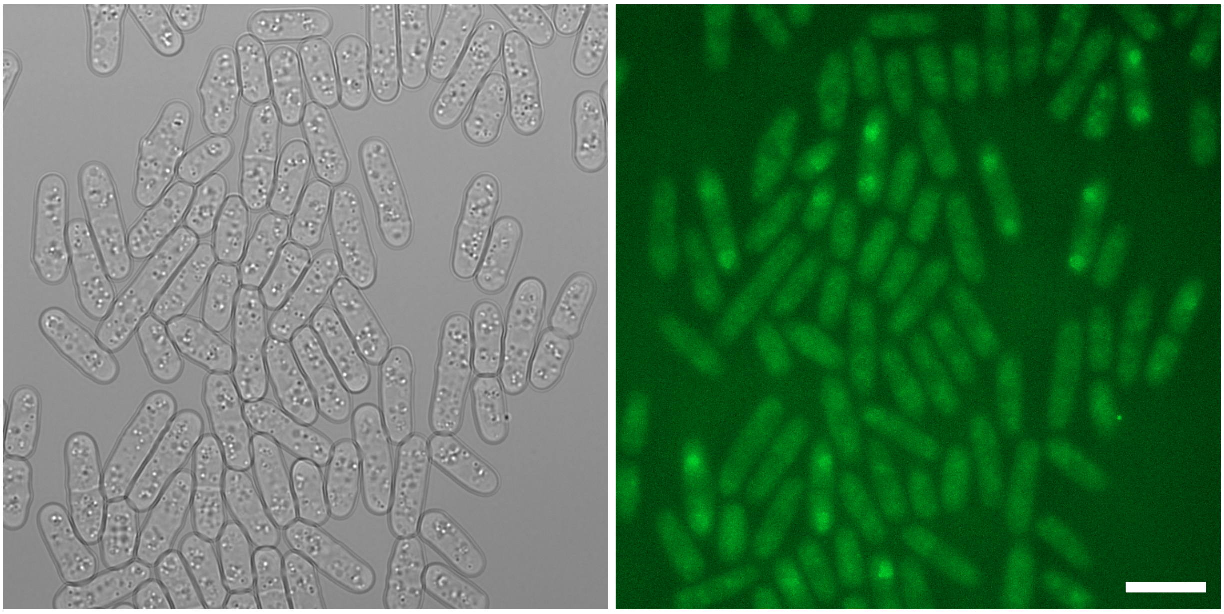

Widefield: 19/10/18

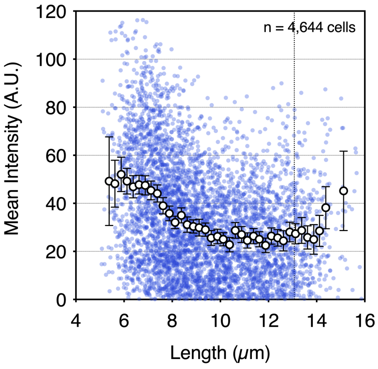


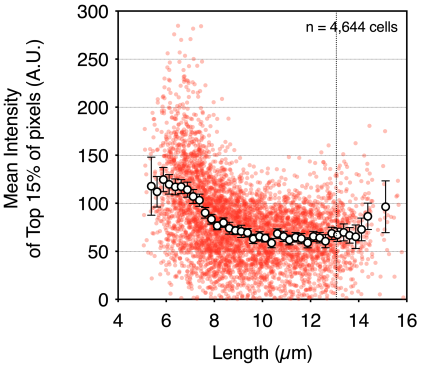


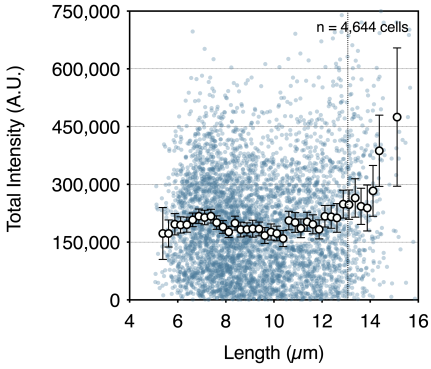


Widefield: 09/08/18

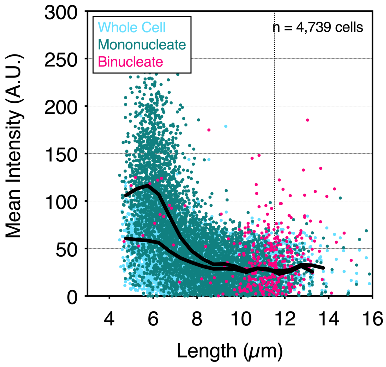


Imagestream: 10/10/18

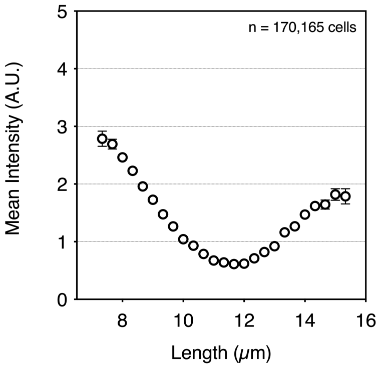


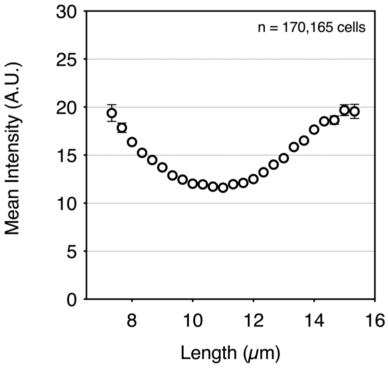

### cdc20-mNG

PN5826
SPBC25H2.13c
DN polymerase epsilon catalytic subunit Pol2

Widefield: 27/04/18

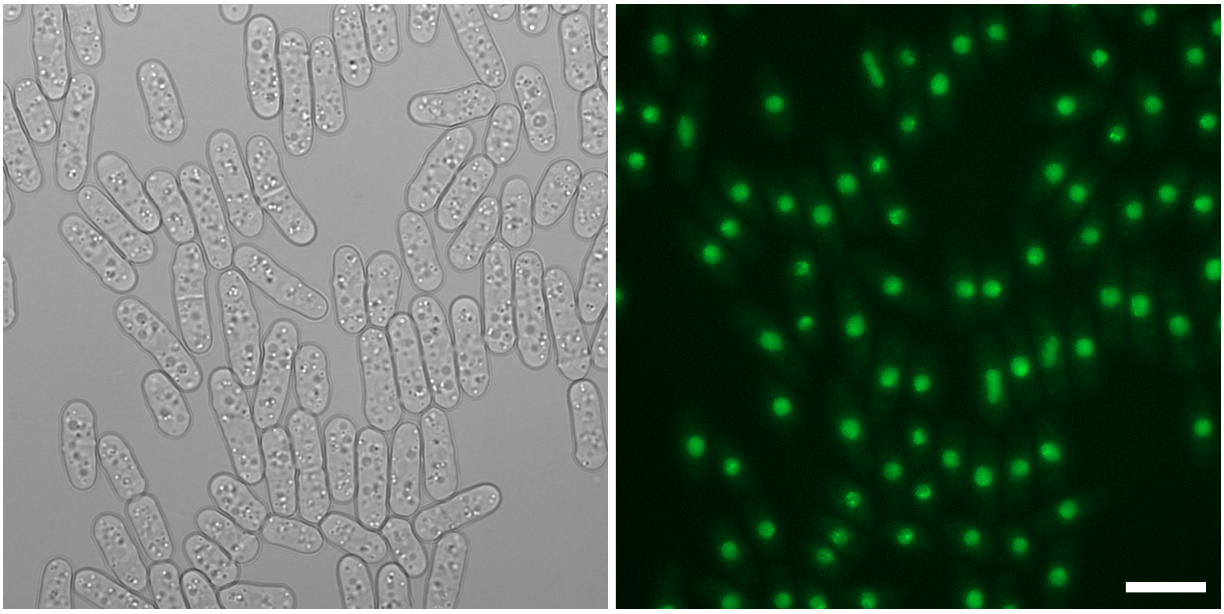

Widefield: 27/04/18

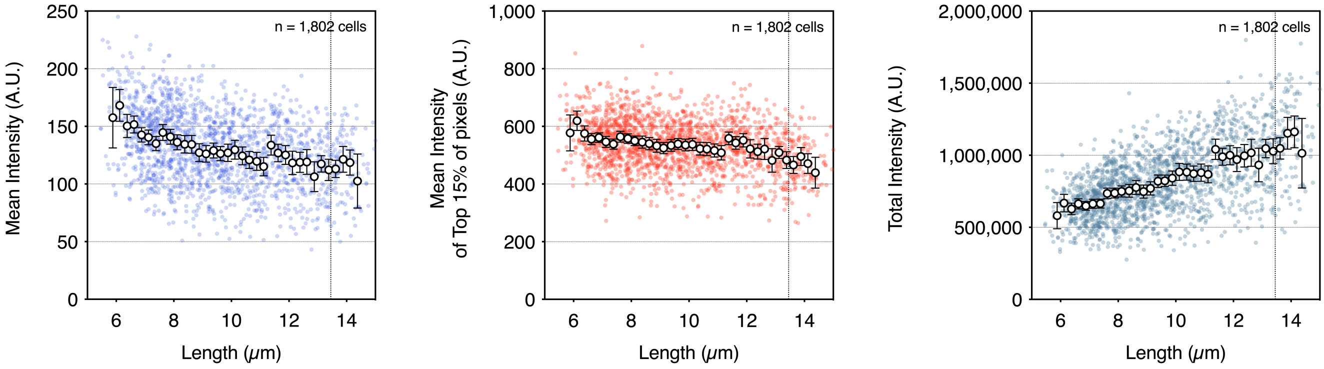


Imagestream: 17/10/18

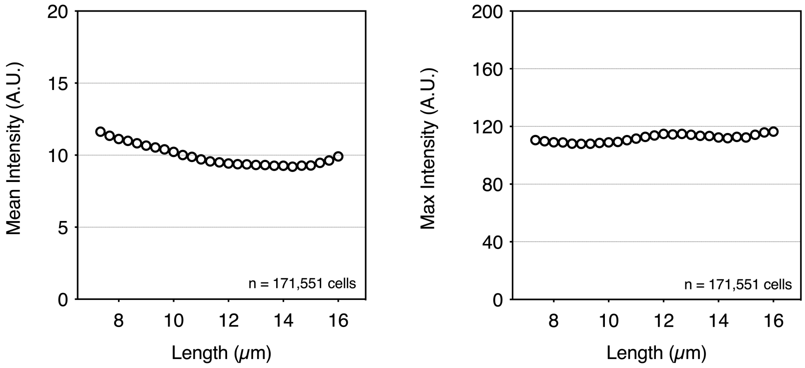

### cig1-mNG

PN5814/PN5885
SPCC4E9.02
cyclin Cig1

Widefield: 19/10/18

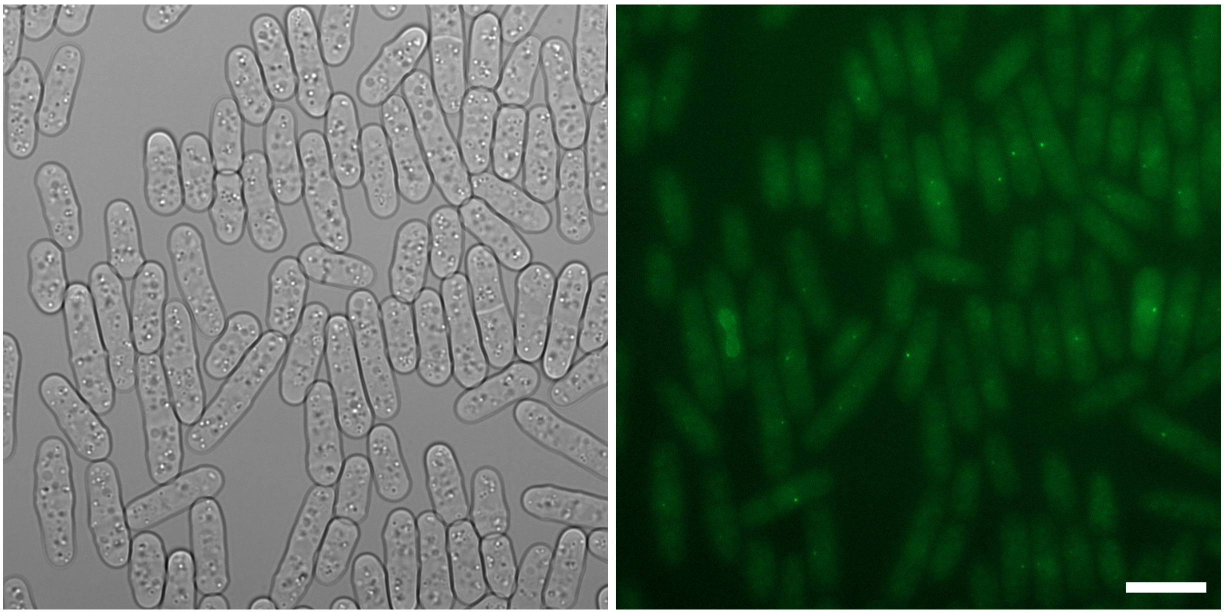

Widefield: 19/10/18

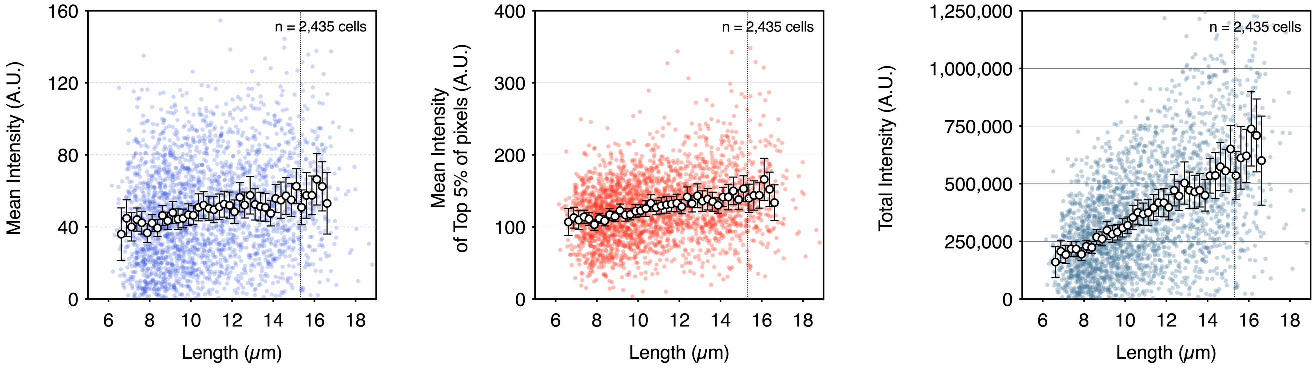


Widefield: 09/08/18

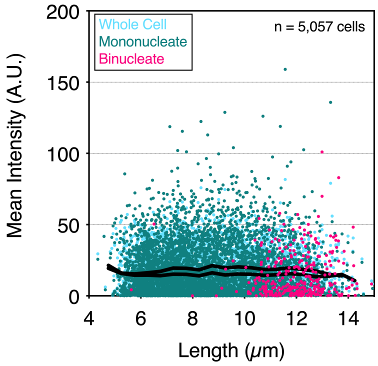


Imagestream: 10/10/18

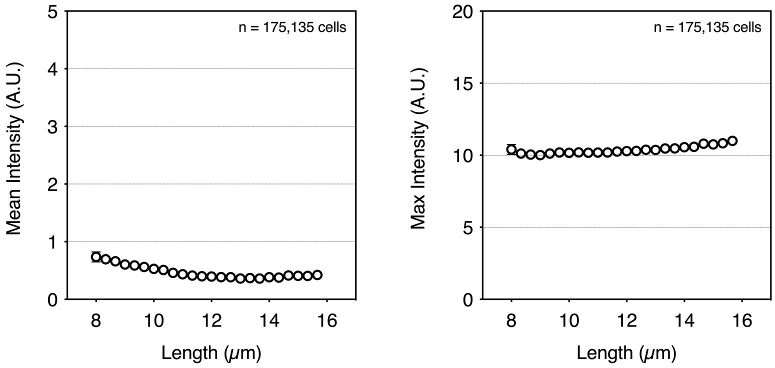

### cig2-mNG

PN5818/PN5899
SPAPB2B4.03
G1/S-specific B-type cyclin Cig2

Widefield: 19/04/18

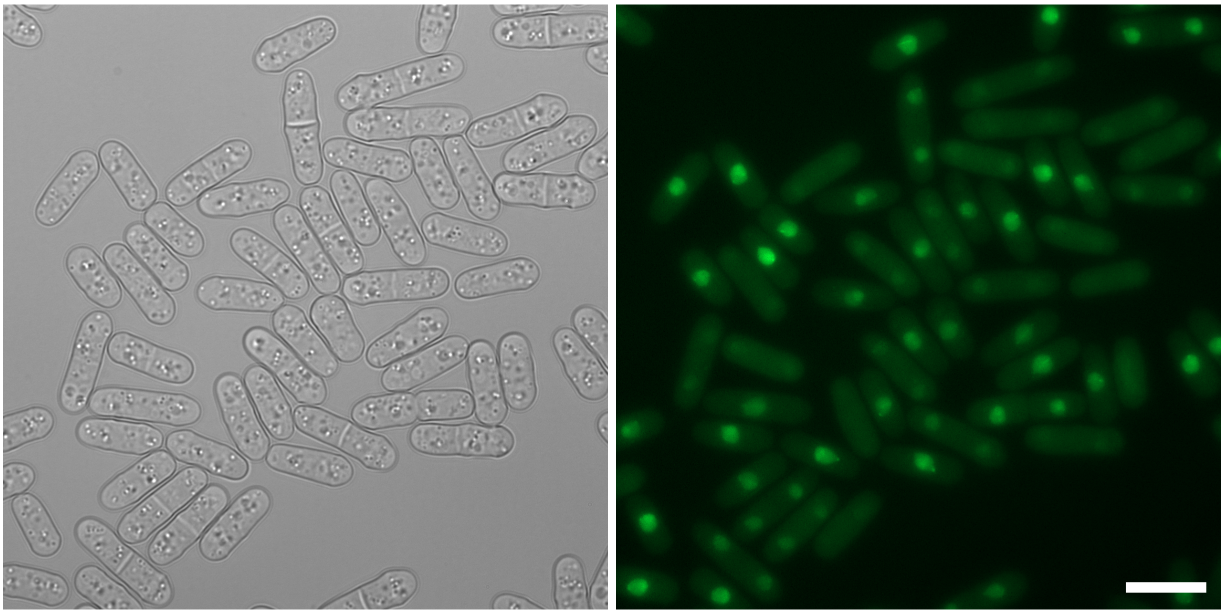

Widefield: 19/04/18

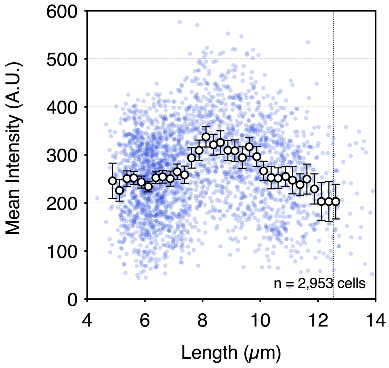


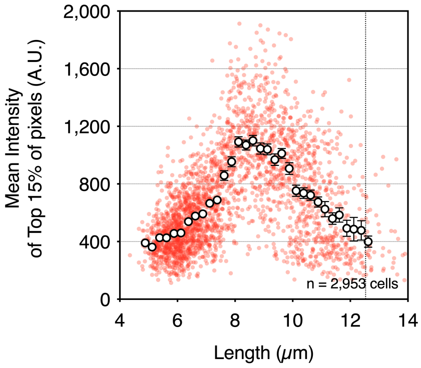


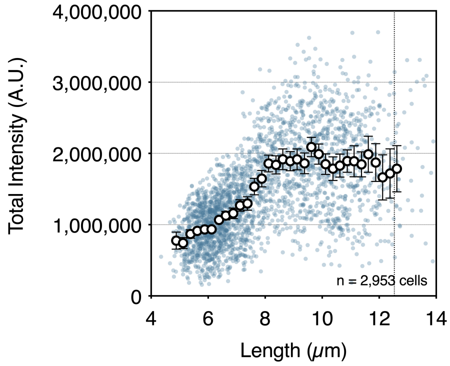


Widefield: 12/03/21

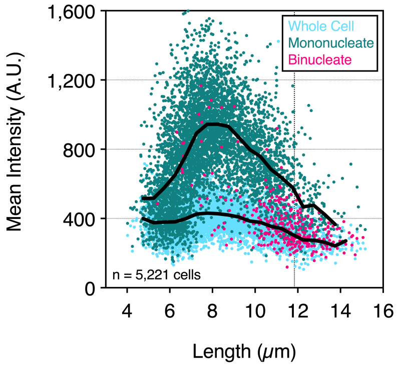


Imagestream: 10/10/18

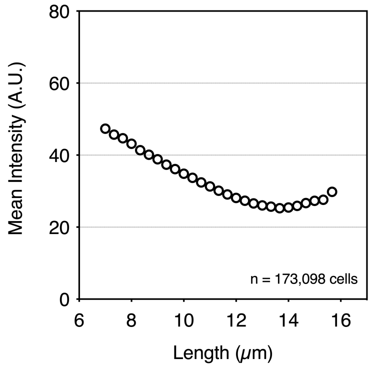


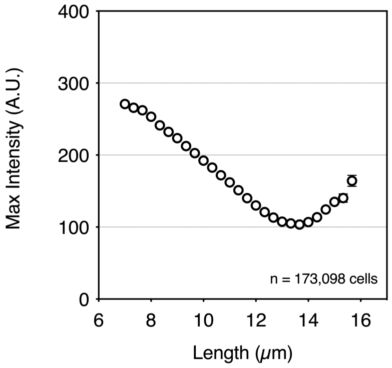

### mik1-mNG

PN5825/PN5887
SPBC660.14
mitotic inhibitor kinase

Widefield: 19/04/18

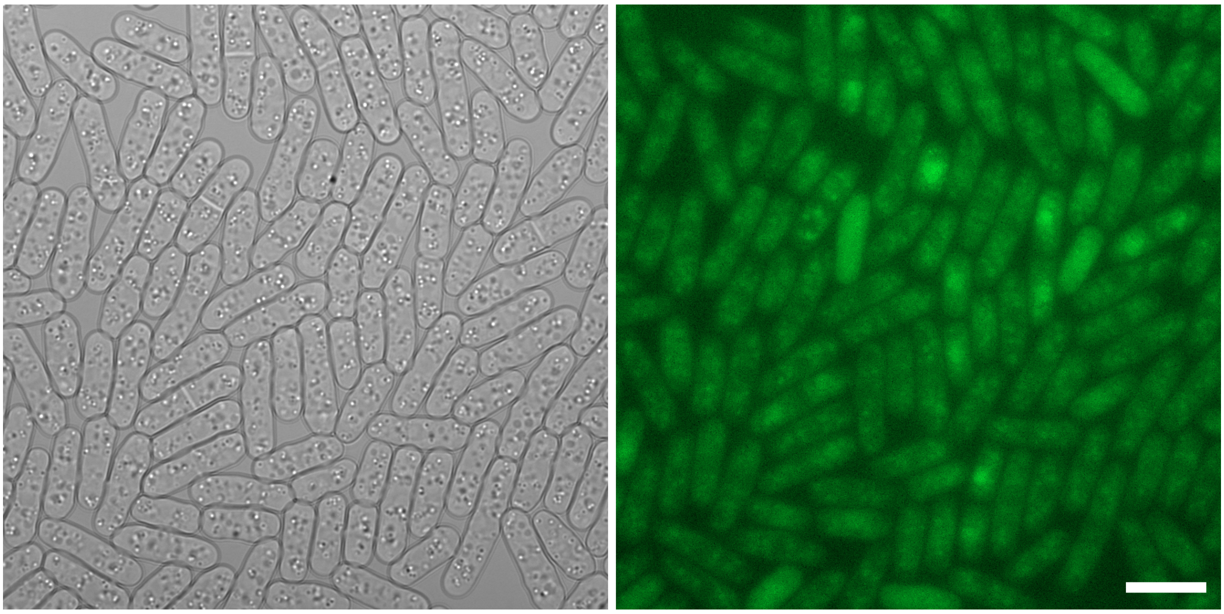

Widefield: 19/04/18

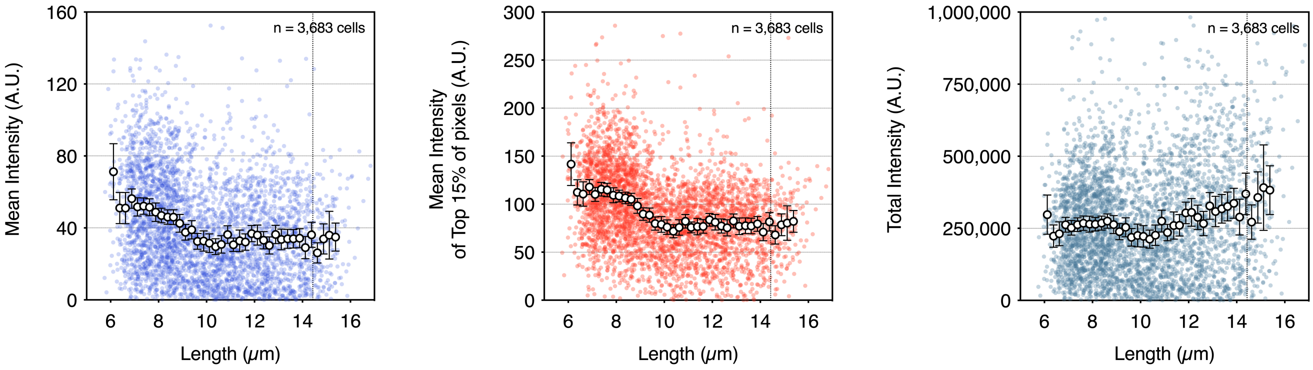


Widefield: 09/08/18

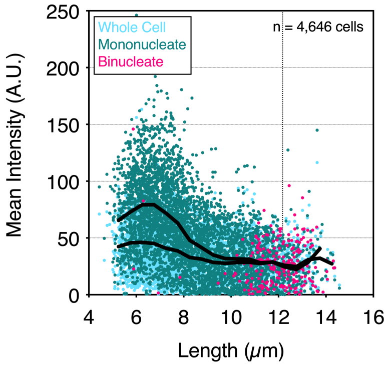


Imagestream: 10/10/18

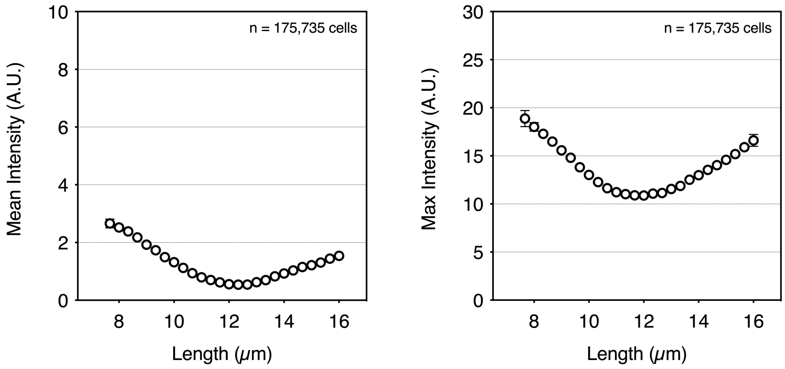

### puc1-mNG

PN5815
SPBC19F5.01c
cyclin Puc1

Widefield: 27/04/18

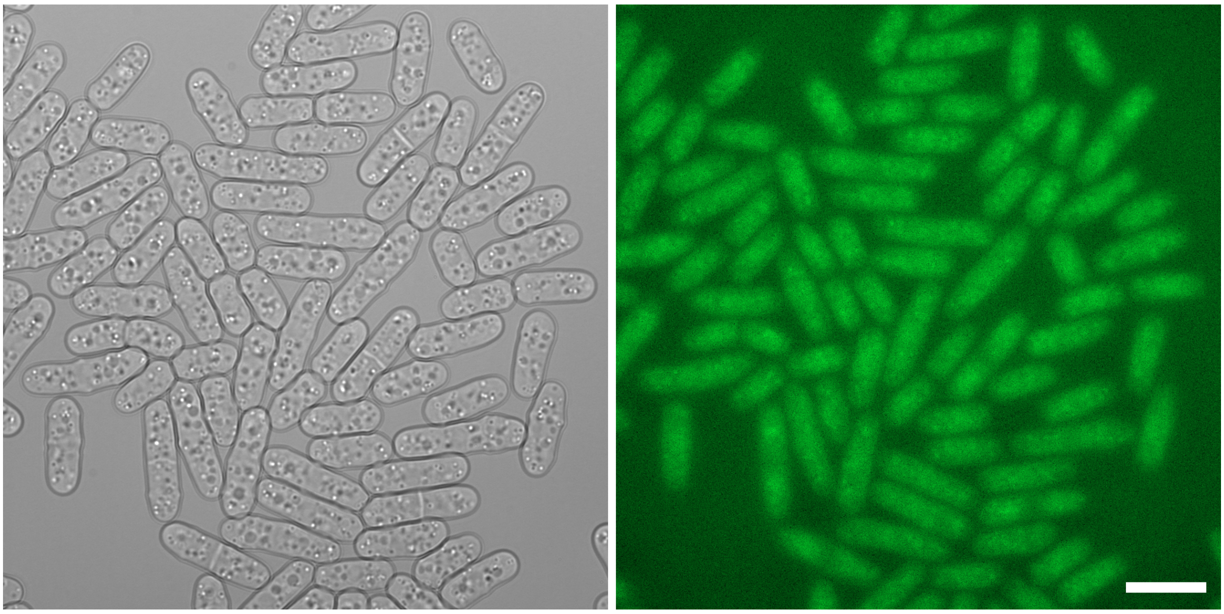

Widefield: 27/04/18

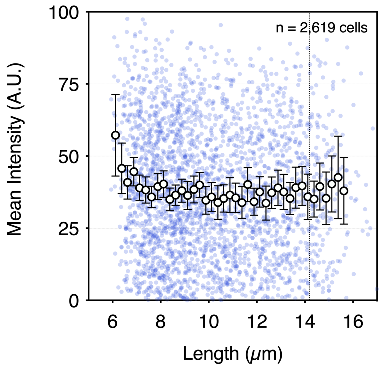


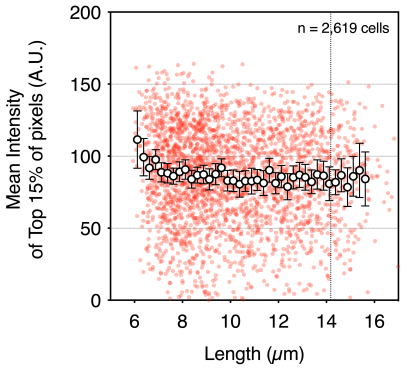


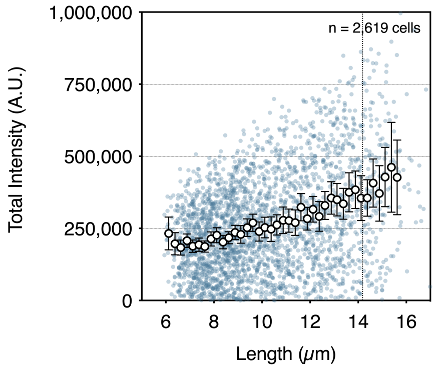


Imagestream: 10/10/18

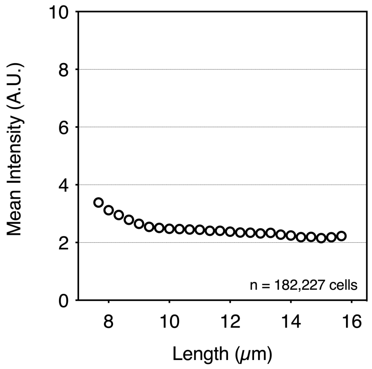


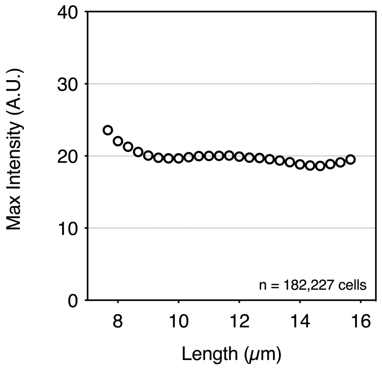

### rum1-mNG

PN5821/SC333
SPBC32F.09
CDK inhibitor Rum1

Widefield: 27/04/18

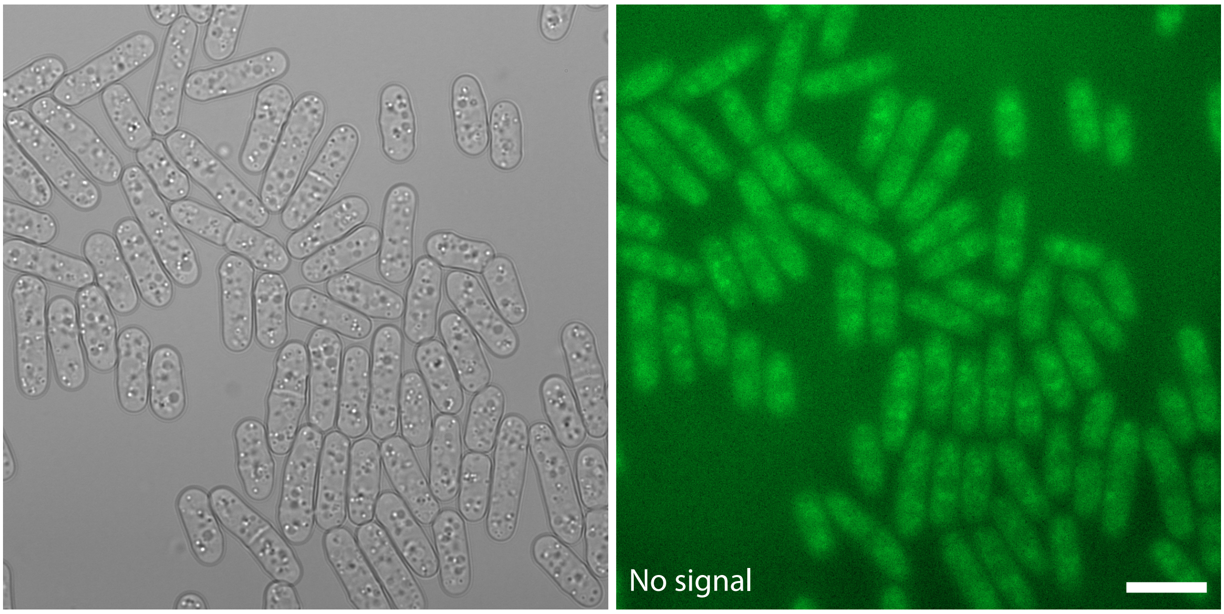


Widefield: 27/04/18

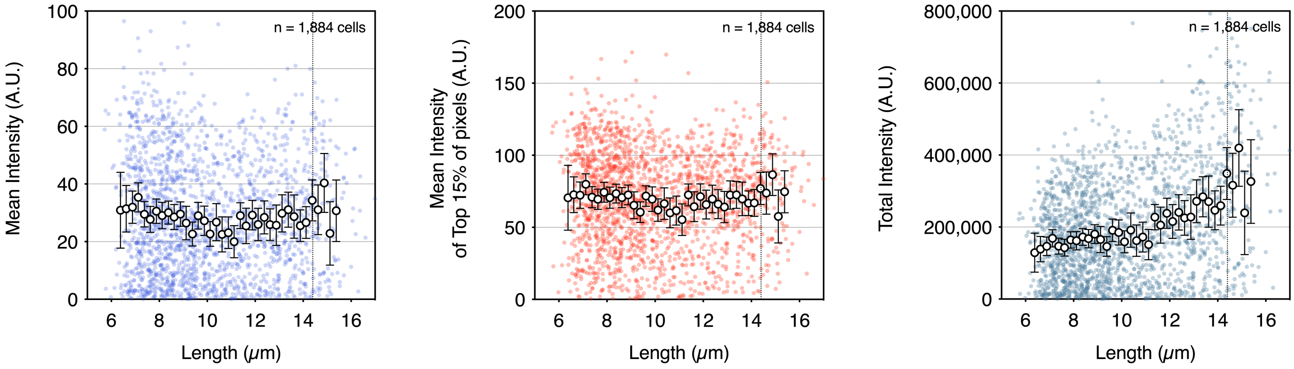


Imagestream: 09/08/18

Imagestream: 10/10/18

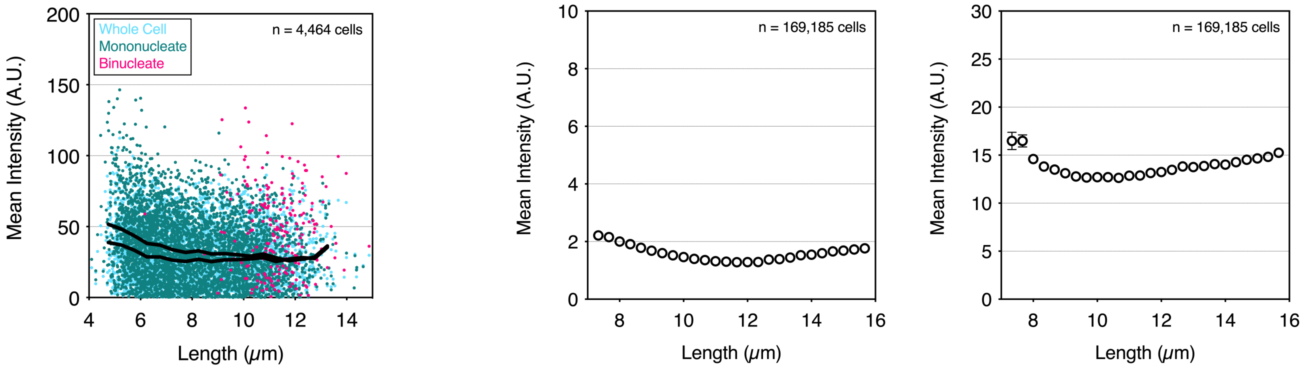

### srw1-mNG

PN5819
SPAC144.13c
substrate-specific mitotic G1 APC coactivator Srw1/Ste9

Widefield: 27/04/18

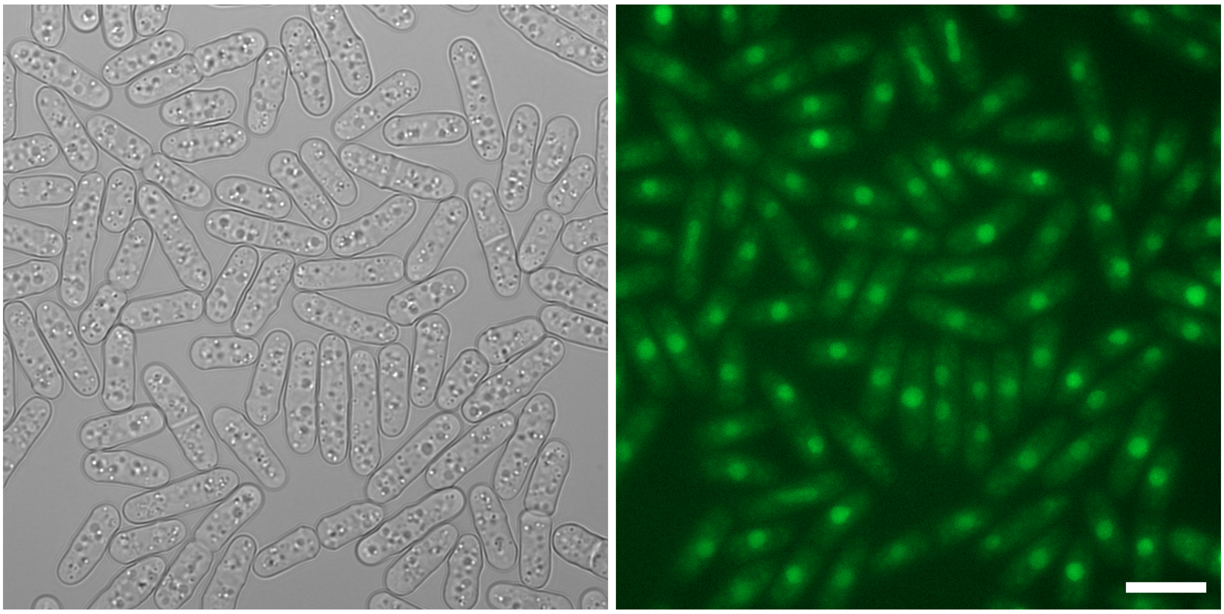

Widefield: 27/04/18

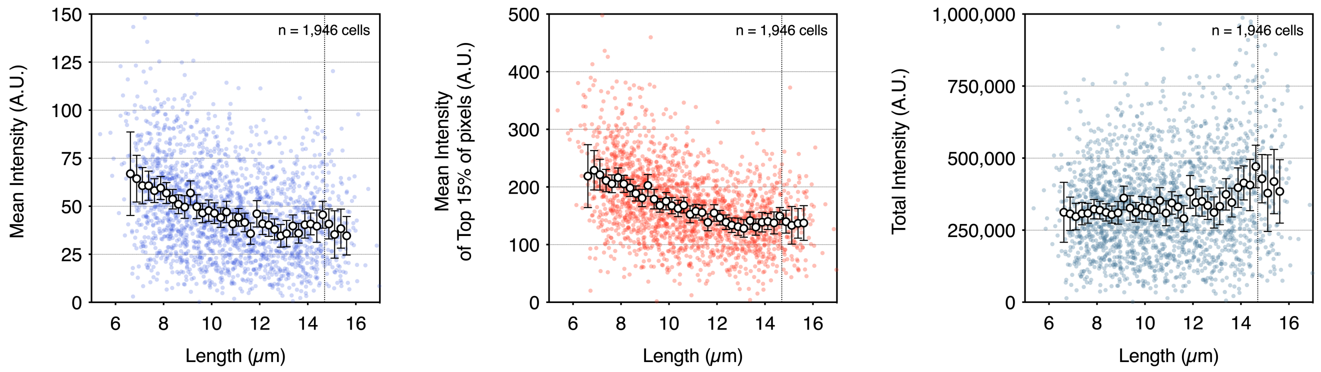


Imagestream: 10/10/18

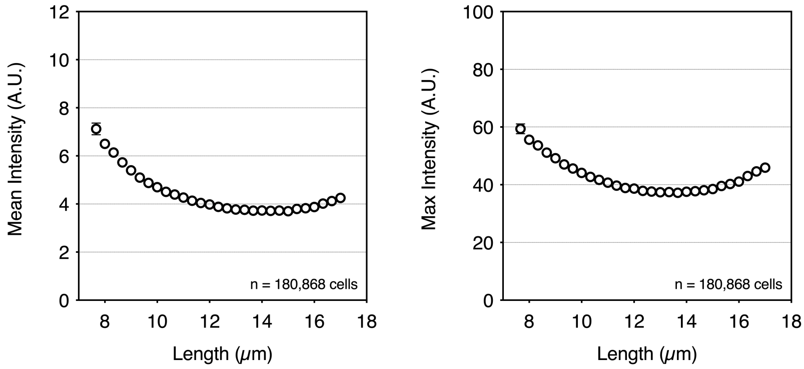

### cdr2-mNG

PN5878
SPAC57A10.02
serine/threonine protein kinase Cdr2

Widefield: 21/12/18

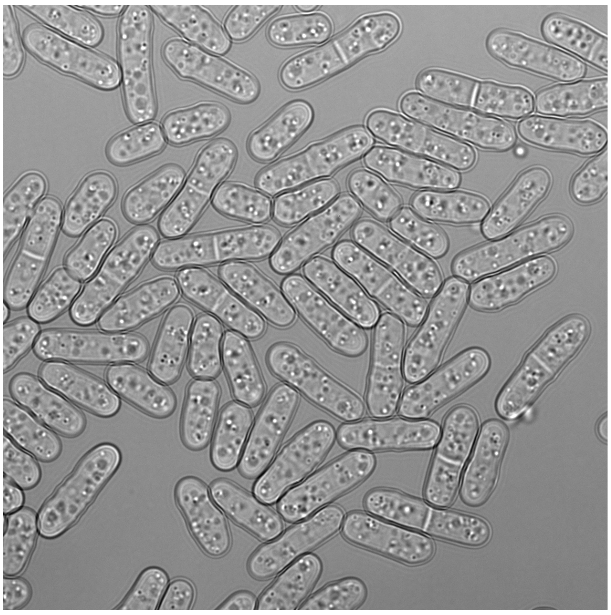

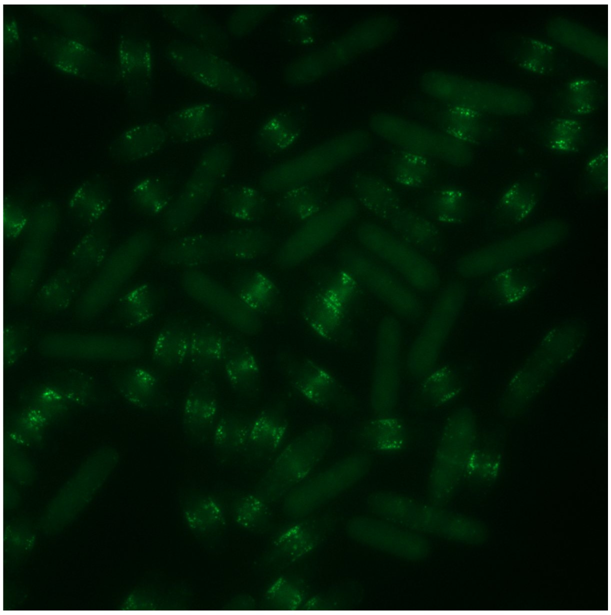

Widefield: 21/12/18

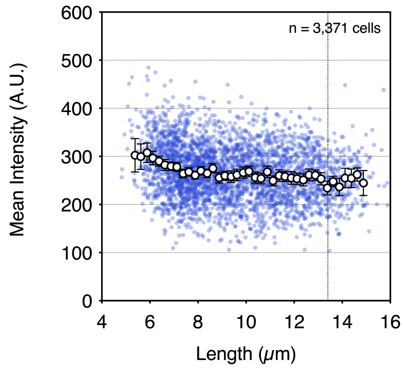


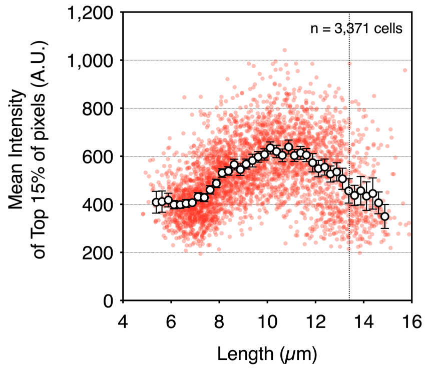


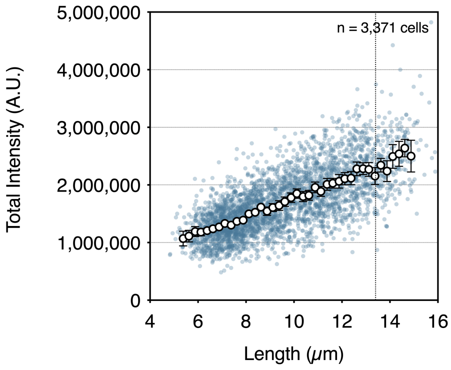


Imagestream: 12/10/18

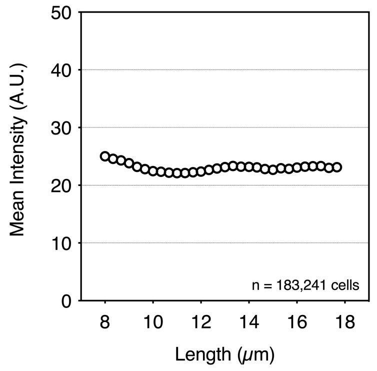


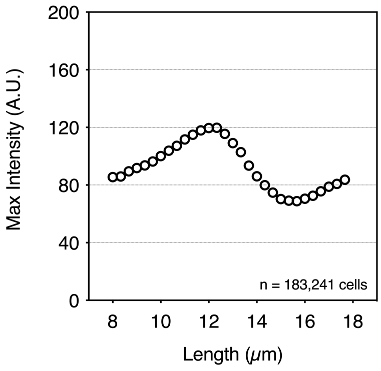

### plo1-mNG

PN5877
SP23C11.16
Polo kinase Plo1

Widefield: 20/04/18

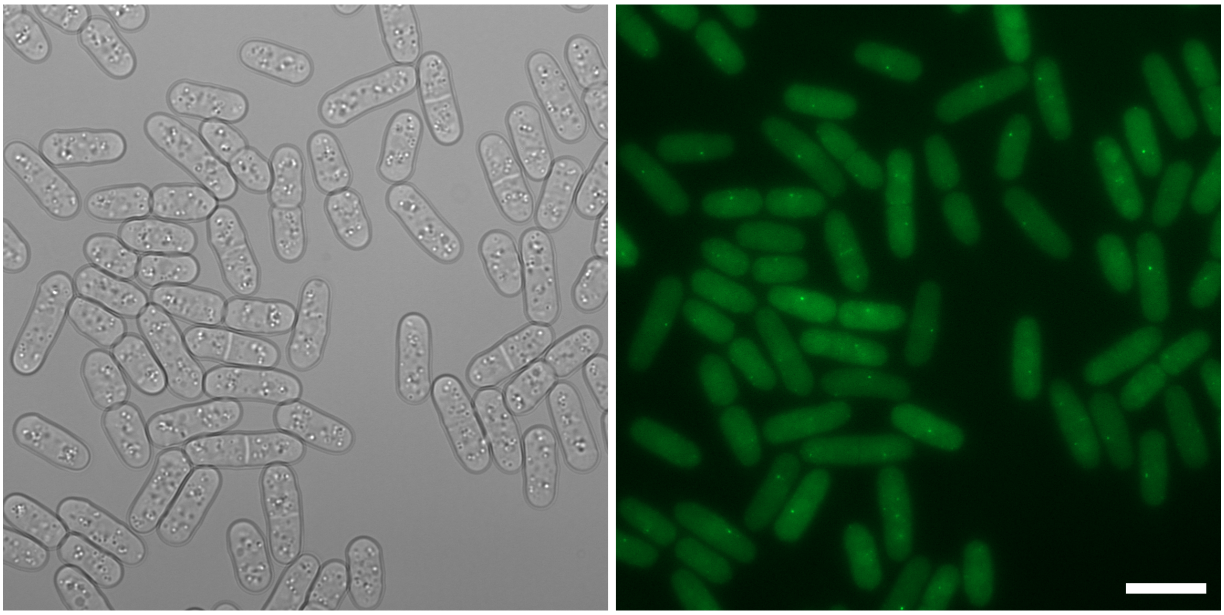

Widefield: 20/04/18

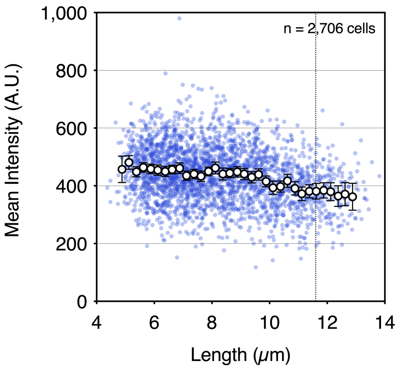


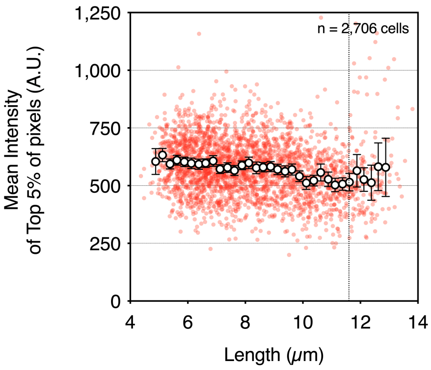


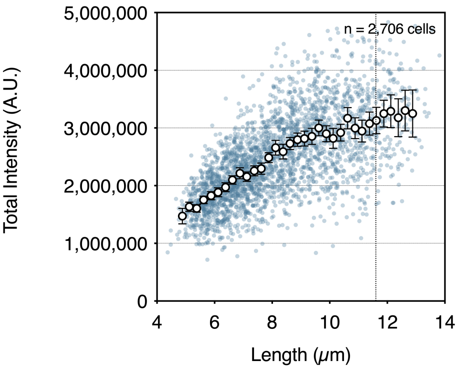


Imagestream: 12/10/18

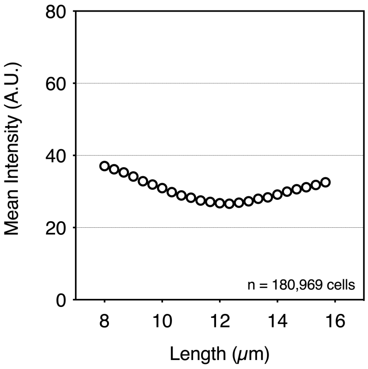


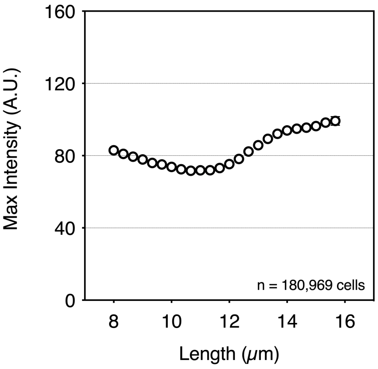

### prp19-mNG

PN5879
SPAC29A4.08c
Prp19 complex subunit, ubiquitin-protein ligase E4 Prp19

Widefield: 20/04/18

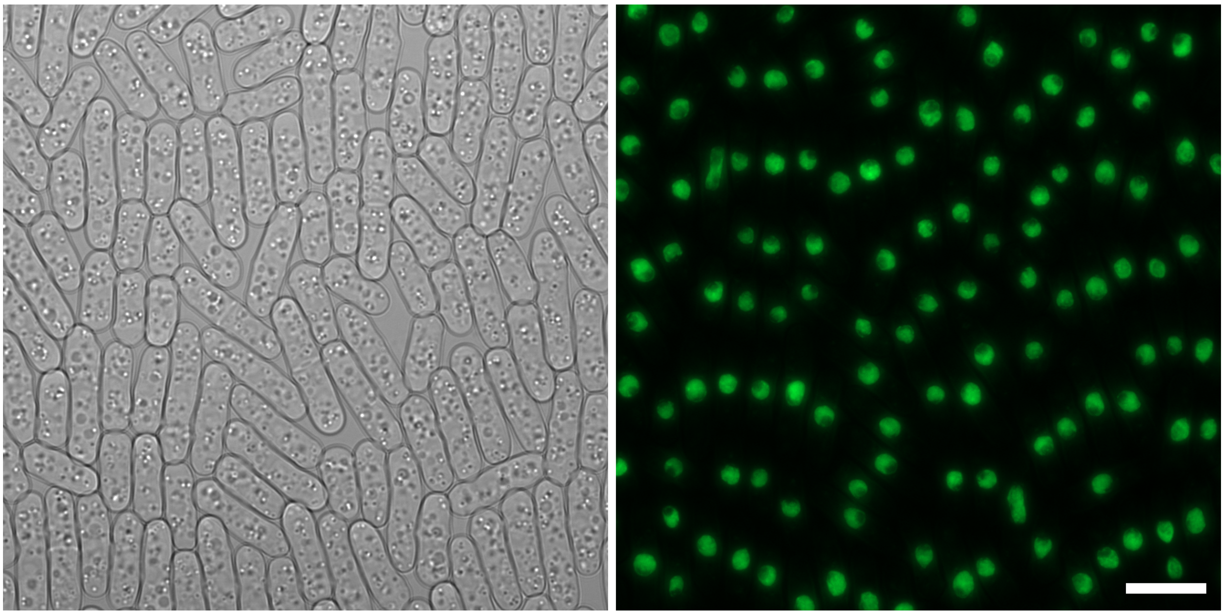

Widefield: 20/04/18

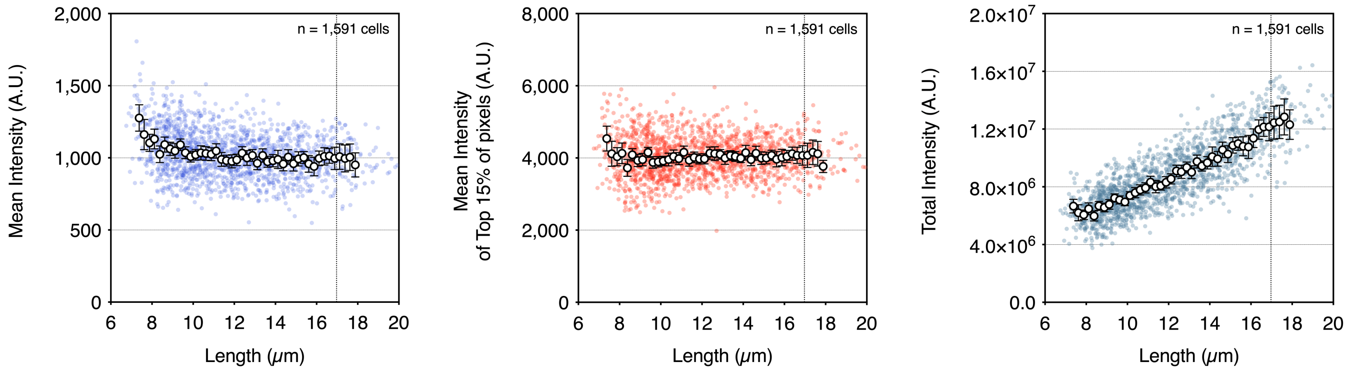


Imagestream: 12/10/18

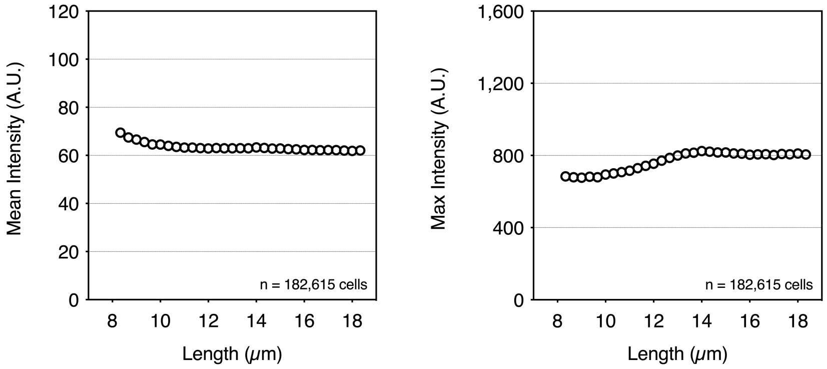

### pyp3-mNG

PN5816
SPAC11E3.09
protein-tyrosine phosphatase Pyp3

Widefield: 27/04/18

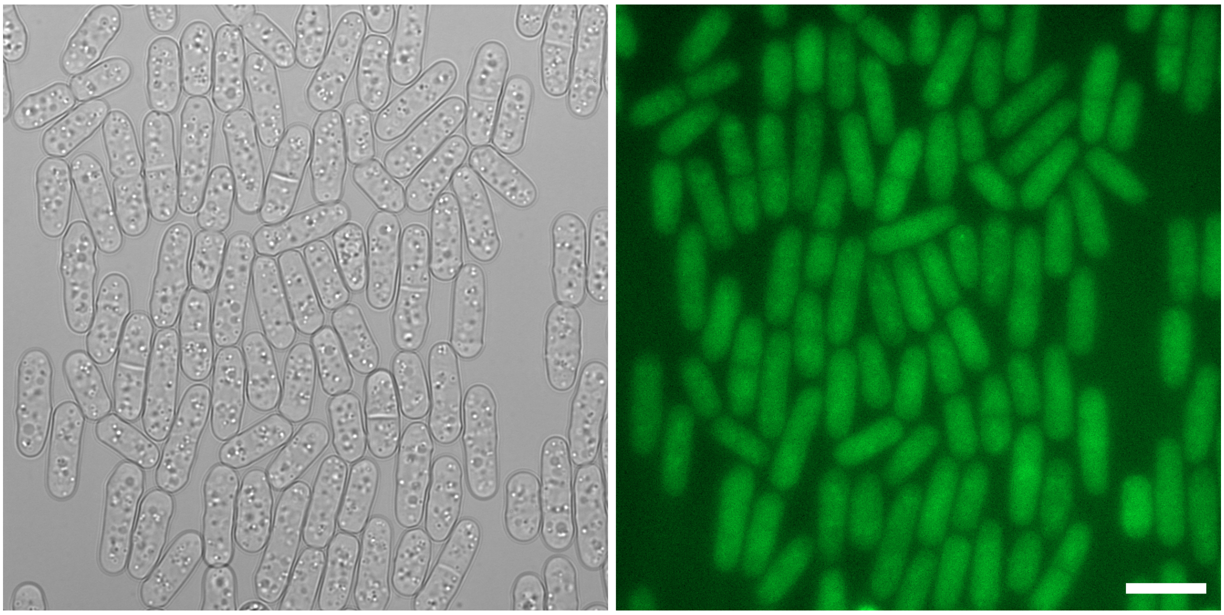

Widefield: 27/04/18

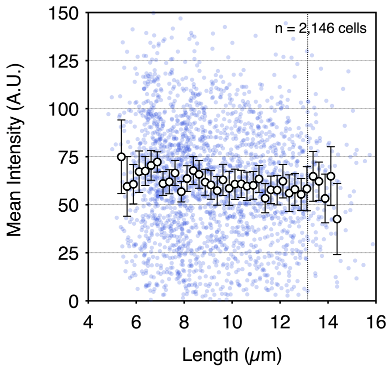


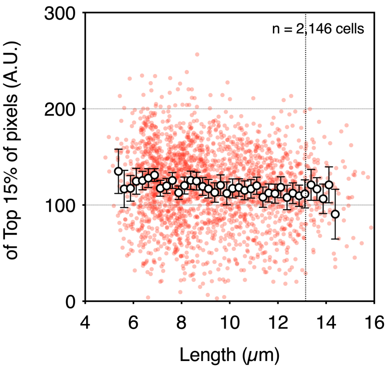


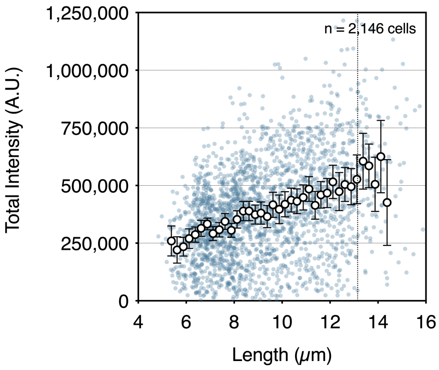


Imagestream: 10/10/18

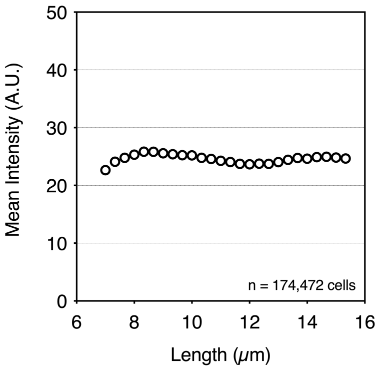


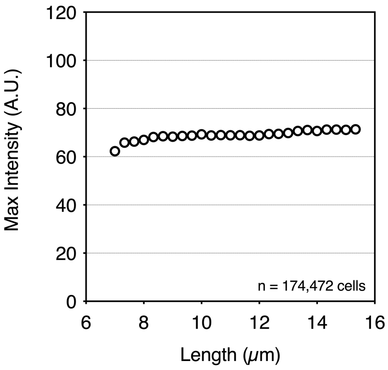

### slm9-mNG

PN5875
SPBC15D4.03
Histone H3.3 H4 chaperone, hira family Slm9

Widefield: 20/04/18

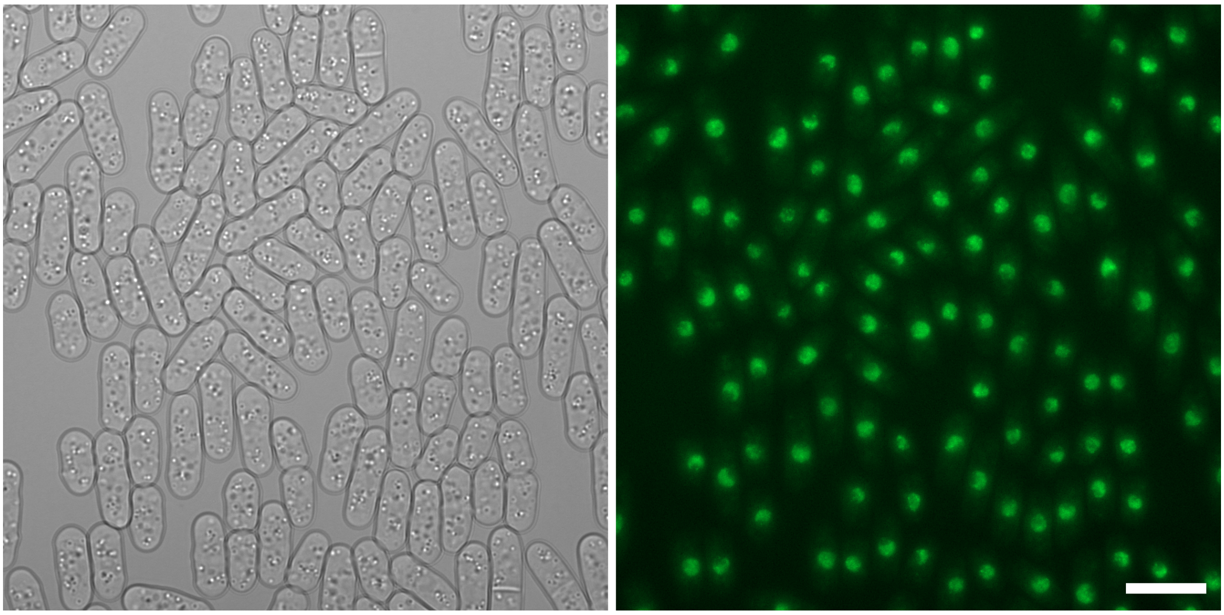

Widefield: 20/04/18

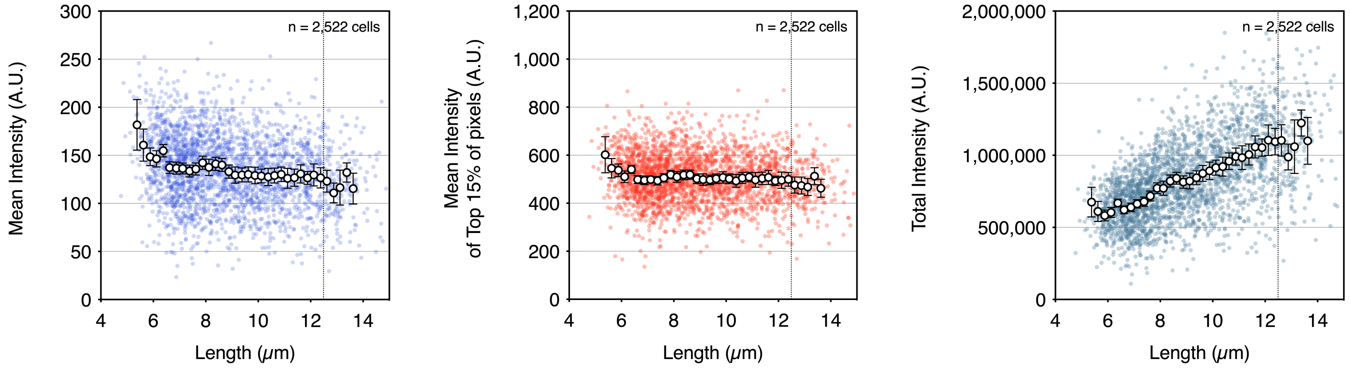


Imagestream: 12/10/18

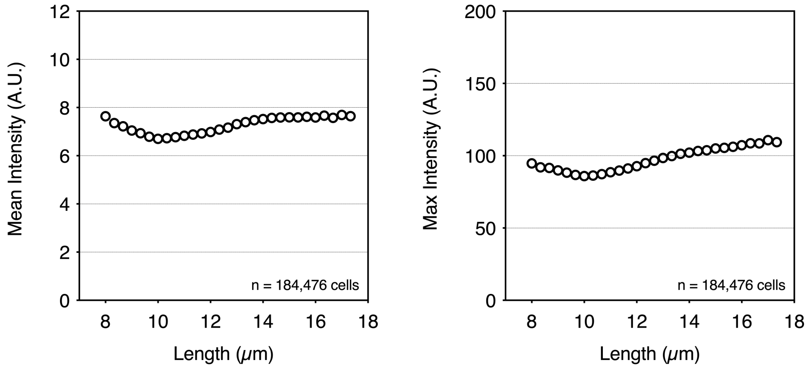

### snf21-mNG

PN5880
SPAC1250.01
RSC-type complex ATP-dependent DNA helicase Snf21

Widefield: 19/04/18

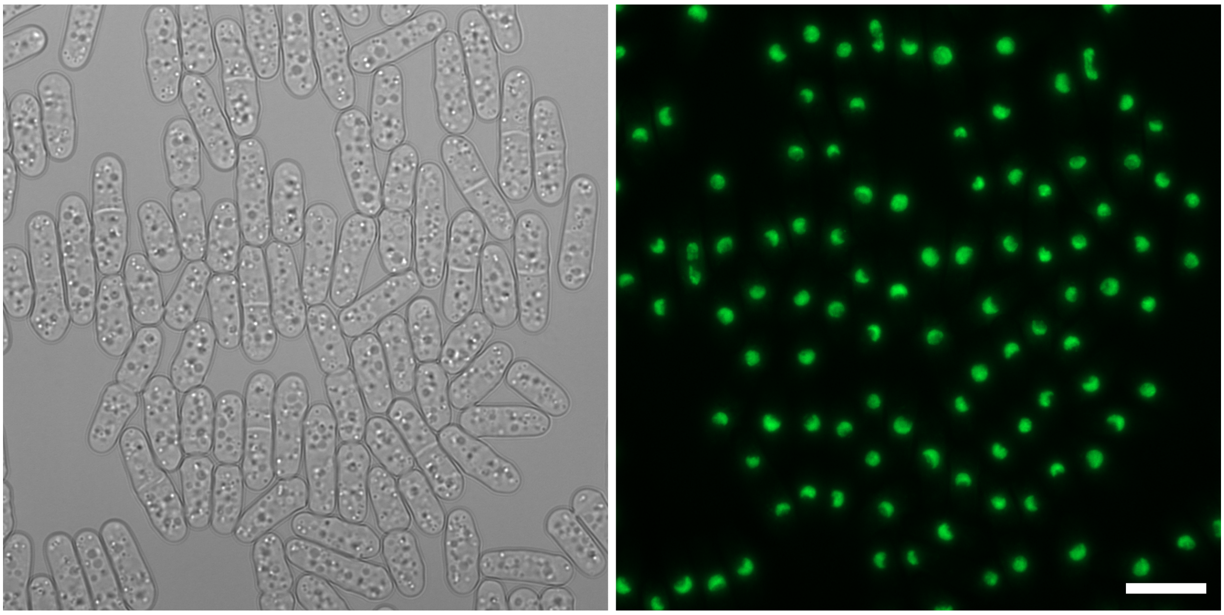

Widefield: 19/04/18

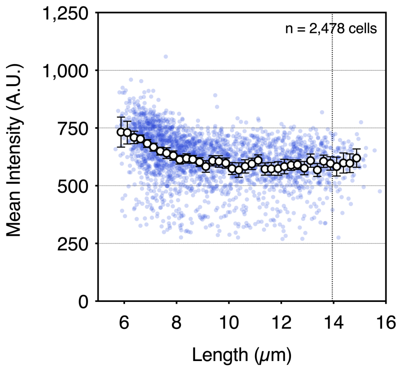


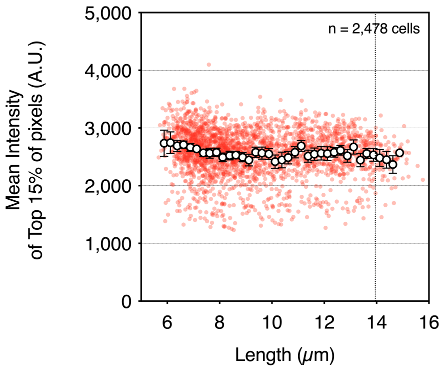


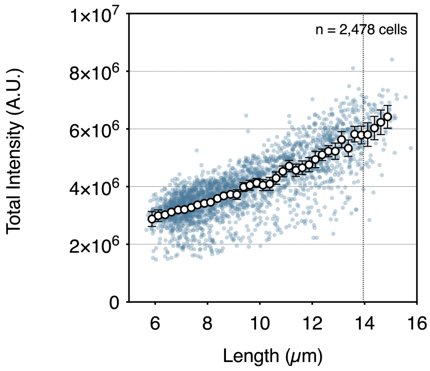


Imagestream: 12/10/18

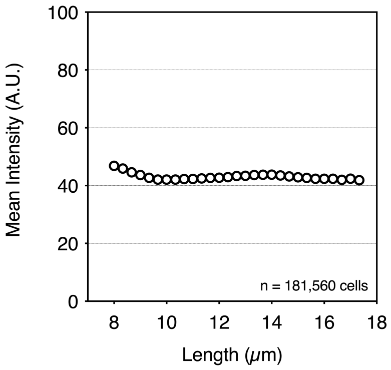


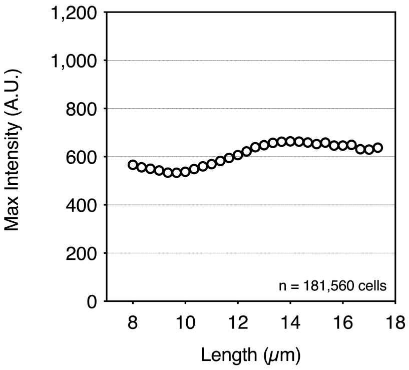

### snf22-mNG

PN5881
SPCC1620.14c
ATP dependent DNA helicase Snf22

Widefield: 19/04/18

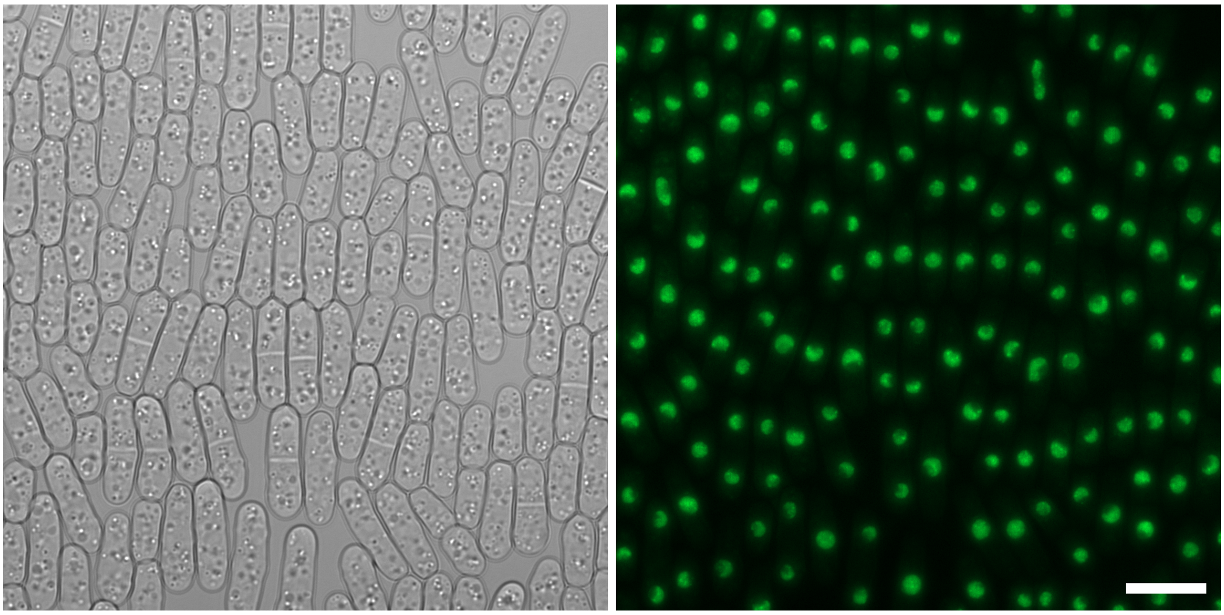

Widefield: 19/04/18

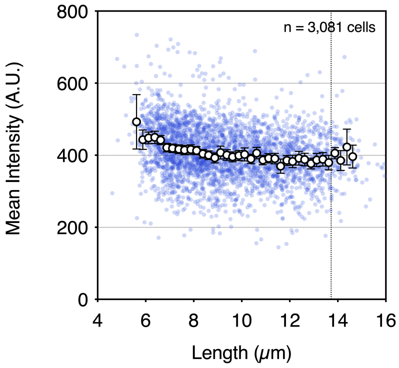


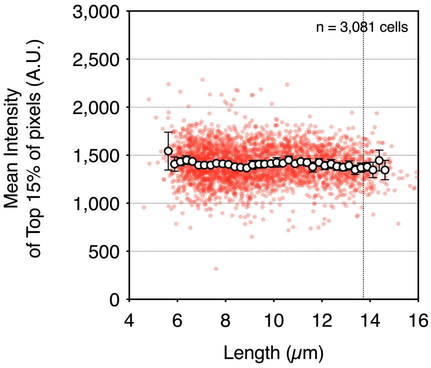


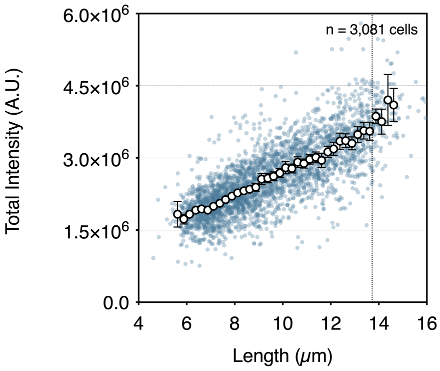


Imagestream: 12/10/18

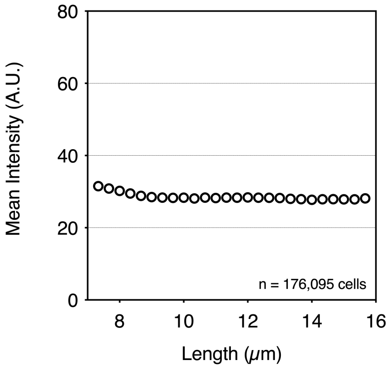


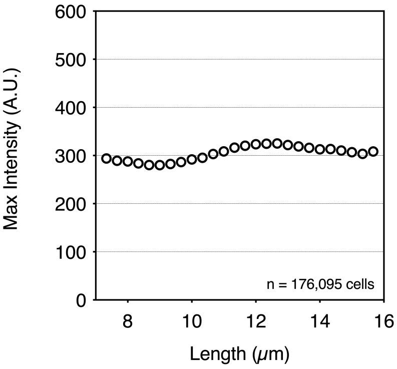

### tif45-mNG

PN5876
SPAC16E8.15
translation initiation factor eIF4E, 4F complex subunit

Widefield: 19/04/18

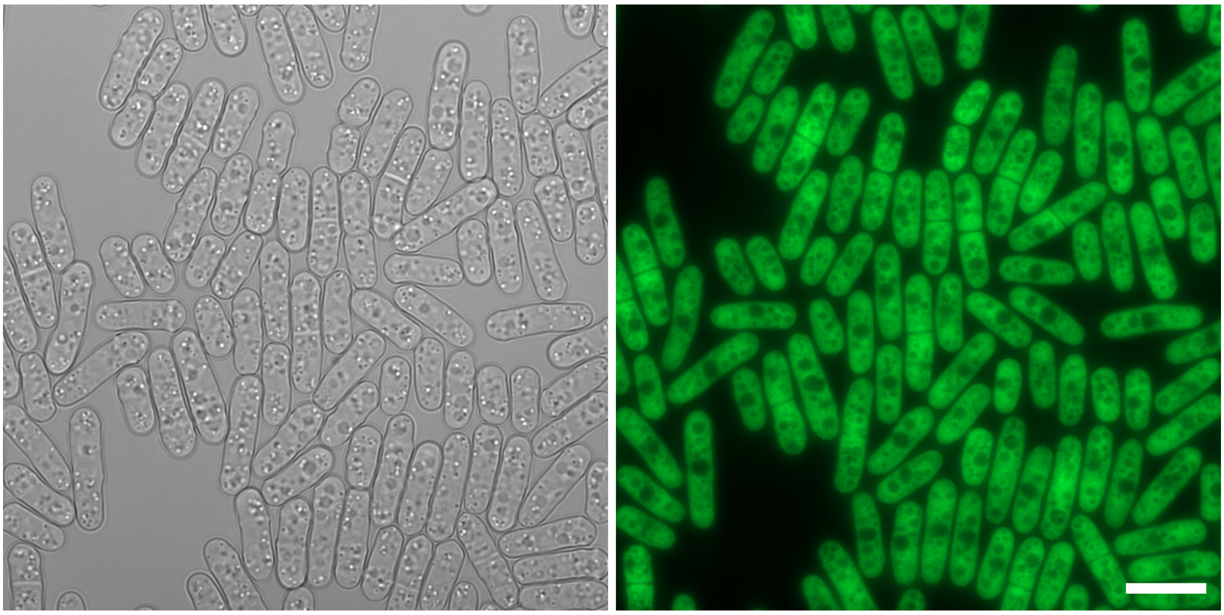

Widefield: 19/04/18

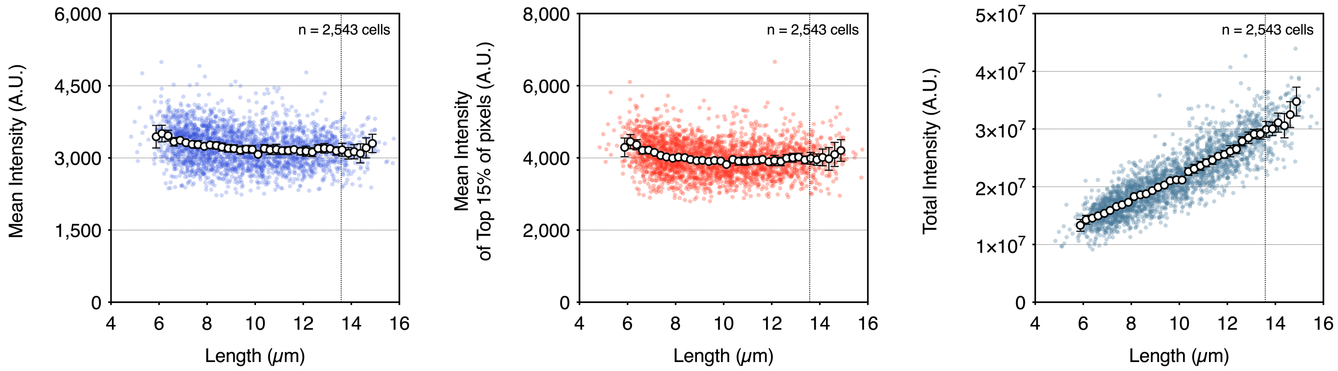


Imagestream: 13/12/18

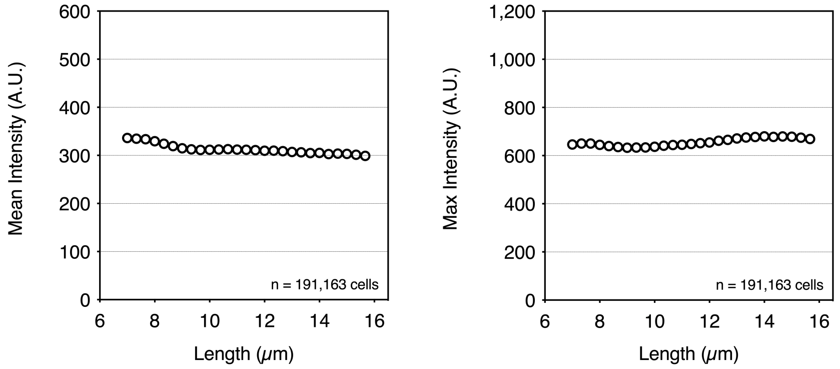
